# Revealing the diversity of commensal corynebacteria from a single human skin site

**DOI:** 10.1101/2024.11.28.625817

**Authors:** Reyme Herman, Sean Meaden, Michelle Rudden, Robert Cornmell, Holly N. Wilkinson, Matthew J. Hardman, Anthony J. Wilkinson, Barry Murphy, Gavin H. Thomas

**Affiliations:** Department of Biology, University of York, Wentworth Way, York, YO10 5DD, UK; School of Life Sciences, University of Hull, Hull, HU6 7RX, UK; York Structural Biology Laboratory, Department of Chemistry, University of York, Wentworth Way, York, YO10 5DD, UK; Unilever Research & Development, Port Sunlight Laboratory, Quarry Road East, Bebington, Wirral, Merseyside, CH63 3JW, UK

## Abstract

Our understanding of the skin microbiome has dramatically improved since the pioneering studies and the improvements in sequencing technologies. Species of the genus *Corynebacterium* are known to form a major part of the human skin microbiome but most detailed studies have focussed on other similarly prevalent genera like *Staphylococcus* and *Cutibacteria*. Prior to this study, there were few complete genomes for skin commensals of the genus *Corynebacterium*, with only 9 complete genomes available for the most commonly identified species *C. tuberculostearicum*. In this study we explored the genus *Corynebacterium* from a single body site by swabbing the axilla/underarm of 4 individuals and using a selective media to enrich for corynebacteria. We then generated whole genome sequencing data of these corynebacteria enriched isolates using long-read sequencing and subsequent bioinformatics analysis to reveal an unparalleled diversity of this genus from a single skin site. Through this approach, we obtained the closed genomes of 215 isolates, 154 derived from a single individual. With this genetic information, we were able to identify 7 different species including species previously not associated to the skin and two novel species provisionally named *C. axilliensis* and *C. jamesii*. We used pangenome analysis on 30 genetically distinct isolates spanning the 7 species to identify putative metabolic differences, antimicrobial resistance profiles, novel biosynthetic gene clusters (BGCs), prophages and phage defence systems. Our culture-based Nanopore sequencing approach has dramatically improved our overall knowledge of skin corynebacteria, and uniquely here also providing in-depth analysis from a single skin site, revealing a multitude of differences between the isolates. Here we not only improved our knowledge of axillary corynebacteria but also greatly expanded the publicly available number of cutaneous corynebacterial genomes complementing recent studies seeking to understand the diversity of skin corynebacteria.

## Introduction

The exploration of the human skin microbiome allows deeper understanding of how commensal microbes and the host interact with each other to maintain skin health. Human skin, generally a nutrient limited environment, hosts a variety of commensal microbes spanning multiple genera. Skin sites can be characterised as dry, moist or oily (sebaceous), depending on the architecture of the skin and abundance of the skin associated glands, namely the eccrine, sebaceous or apocrine glands. These different environmental conditions can play a major role in determining microbial diversity [1]. While *Cutibacterium* and *Staphylococcus* are the two major commensal genera identified, other skin associated genera like *Corynebacterium* are also highly abundant [2, 3]. In a study by Grice *et al*., out of over 112,000 16S rRNA sequences derived from 20 different skin sites, corynebacteria comprises of 22.8% of the sequences second only to cutibacteria (23%) [4]. Some corynebacterial species have been found to be lipophilic and hence require lipids derived from the sebaceous gland or the stratum corneum [5, 6]. Commonly identified commensal skin corynebacteria like *C. jeikeium* [7], *C. amycolatum* [8] and *C. kroppenstedtii* [9] have been suggested to be opportunistic pathogens. *C. accolens* [10] and *C. tuberculostearicum* [11] have been shown to interact with human immune system causing inflammation. *C. accolens* has been shown to also inhibit *Streptococcus pneumoniae* [12, 13] while *C. striatum* is proposed to be able to shift *Staphylococcus aureus* from the virulent to a commensal state [14], thus demonstrating potential protection capabilities. Together, these studies highlight the importance of understanding the healthy human skin microbiome in our efforts to combat pathogenic species.

Human malodour is microbially generated from a metabolite secreted from the apocrine gland. Although a monophyletic group of staphylococci was attributed to this phenomenon [15], some studies have proposed a role for axillary *Corynebacterium* in the human malodour formation due to the correlation of abundance and odour profiles [16] and the presence of enzymatic components for the biotransformation of the human derived precursor [17–19]. Further, a genus level study analysing the diversity of axillary commensals of 9 individuals found that 59.5% of sequences were binned to the genus *Corynebacterium* [20]. Our understanding of the role of cutaneous corynebacteria in human malodour remains unclear. Attaining genomic information of these axillary corynebacteria could reveal if they do possess the relevant genes for malodour production.

Despite the abundance of corynebacteria on the human skin, the amount of genetic information on cutaneous corynebacterial species is surprisingly lacking which implies a lack of understanding of its true diversity within the collective skin microbiome. In the case of *C. tuberculostearicum*, a species widely found on various skin sites, only 7 complete genomes have been made publicly available on GenBank prior to this study. In comparison, there are 221 available complete genomes of another skin commensal, *Staphylococcus epidermidis*. Two recent comparative genomics studies have sought to address the lack of genetic information through a culture-based approach. The first focusses on skin isolates of the *C. tuberculostearicum* species complex which is made up of multiple closely related species including *C. kefirresidentii* [21]. In addition, the study reports the identification of a novel species of corynebacteria that was predominantly found on the toe web and toenails of multiple individuals. A second comparative genomics study on human skin corynebacteria [22]revealed the diversity of corynebacteria on the human facial skin. This study too identified two novel species of corynebacteria.

Here, we sought to elucidate the diversity of corynebacteria within the human underarm (axilla). This is a moist skin site with metabolites derived from the three skin associated glands and the skin itself, making it a relatively nutrient rich skin environment when compared with most of the other skin sites. In this study we took a culture-based approach followed by sequencing, consistent with recent comparative genomics studies. However, these studies adopted similar approaches by initially performing 16S rRNA (V1-V3 region) sequencing and identifying representatives within amplicon sequence variants (ASVs) to whole genome sequence. We instead adopted a direct-to long-read whole genome sequencing strategy by omitting the 16S rRNA gene sequencing which could introduce sampling bias. In this study, we generated complete genomes of over 200 corynebacteria isolates derived from axillary swabs of 4 individuals using the Oxford Nanopore sequencing technology. Our sequencing revealed 7 distinct species of axillary corynebacteria, two of which are proposed to be novel species of corynebacteria. We also further clarified the diversity within the *C. tuberculostearicum* species complex to complement recent studies. Downstream bioinformatics analysis revealed the genetic and functional diversity within our collection of axillary corynebacteria which would help reveal the roles of this genus of skin commensal bacteria.

## Results

### Assessing the diversity of axillary corynebacteria isolates

Axillary swabs from 4 human volunteers (A1, A3, A4 and A5) were collected and then enriched for corynebacteria on selective agar. To assess the ability of our approach to capture the diversity of corynebacteria within our isolates, we generated complete genome and plasmid sequences for 150 corynebacteria isolates from a single individual A1 and complemented the study with small subsets of isolates from the three other volunteers; 22 isolates from A3, 26 isolates from A4 and 13 isolates from A5 (**Fig. 1A**).

**Figure 1.**
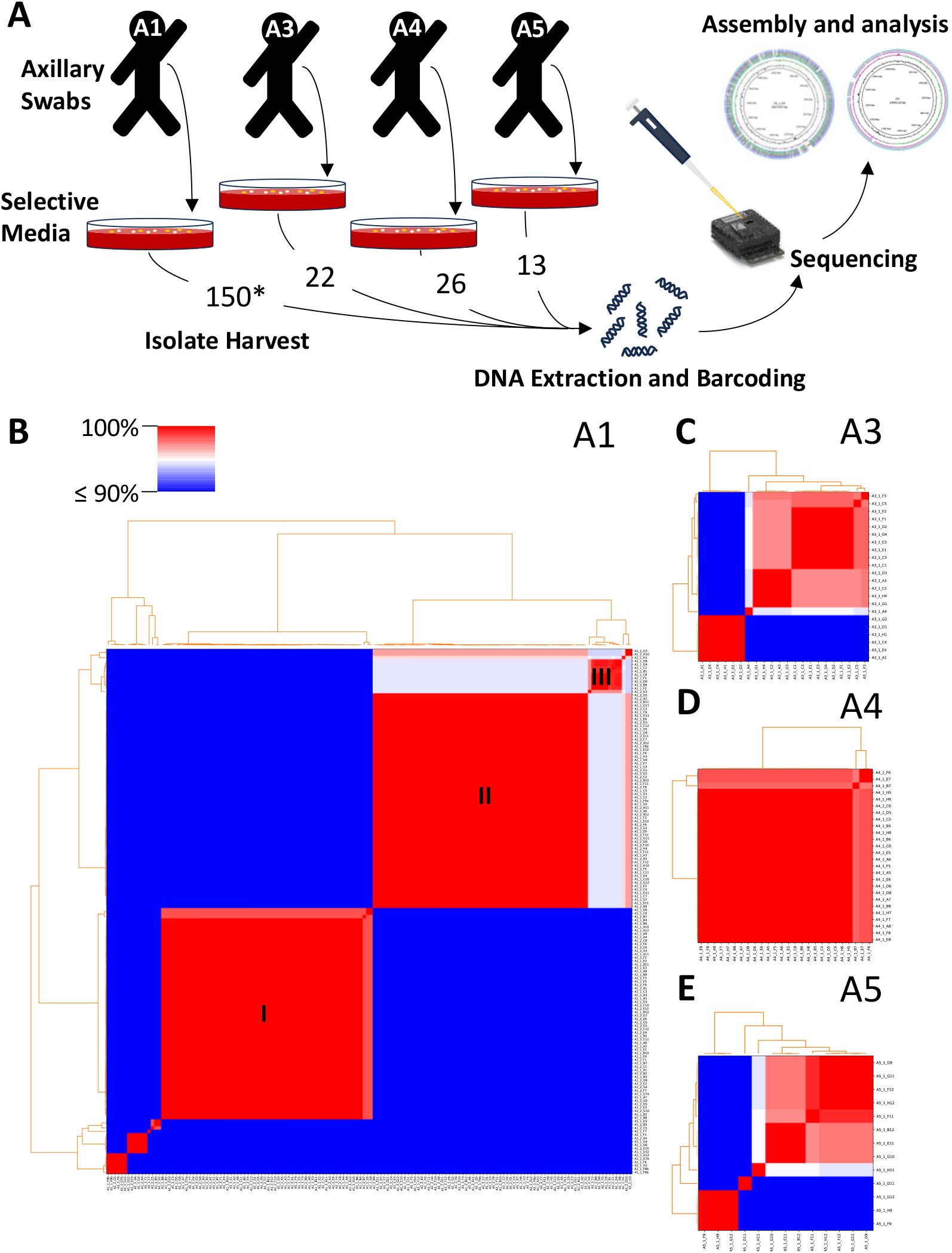
Capturing the diversity of axillary corynebacteria. (A) Axillary *Corynebacterium* isolates were harvested from swabs from 4 volunteers. Long-read whole genome sequencing and downstream analysis was performed on a 150 isolates of volunteer A1, 22 from volunteer A3, 26 from volunteer A4 and 13 from volunteer A5. (B-E) Pairwise average nucleotide identities (ANI) were calculated with pyANI and used to assess the genetic diversity of the isolates. ANI values were placed onto a coloured pairwise matrix according to the colour scale depicted in (B) for isolates derived from each volunteer. Three major groups of isolates were identified with I, II and III from the isolates from A1.

In order to initially assess the diversity of the corynebacterial isolates from all 4 volunteers, [23]we compared pairwise average nucleotide identities (ANI) between assembled genomes from each individual (**Fig. 1B-E**) and collectively (**Supplementary Fig. 1**). The pairwise percentage identity plots for the 150 Isolates from A1 revealed that the majority of isolates recovered by culture belonged to two discrete species (labelled as I and II) (**Fig. 1B**). However, the depth of sampling also revealed a third cluster (group III) and number of other smaller groups, some only containing one isolate, suggesting the approach had recovered significant diversity from this single body site from a single individual. For individuals A3 and A5 there was a similar pattern of a number of different clusters identified, despite there being a much smaller total number of genomes (**Fig. 1C** and **Fig. 1E**). Surprisingly in individual A4 we found that almost all of the 26 isolates were very similar to each other, suggesting that one strain/species dominates this community (**Fig. 1D**).

We identified representative genomes from groups of genetically similar isolates, with a 95% ANI used to distinguish between different species and an ANI of 99.5% to capture strain diversity (**Supplementary Fig. 2**). The isolates bin to 7 different species and the secondary clustering further identified 30 distinct isolates which we focussed further analysis on. To initially identify the species of these 30 isolates, their 16S rRNA (V1-V3 region) sequences were analysed using BLASTn and collectively aligned with the closest hits of each isolate (**Supplementary Fig. 3A**). Using this analysis, we observe 5 distinct clades with the largest clade containing 24 of our isolates which were found to be ≥ 99.8% identical (**Supplementary**

**Figure 2.**
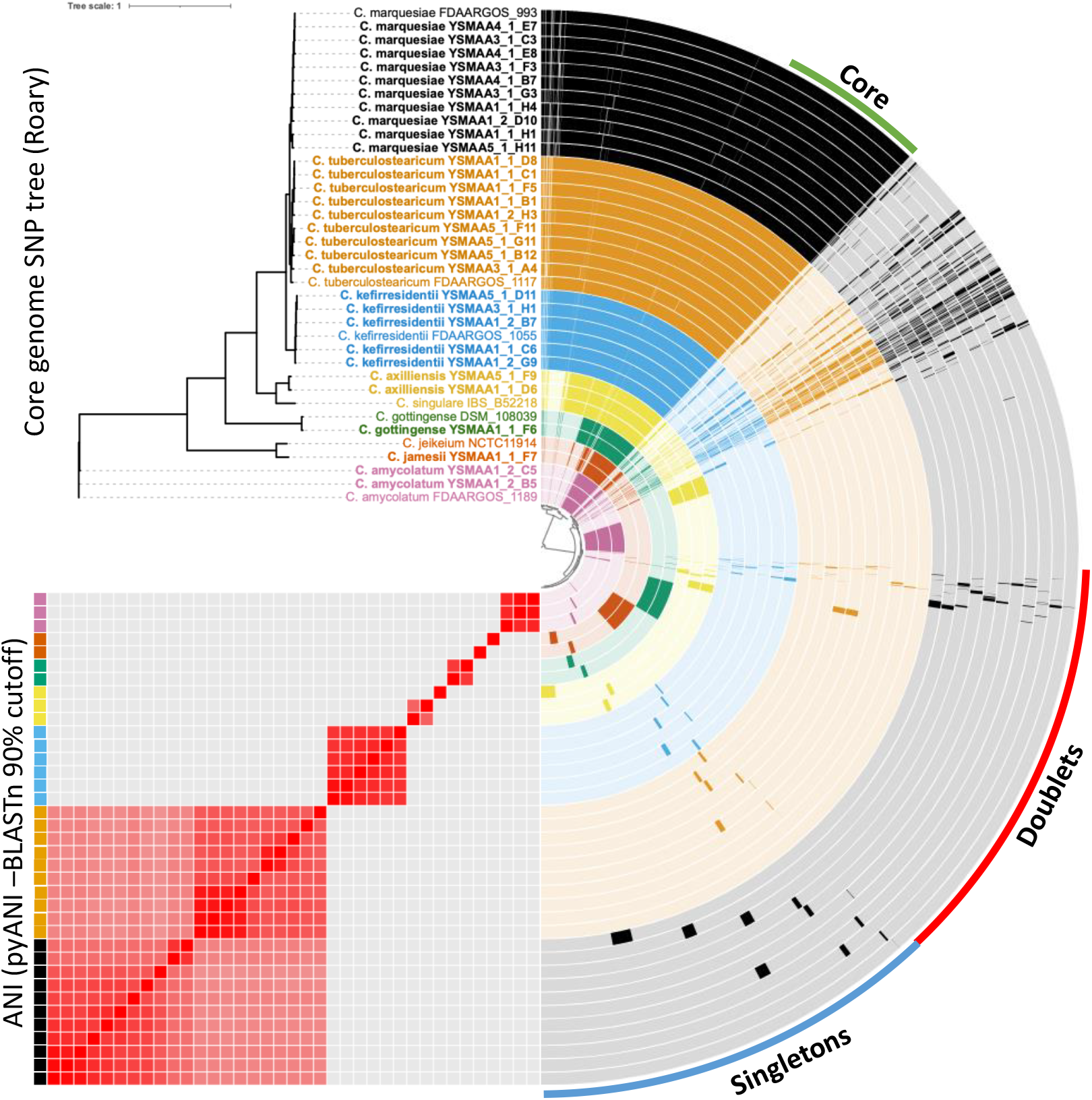
The pangenome of the axillary corynebacteria isolates (bold) with the closest RefSeq representatives for each species calculated and visualised using anvi’o. Gene clusters found in either one (Singletons) or two (Doublets) were identified. A core genome was calculated using Roary with an 80% BLASTp cutoff and visualised on a tree generated using PhyML and visualised on iTOL. An ANI comparison using BLASTn on pyANI with a 90% cutoff was generated and visualised on anvi’o. Isolates of the 7 clusters of species were colour coded.

**Figure 3.**
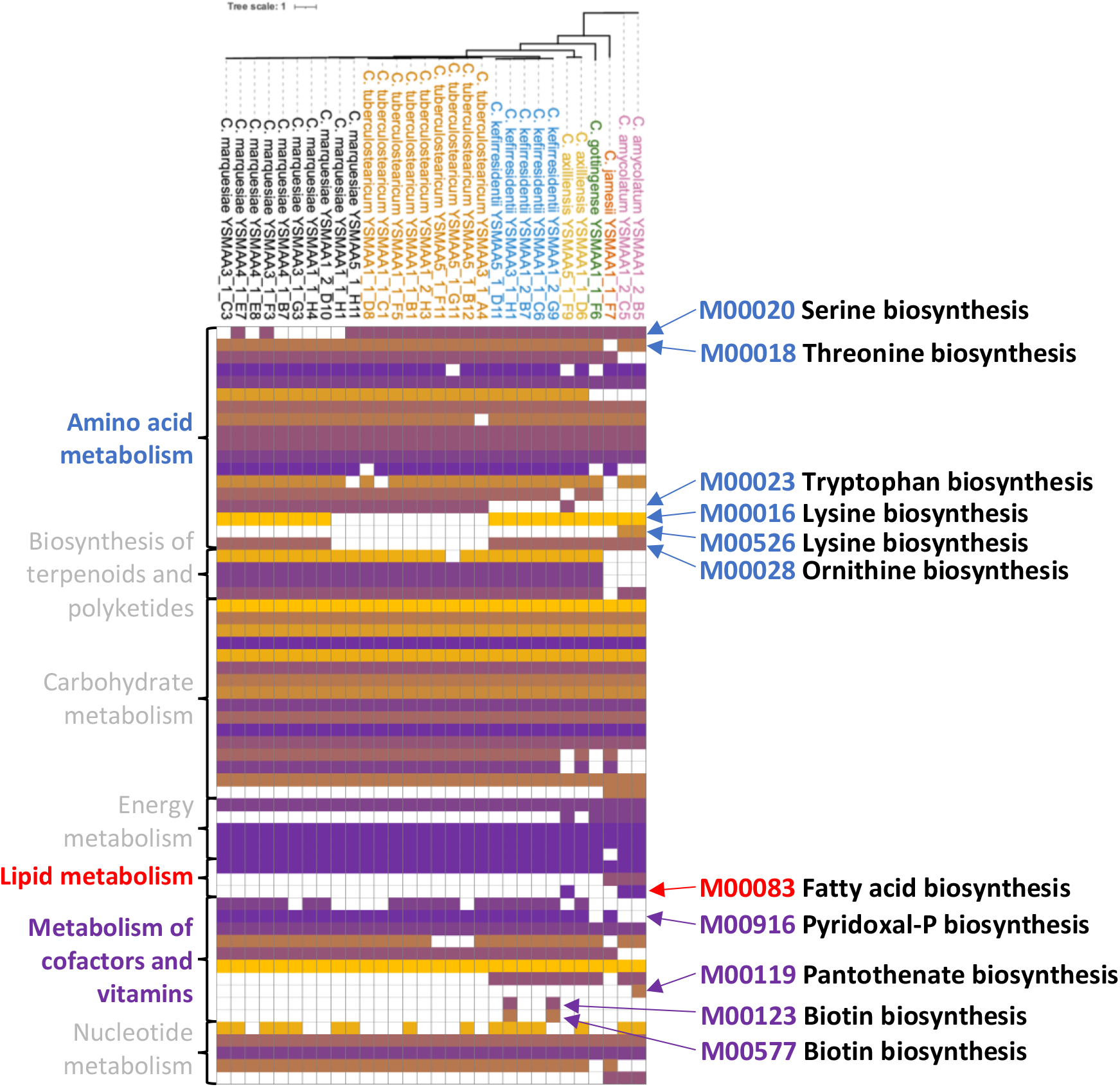
Metabolic pathways in axillary corynebacteria reveals species specific general profiles and some variation within species. Each KEGG module for each isolate were placed on a heatmap where the modules with the largest number of reactions are in yellow and lowest in purple with absent modules in white. Modules of interest were described on the right of the heatmap. KEGG modules were grouped according to the categories on the left of the heatmap. Isolates were grouped by species with a core genome tree depicting the variation.

**Fig. 3A, red shading**). Further, the 16S rRNA (V1-V3 region) sequences of these 24 isolates were found to be ≥ 99.8% identical to three closely related but ultimately different species of corynebacteria which demonstrates the difficulty to speciate some corynebacterial isolates using this method. From here henceforth, the representative genomes will be referred to with the prefix YSMA (**Y**ork **S**kin **M**icrobiome: Study **A**)

While the 16S information alone could not accurate speciate our isolates, we were able to do this using the closed genomes using GTDB-tk [23, 24] (**Supplementary Fig. 3B, Supplementary Table 1**). Using this analysis, we observe further classification of the 24 isolates into three separate species, *C. kefirresidentii* (blue bar), *C. tuberculostearicum* (including *C. tuberculostearicum*_C, orange bar) and *C. aurimucosum*_E (black bar). These three species are commonly found on the skin and are thought to be part of the *C. tuberculostearicum* species complex which was previously reported [21, 25], confirming the importance of these organisms more generally at multiple human skin sites. ANI analysis of the GTDB representative for the species *C. aurimucosum*_E suggest that the species is misassigned and matches best with *C. marquesiae* [26]. From here henceforth, we classify our isolates that matched with *C. aurimucosum*_E on GTDB as *C. marquesiae*. Although two isolates (YSMAA1_1_B1 and YSMAA5_1_F11) matched closest with *C. tuberculostearicum* and *C. tuberculostearicum_C*, they were not classified to a specific species due to the predefined ANI reference radius of the analysis. As they share more than 95% identity to their closest matches, we classified both as *C. tuberculostearicum*.

**Table 1.** The closest phages and bacteria genomes were identified using a BLASTn analysis on the putative prophages identified by geNomad. The percentage sequence identity and percentage coverage of each hit were included.

| Putative prophage name | Putative prophage length | Source species | Closest phage hit (% identity, % query cover) | Closest bacteria hit (% identity, % query cover) |
| --- | --- | --- | --- | --- |
| A1_1_C6_phi1_2 | 52250 | *C. kefirresidentii* | ctNWy2 (92.03%, 69%) | *C. tuberculostearicum* CT (91.51%, 76%) |
| A1_1_D6_phi1 | 42557 | *C. axilliensis* | ctIBL2 (83.98%, 29%) | *C. aurimucosum* DSM 44532 (87.61%, 26%) |
| A1_1_F7_phi1 | 56384 | *C. jamesii* | ctwl93 (77.01%, <1%) | *C. jeikeium* DSM 7171 (91.59%, 7%) |
| A1_1_H1_phi1 | 49192 | *C. marquesiae* | ctfR81 (90.01%, 67%) | *C. aurimucosum* FDAARGOS_1110 (84.54%, 49%) |
| A1_2_B5_phi1 | 38625 | *C. amycolatum* | - | *C. jeikeium* FDAARGOS_574 (92.93%, 37%) |
| A1_2_D10_phi1 | 42401 | *C. marquesiae* | ctKpD5 (81.46%, 43%) | *C. accolens* CA1 (80.24%, 49%) |
| A1_2_G9_phi1 | 26374 | *C. kefirresidentii* | ctOSk1 (95.99%, 64%) | *C. tuberculostearicum* CT (91.72%, 65%) |
| A3_1_C3_phi1 | 38823 | *C. marquesiae* | ctSSL1 (93.44%, 45%) | *C. imitans* NCTC13015 (81.58%, 28%) |
| A3_1_F3_phi1 | 54302 | *C. marquesiae* | ctcbl1 (92.73%, 70%) | *C. accolens* CA9 (75.43%, 38%) |
| A3_1_G3_phi1 | 47617 | *C. marquesiae* | ct1BD1 (92.15%, 54%) | *C. tuberculostearicum* CTNIH13 (89.64%, 42%) |
| A3_1_H1_phi1 | 47347 | *C. kefirresidentii* | ctLCM1 (95.25%, 44%) | *C. tuberculostearicum* CTNIH23 (97.62%, 9%) |
| A4_1_E7_phi1 | 47738 | *C. marquesiae* | ct1BD1 (86.18%, 54%) | *C. tuberculostearicum* CTNIH13 (89.94%, 40%) |
| A5_1_D11_phi5_2_1 | 49670 | *C. kefirresidentii* | ctNWy2 (95.17%, 85%) | *C. tuberculostearicum* CT (99.84%, 100%) |
| A5_1_D11_phi3_4 | 50578 | *C. kefirresidentii* | ctNWy2 (96.26%, 64%) | *C. tuberculostearicum* CT (96.35%, 81%) |
| A5_1_F9_phi1 | 43162 | *C. axilliensis* | ctIBL2 (84.39%, 19%) | *C. aurimucosum* DSM 44532 (88.11%, 56%) |
| A5_1_H11_phi1 | 50212 | *C. marquesiae* | ctXFF1 (97.39%, 85%) | *C. accolens* CA9 (84.87%, 51%) |
| A5_1_H11_phi2 | 49906 | *C. marquesiae* | ct1BD1 (92.69%, 46%) | *C. tuberculostearicum* CTNIH13 (91.20%, 75%) |

YSMAA1_2_B5 and YSMAA1_2_C5 were classified as *C. amycolatum* which have been identified not only on the skin but also specifically in the human axilla [27, 28] (**Supplementary Table 1**). Interestingly, YSMAA1_1_F6 was identified to be *C. gottingense* which was first reported in 2017 [29] but until now, not being associated to the human skin. Finally, three isolates (YSMAA1_1_D6, YSMAA5_1_F9 and YSMAA1_1_F7) were matched with metagenome derived genomes of uncultured isolates suggesting that these are two new species. Isolates YSMAA1_1_D6 and YSMAA5_1_F9, tentatively named *C. axilliensis*, are most closely related to *C. singulare* with an average nucleotide identity of ∼84% calculated using FastANI [30] (**Supplementary Fig. 4A**). *C. axilliensis* isolates were found on individuals A1 (∼4% of corynebacteria genomes) and A5 (∼23% of corynebacteria genomes). Isolate YSMAA1_1_F7, tentatively named *C. jamesii*, is most similar to *C. jeikeium* with an average nucleotide identity of ∼87.5% calculated using FastANI [30] (**Supplementary Fig. 4B**). Overall, through our direct to whole genome sequencing approach, we not only revealed the species diversity of our axillary isolates from the four volunteers which would not be captured if a prior 16S rRNA screen was carried out but were also able to identify species not previously associated to healthy human skin including two novel species.

**Figure 4.**
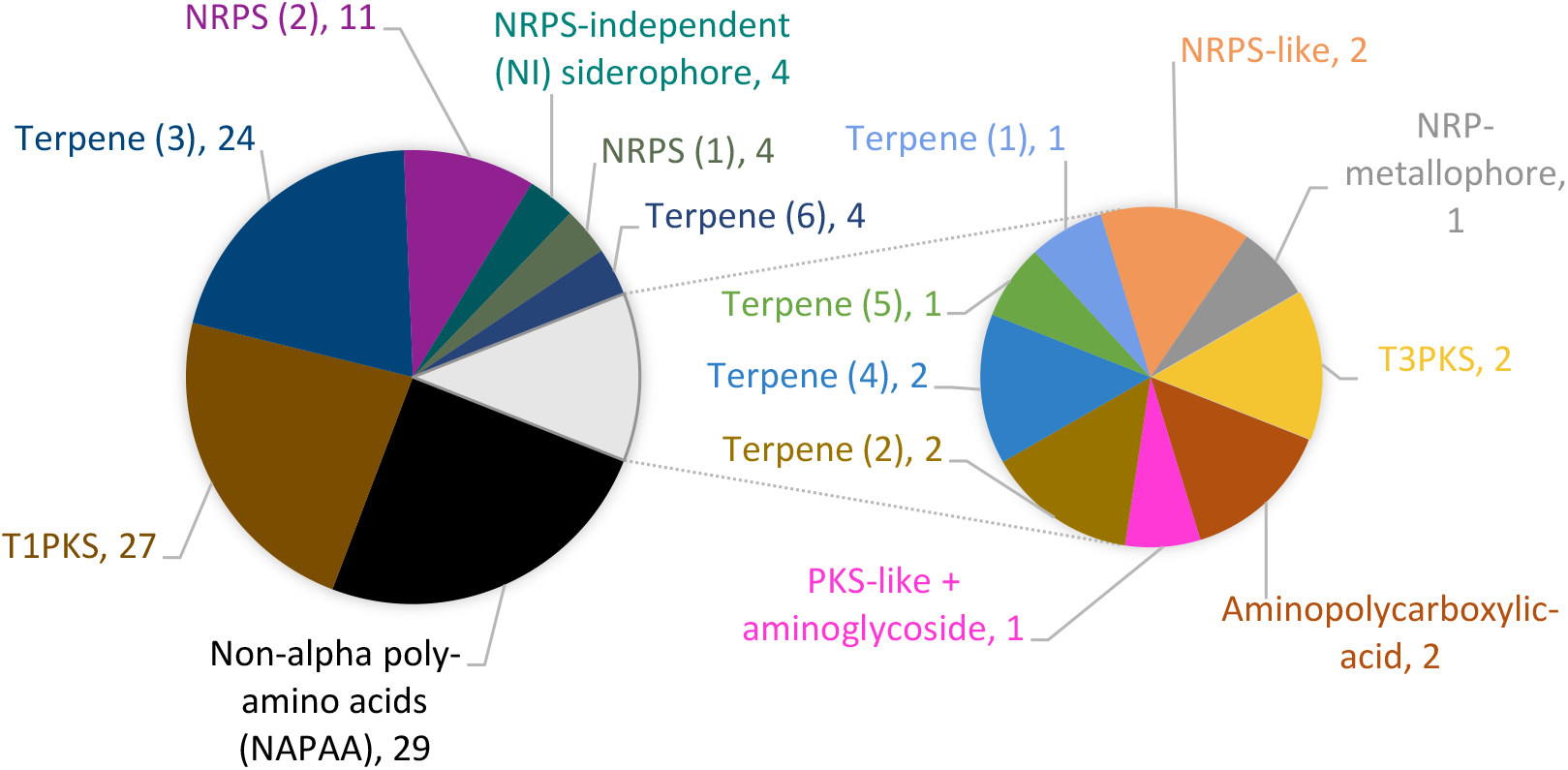
Multiple putative biosynthetic gene cluster (BGC) types were identified in the representative isolate genomes using antiSMASH 7. Names of the different BGC types were included with different cluster architectures in numbered parentheses. The abundance of each BGC type were also included.

### Pangenome analysis of the 30 isolates

We performed pangenome analysis of the 30 representative isolates to understand their genetic differences. Firstly, we used Roary [31] to derive a core genome maximum likelihood SNP tree using a BLASTp cutoff of 80% and seeded the tree with species representatives from GenBank (**Supplementary Fig 5**). Consistent with the GTDB-tk analysis, we clearly see 7 different species including distinctions between the isolates from the species of the *C. tuberculostearicum* species complex and obvious divergence between the two novel species and their nearest related species. To further elucidate the genetic differences between our isolates and their closest GenBank representatives, we performed a pangenome analysis using anvi’o [32] (**Fig. 2**). Gene clusters of interest were then functionally predicted with eggNOG-mapper[33, 34].

**Figure 5.**
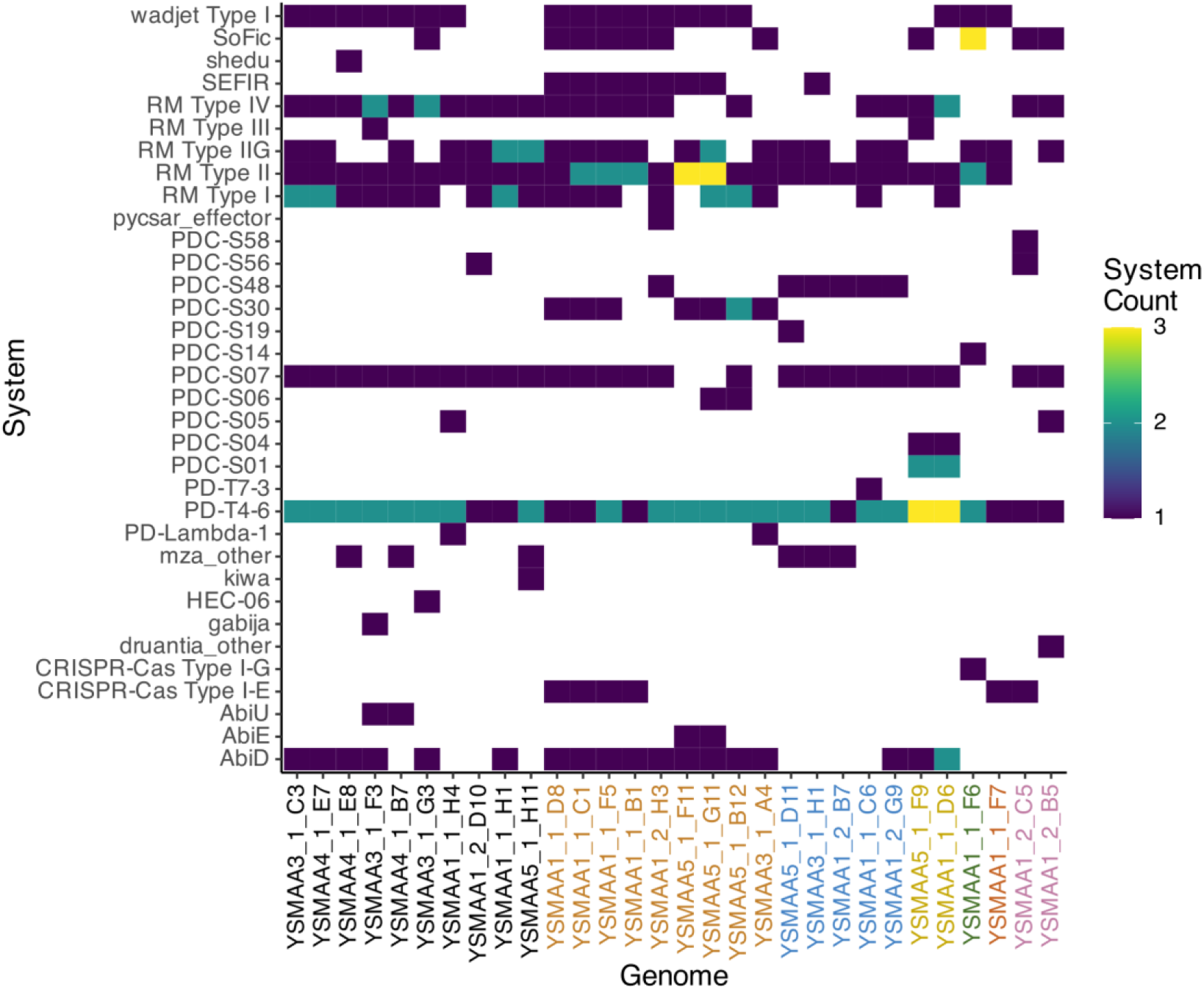
Various phage defence systems were predicted in the representative genomes using PADLOC. The heatmap depicts the abundance of each phage defence system within each isolate according to the scale on the right with absent systems in white. The genomes were arranged according to the different corynebacteria species and coloured accordingly.

The two representative *C. axilliensis* isolates share a specific cluster of genes with the largest proportion of genes with assigned functions found to be associated to transcription (K) or replication, recombination and repair (L) (**Supplementary Fig. 6A**). Interestingly, analysis of the gene clusters specific to either *C. axilliensis* YSMAA5_1_F9 or YSMAA1_1_D6 places the largest proportion of genes with annotated functions in the cell wall/membrane/envelope biogenesis (M) category suggesting differences in their cell envelope (**Supplementary Fig. 6B and 6C**). Functionally annotated gene clusters specific to *C. jamesii* YSMAA1_1_F7 are mainly found in the replication, recombination and repair (L) (**Supplementary Fig. 7**).

As expected, *C. kefirresidentii, C. marquesiae* and *C. tuberculostearicum* isolates share a larger number of gene clusters than with other species. To clarify the main differences between the three species, we generated the core genomes for each species from isolates within our collection followed by a pangenome analysis (**Supplementary Fig. 8**) to extract the singletons found within each species for functional analysis. Within the functionally predicted singletons, genes classified to the inorganic ion transport and metabolism (P) category make up the largest proportions in each of the three core genomes (**Supplementary Fig. 9A-C**) consistent with a previous study on skin isolates of this species complex [21].

We further investigated the putative metabolic differences between these isolates by identifying complete metabolic pathways present in each skin isolate using KEGG (**Fig. 3**). As observed in the pangenome analysis, we generally observe KEGG modules clustering with more similar species. Within the *C. tuberculostearicum* species complex isolates, we see conservation of most KEGG modules but the largest differences seem to lie within the amino acid metabolism pathways. Unlike other species within the complex, most *C. maquesiae* isolates do not seem to possess the serine biosynthetic module (M00020) while *C. tuberculostearicum* lack both lysine biosynthetic modules (M00016, M000526) and the ornithine biosynthesis (M00028) modules and *C. kefirresidentii* lacks the tryptophan biosynthetic module (M00023).

The tryptophan biosynthetic module (M00023) seems to also be absent in isolates all of *C. gottingense, C. jamesii, C. amycolatum* and also *C. axilliensis* YSMAA1_1_D6. Interestingly, the two isolates of *C. axilliensis* seem to possess different metabolic pathway profiles, not limited to tryptophan biosynthesis. Another example is the fatty acid biosynthesis module (M00083), found only in *C. axilliensis* YSMAA5_1_F9 and both *C. amycolatum* isolates.

Another category of metabolic pathways that we observe a large number of differences is the metabolism of cofactors and vitamins. All but two isolates of *C. kefirresidentii* seem to be biotin auxotrophs while *C. gottingense* and *C. amycolatum* isolates were suggested to be pyridoxal phosphate auxotrophs. Finally, *C. amycolatum* YSMAA1_2_B5 seems to be the only isolate that is able to synthesise pantothenate from valine or aspartate (M00119).

Overall, these differences including intra-species variation are captured in the 16 corynebacteria isolates from a single individual A1 (**Supplementary Fig 10**). Although we have observed differences between isolates of the same species, we identified mainly species-specific metabolic pathways which could be used to understand the potential modulate the cutaneous corynebacterial community through metabolic means [37][38][39].

### Potential antimicrobial biosynthetic gene clusters and resistance cassettes

Bacteria often produce complex small molecules that have various functions relating to microbial competition such as antibiotics and siderophores, that are encoded within recognisable biosynthetic gene clusters (BGCs). Resident bacteria of highly complex microbial communities like the skin are known to produce antibiotics to suppress other species in their niche, including preventing colonisation by pathogenic species [35] and conversely, allow for the proliferation of others [36]. We used antiSMASH to identify putative BGCs within the representative set of isolates [37]. For the purposes of this global corynebacteria BGC analysis, we group similar predicted clusters together and determined the abundance of each discrete cluster (**Fig 4**). The majority of the isolates share a predicted non-alpha poly-amino acid (NAPAA), a type 1 polyketide synthase (T1PKS) and a terpene biosynthetic cluster. Other predicted BGCs include non-ribosomal polyketide synthase (NRPS) and multiple terpene clusters with different architectures (**Supplementary Fig. 11 and 12**). The *C. tuberculostearicum* species complex isolates were generally found to possess similar BGCs with *C. tuberculostearicum* isolates having the largest variation (**Supplementary Fig. 13**). As expected, distantly related species like *C. amycolatum, C. gottingense* and *C. jamesii* were predicted to have BGCs that are more distinct or of different classes (**Supplementary Fig. 14**). Despite being outside the *C. tuberculostearicum* species complex, the analysed *C. axilliensis* isolates seemed to also share similar predicted BGCs. With the exception of the cationic peptide ε-Poly-L-lysine (NAPAA), we were not able to confidently predict the products of these BGCs as the most similar known clusters for all of the predicted BGCs shared low percentage similarities with their respective putative clusters. Nevertheless, our analysis suggested different BGCs harboured by the corynebacterial isolates which could contribute to the modulation of the skin microbiome through the production of metabolites like siderophores (e.g. NI siderophore, aminopolycarboxylic-acid, NRPS) and even antimicrobials (e.g. T1PKS, PKS-like + aminoglycoside).

The likely presence of antimicrobials on the human skin may promote the horizontal acquisition of antimicrobial genes or develop beneficial mutants of existing cellular proteins. ResFinder analysis of the isolates revealed 15 isolates with varying numbers and types of acquired resistance genes but no predicted chromosomal mutations leading to resistance (**Supplementary Table 2**). The most common resistance gene identified was *erm(X)* which is proposed to confer resistance to erythromycin-related antibiotics and is widely found across the genus [38–42]. *C. tuberculostearicum* YSMAA5_1_F11 and YSMAA5_1_G11 were also found to have an aminoglycoside resistance gene which could confer resistance to gentamycin and tobramycin. *C. marquesiae* YSMAA1_1_H4, *C. tuberculostearicum* YSMAA3_1_A4 and *C. axilliensis* YSMAA5_1_F9 seem to have a resistance gene to an unknown aminoglycoside initially found in *C. striatum*. Interestingly, *C. axilliensis* YSMAA5_1_F9 has genes conferring resistance to chloramphenicol (cmx), streptomycin (aph(6)-Id, aph(3”)-Ib) and kanamycin (aph(3”)-Ia). These were found in tandem in the same region of the genome surrounded by multiple insertion sequence (IS) elements and were also not found in *C. axilliensis* YSMAA1_1_D6, suggesting a recent acquisition. Finally, we identified a sole tetracycline resistance gene in *C. tuberculostearicum* YSMAA5_1_B12 which also resides near IS elements, suggesting another example of horizontally acquired resistance mechanisms. We attempted to experimentally verify what we identified using ResFinder using the EUCAST disc diffusion assay (**Supplementary Table 3**). Despite the common identification of erm(X) which should confer resistance to clindamycin, only 10 out of the 14 isolates predicted to be resistant were as such. *C. axilliensis* YSMAA5_1_F9 was experimentally resistant to chloramphenicol, as predicted by ResFinder. However, *C. tuberculostearicum* YSMAA5_1_B12 was thought to be doxycycline resistant but did not display the phenotype in our disc diffusion assay. The presence of certain resistance genes may not confer resistance in lab-based disc diffusion assays as we may not have captured the most appropriate conditions in which these isolates would be resistant. Interestingly, *C. jamesii* YSMAA1_1_F7 seem to be experimentally resistant to Penicillin G even though a β-lactamase was not predicted to be in the genome suggesting this isolate has some other innate mechanism of resistance.

### Potential phage clusters and resistance mechanisms

Another type of commonly acquired mobile genetic element are prophages. Phages are commonly found in the microbiome and could play an important role in skin health by regulating the bacterial community. We identified 17 putative prophage sequences in our 30 representative genomes, with an average of 0.6 (± 1.4) prophages per genome [50] (**Table 1**). Putative prophages were identified using geNomad [43] followed by gene annotation using pharokka and phold[44]. Only putative prophages containing genes which were functionally annotated with multiple common features of a phage (e.g. tail proteins, head and packaging) were retained for further analysis. These putative prophages were then searched against the NCBI database using BLASTn to identify the closest related phage and bacteria species. Some putative prophages were either matched with phages but retaining poor coverage (A1_1_F7_phi1, A5_1_F9_phi1) or not matched with any (A1_2_B5_phi1). As expected, various regions of these putative prophages also matched with other corynebacteria and similar to prophage hits, we observe some bacteria hits with poor coverage (A1_1_F7_phi1, A3_1_H1_phi1). As some of these isolates were derived from the same skin site of the same individual, we sought to understand the diversity of these putative prophages as some prophages may be shared between isolates. We performed a multisequence alignment using MAFFT [45] to determine the genetic identities between these putative prophages (**Supplementary Figure 15**). Neither of these putative prophages seem to be shared between isolates, even within the isolates from the same individuals. Remarkably, none of the 9 representative *C. tuberculostearicum* genomes were found to possess any putative prophages.

In addition to the presence of prophages, we analysed the representative genomes with PADLOC [46] to identify potential phage defence systems (**Fig 5**). Type I, II, IIG and IV restriction modification (RM) systems are commonly found in all isolate genomes while the Type III RM system is much less abundant. Other commonly identified systems include the plasmid transformation protection system Wadjet Type I [47], the abortive phage infection system AbiD [48]. We also observed the CRISPR-Cas Type I-E system in some isolates of *C. amycolatum, C. tuberculostearicum* and *C. jamesii* and also the CRISPR-Cas Type I-G system only in *C. gottingense*. We also predicted the presence of multiple low abundance phage defence systems like the putative nucleotide-sensing Gabija defence [49], the pyrimidine cyclase Pycsar [50] and other defence systems of unknown mechanisms like shedu and kiwa [47]. Global PADLOC analysis of all 215 complete genomes of our axillary corynebacteria isolates suggested typical conservation of these predicted phage defence systems within the drep secondary clusters with some exceptions (**Supplementary Fig. 16 - 19**). Some of these differences include select isolates from *C. marquesiae* cluster 4_5 and 4_8 and *C. tuberculostearicum* cluster 3_2 which lack the RM type IV system. A minority of *C. marquesiae* cluster 4_5 isolates were also predicted to have the putative phage RNA interacting Mokosh Type II system [51]. Similarly, only two isolates of *C. kefirresidentii* cluster 2_5 lack the RM Type IIG system and two other isolates of the same cluster were predicted to have the AbiD system. Together, we have comprehensively covered the range of phage defence systems predicted to be present in our axillary isolates and we expect these mechanisms to be present in other skin associated corynebacteria.

## Discussion

The rise in number of recent studies exploring the various roles of commensals in skin health, be it antagonistic or mutualistic, accentuates the need to reveal the true constituents of the skin microbiome. Within the population of skin commensal microbes, staphylococci and cutibacteria have been widely studied since the pioneering skin microbiome studies [2–4, 52]. Even though a large proportion of the microbiomes of multiple types of skin sites are made up of corynebacteria, commensals of this genus have been poorly studied highlighting the importance of studies like this to not only uncover the roles of these commensals but to also discover novel skin associated species. Others have also sought to address this by attempting to clarify the diversity of cutaneous corynebacteria [21, 22, 25] but the genetic information for this genus still lagged behind that of cutaneous staphylococci and cutibacteria. The recent advances in long-read sequencing, particularly the R10.4.1 flow cell and Kit 14 chemistry by Oxford Nanopore Technologies, have allowed for high quality sequencing to be more accessible at lower costs to researchers. With this we developed and optimised a direct-to sequencing method enabling us to contribute another 30-representative cutaneous corynebacteria genomes with another 195 closed genomes which were filtered out during dereplication but are available for analysis.

Unlike our study, prior efforts to understand the skin microbiome usually involves an initial 16S rRNA amplicon sequencing step which acts as a filter to select for specific isolates to proceed with whole genome sequencing. The high throughput nature of 16S rRNA gene amplicon sequencing makes it an appealing first step in determining the diversity of isolates. A common way to compare the 16S rRNA gene sequences is by aligning their respective V1-V3 fragment as it has been shown to be able to distinguish between species more reliably [53, 54]. However, this approach does not resolve the differences between isolates of closely related but ultimately different species. For example, the skin commensals *C. kefirresidentii* strain FDAARGOS_1055 and *C. tuberculostearicum* strain FDAARGOS_1198 share a 16S rRNA (V1-V3 fragment) sequence identity of 99.8% but only share a ∼89% genomic sequence identity when calculated using FastANI. Clearly, the difference between the two species is only obvious when whole genome sequences are available. The difficulties in resolving the diversity of isolates were also previously reported in multiple other studies as microbial “dark matter”, or isolates that cannot easily speciated using the current tools and information available [2, 55, 56]. Further, common mobile elements like prophages which known contributors to genetic variation would be missed without a whole genome sequencing approach. We were able to keep our whole genome sequencing cost low (∼2.5x the cost of 16S rRNA gene sequencing) negating the need for an initial 16S rRNA gene filtering approach as the invaluable genetic information from whole genome sequencing outweighs the additional costs. While our method provides deeper insight into the species and strain diversity within single skin sites on single individuals, we are still likely undersampling all the diversity that is there, while obtaining a much deeper understanding of the recovered microbes and the power of this technique in assessing the complete axilla community will be discussed elsewhere.

Using our method, we were able to strongly confirm the widespread presence of the species *C. tuberculostearicum, C. kefirresidentii* and *C. marquesiae* (formerly *C. aurimucosum*_E) (**Supplementary Fig. 20**), which together form the *C. tuberculostearicum* species complex [26]. In addition, we found more rarely *C. amycolatum* strains in the axilla and unexpectedly, we also identified isolates of *C. gottingense*, a species previously not associated to the skin. This species has been isolated from blood of patients with bacteraemia, the cerebrospinal fluid (CSF) and urine [29, 57], suggesting that it possibly originates from the skin. In addition to known species, we also identified two novel species tentatively named *C. jamesii* and *C. axilliensis*. Isolates of *C. axilliensis* were found on individuals A1 and A5 and makes up a significant 23% of sequenced corynebacterial isolates from the latter suggesting this novel species could be a major previously unrecognised components of the axillary microbiome of some individuals.

Although the fundamental roles of cutaneous commensal corynebacteria are still poorly understood, there have been increasing studies demonstrating how bacteria from this genus can contribute to colonisation protection including against other human pathogens. For example, *C. striatum* has been demonstrated to shift *S. aureus* towards commensalism through the suppression of the *agr* quorum sensing system [14]. In the presence of *C. striatum, S. aureus* exhibited increased human epithelial cell adhesion, decreased haemolysin activity and decreased fitness during infection indicating a decrease in virulence as compared to a monoculture. Rosenstein *et al*. [36] have proposed a cooperative role of corynebacteria against *S. aureus* in the nasal cavity with some species of commensal corynebacteria aiding in the proliferation of commensal *Staphylococcus lugdunensis* which in turn produces the antibiotic lugdunin that is active against *S. aureus*. These studies have highlighted the importance in understanding the true diversity of cutaneous corynebacteria species as we may yet uncover more roles that could be exploited to improve tools and treatments for skin health.

Studies advancing our understanding of the skin microbiome including reports focussing on cutaneous corynebacteria seem to only be scratching the surface of the skin, the biggest organ of the human body. Our knowledge of commensal corynebacteria lags behind other equally prevalent genera like *Staphylococcus* and *Cutibacterium*. Here, we sought to further understand the genetic diversity of the genus *Corynebacterium* using a low cost direct-to whole genome sequencing approach on isolates derived from a single skin site, the axilla. Using the invaluable genetic information, we not only identified species never before seen on the skin including two novel species but we also identified a multitude of genetic differences within all 215 closed genomes including putative metabolic differences, the presence of biosynthetic gene clusters and prophages providing multiple new avenues of exploration. These publicly available genomes can be further mined by other researchers to build on our analysis to further elucidate the roles of this poorly understudied genus on the skin. Further, these and other matched isolates of the York Skin Microbiome (YSM) library are made available for multiple other studies to facilitate our efforts to understand the complex environment that is the skin microbiome.

## Materials and Methods

### Sampling procedures

Four adult male volunteers (Age range: 32 to 78) were recruited for axillary swabbing at Unilever R&D, Bebington, UK. The volunteers were in good general health, did not use any body products (e.g. deodorants) for 24 hours before the study, were not on antibiotics, with no active skin conditions on any part of their bodies, not suffered from eczema in the last 5 years nor have they ever had psoriasis. Prior to swabbing, the volunteers were “sniff tested” by a panel of testers and scored according to odour levels (High malodour: A1 and A3, Low malodour: A4 and A5). Axillary swabs were then collected by swabbing in a linear motion 20 times using eSwabs (Copan) on both underarms and then placed into Amies transport medium.

### Bacterial culturing

Swabs were plated onto ACP solid media (39.5 g L^−1^ Blood Agar base no. 2, 3 g L^−1^ yeast extract, 2 g L^−1^ glucose, 5 mL L^−1^ Tween 80, 50 mL L^−1^ defibrinated horse blood, 100 mg L^−1^ fosfomycin) to enrich for corynebacteria and grown for 2 days at 37°C. Colonies were picked and grown overnight in brain-heart infusion broth + 1 % Tween-80 (BHIT) at 37°C, shaking at 200 rpm. Cultures were stocked in BHIT supplemented with 10 % glycerol in 2 mL 96 well plates.

### DNA extraction

For genomic and plasmid DNA extraction, 0.5 mL of BHIT was inoculated with each isolate in 2 mL 96 well plates and grown for 20 hours at 37°C, shaking at 200 rpm. Cultures were then pelleted by centrifugation before resuspension in 100 μL enzymatic lysis buffer (50 mM Tris-HCl, pH 8, 50 mM EDTA, pH 8, 0.5% Tween 20, 0.5% Triton-X100, 25 mg/mL lysozyme). The mixtures were incubated at 37°C overnight with gentle shaking (<50 rpm). 500 μL of lysis buffer (0.5 M NaCl, 100 mM Tris-HCl, pH 8, 50mM EDTA, pH 8, 1.5% SDS) was then mixed into each well and incubated at 80°C for 30 minutes in a thermomixer. The mixture was allowed to cool before the addition of 4µl of 4mg/mL RNase A Solution (Promega) before incubation at 37°C for a further hour. Cellular proteins were then degraded with 20 µl of 20 mg/mL Proteinase K Solution (Promega) and incubation at 56°C for 15 minutes. Remaining proteins were precipitated with 200 µl of protein precipitation buffer (5 M Potassium acetate). The lysates were then clarified by centrifugation at 16,000 xg for 5 minutes. DNA was precipitated from the supernatant using 600 µl isopropanol and incubated overnight at 4°C. The precipitated DNA were pelleted by centrifugation at 16,000xg for 5 minutes. The DNA pellets were washed with 600 µl 70% ethanol and then air dried for 15 minutes. Finally, the DNA pellets were rehydrated with 50 µl nuclease-free water and incubation overnight at room-temperature. The DNA concentrations and QC measurements were performed using Qubit (Thermo Fisher) and Nanodrop (Thermo Fisher).

### DNA library preparation, whole genome sequencing and assembly

The Oxford Nanopore Native Barcoding Kit 96 V14 was used to prepare and barcode the prepared genomic and plasmid DNA. The recommended protocol was followed with the following changes: 1. Half the recommended volume of the DNA repair and end-prep reaction was prepared. 2. Half the recommended volume of the native barcode ligation reaction was prepared. The prepared library was loaded into a PromethION R10.4.1 flow cell and sequencing was allowed to take place over 72 hours and base-called using the singleplex high-accuracy model, 400 bps on MinKNOW 23.07.5. The reads were assembled with the EPI2ME Labs wf-bacterial-genomes isolate workflow (Flye [58], Medaka (Oxford Nanopore Technologies), Prokka [59], ResFinder [60, 61]).

### Bioinformatics tools

Pairwise genome comparisons of all 215 assembled genomes were performed using the ANIm [62] analysis on pyANI v0.2.12 [63]. The genomes were dereplicated with drep v3.4.2 [64] using Mash [65] and MUMmer 3.0 [66], employing a primary clustering ANI cutoff of 95% to distinguish between different species and a secondary clustering ANI cutoff of 99.5% to capture strain diversity to derive a representative set of genomes. The core genomes were determined using Roary[31] with an 80% BLASTp cutoff. PhyML 3.3 [67] trees 16S rRNA V1-V3 and core genome alignments of the representative genomes were constructed using the HKY85 substitution model with approximate likelihood ratio tests (aLRT). Trees were visualised on iTOL v6.9 [68]. Speciation using GTDB-tk v2.3.2 [23] was performed on KBase [24]. The genome alignments of *C. axilliensis* and *C. jamesii* were prepared using BLASTn and visualised using BRIG [69].

Pangenome analysis was performed using the anvi-pan-genome package and visualised on anvi’o v8 [32]. The representative genomes were also compared using pyANI and visualised with the pangenome on anvi’o. Relevant gene clusters were extracted and functionally analysed with eggNOG-mapper v2 [70, 71] to determine the respective COG categories. Metabolic pathways were identified using blastKOALA and KEGG Reconstruct [72, 73]. Biosynthetic gene cluster analysis was performed using the default settings on antiSMASH v7 [37]. Putative prophages were predicted using geNomad v1.8 [43] and then annotated using pharokka 1.7.2 and phold [44]. Phage defence systems were predicted using PADLOC v4.3 [74].

### Antibiotic susceptibility testing

Antibiotic susceptibility testing was performed using plate-based disc-diffusion assays using the recommended manufacturers protocol and EUCAST instructions. Overnight cultures were prepared BHIT and grown at 37°C. Cultures were then diluted to a 0.5 McFarland standard (Remel, Thermo Scientific) before plating onto BHIT-agar. Six antibiotic susceptibility discs were placed on the agar (Doxycycline-30μg, Chloramphenicol-30μg, Fosfomycin-200μg, Clindamycin-10μg, Vancomycin-30μg, Penicillin G-1μg) (Oxoid). Plates were allowed to grow for 24 hours at 37°C before measurements of any zones of inhibition.

## Ethics statement

This study was conducted in accordance with the ethical principles of Good Clinical Practice and the Declaration of Helsinki. The local ethics committees of Unilever R&D (Port Sunlight, UK) approved the protocols before commencement of the studies and all subjects gave written informed consent.

## Data availability

The 24 dereplicated genomes and associated plasmids were deposited to GenBank within BioProject PRJNA1191350.

## Notes

### Competing Interest Statement

The authors declare that Barry Murphy and Robert Cornmell are current employees of Unilever.

### Summary of Updates

Authorship and section headings were edited. A data availability section was added to include the accession numbers for the dereplicated genomes used in this study. Line numbers were added.

