## Supplementary Data for "Revealing the diversity of commensal corynebacteria from a single human skin site"

**Supplementary Figure 1. Pairwise average nucleotide identities (ANI) of all isolates sequenced in this study.** ANI values were placed onto a collective coloured pairwise matrix according to the colour scale depicted Three major groups of isolates were identified with I, II and III. Group I II and III isolates from each of the four individuals were denoted with the coloured bars as indicated in the legend.

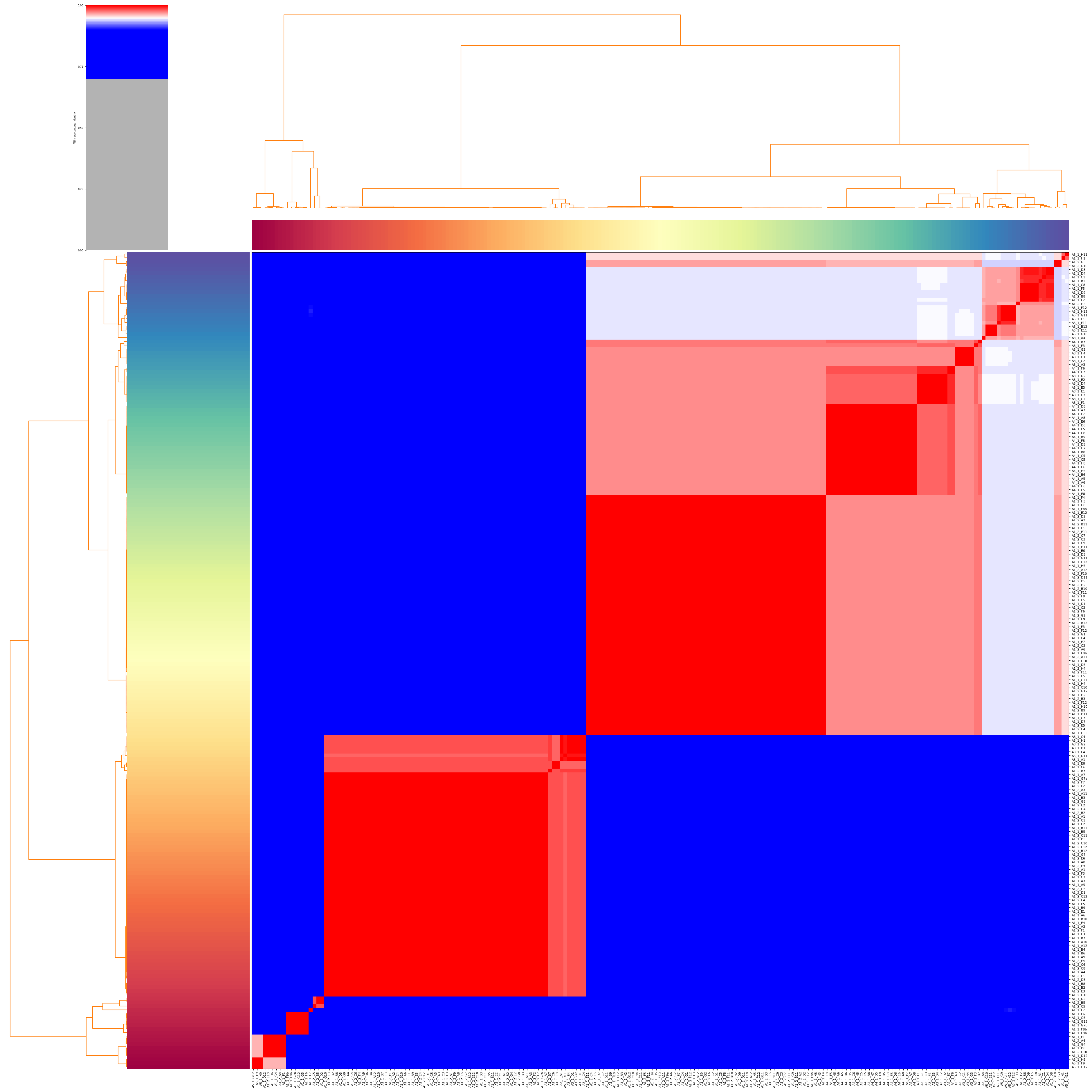

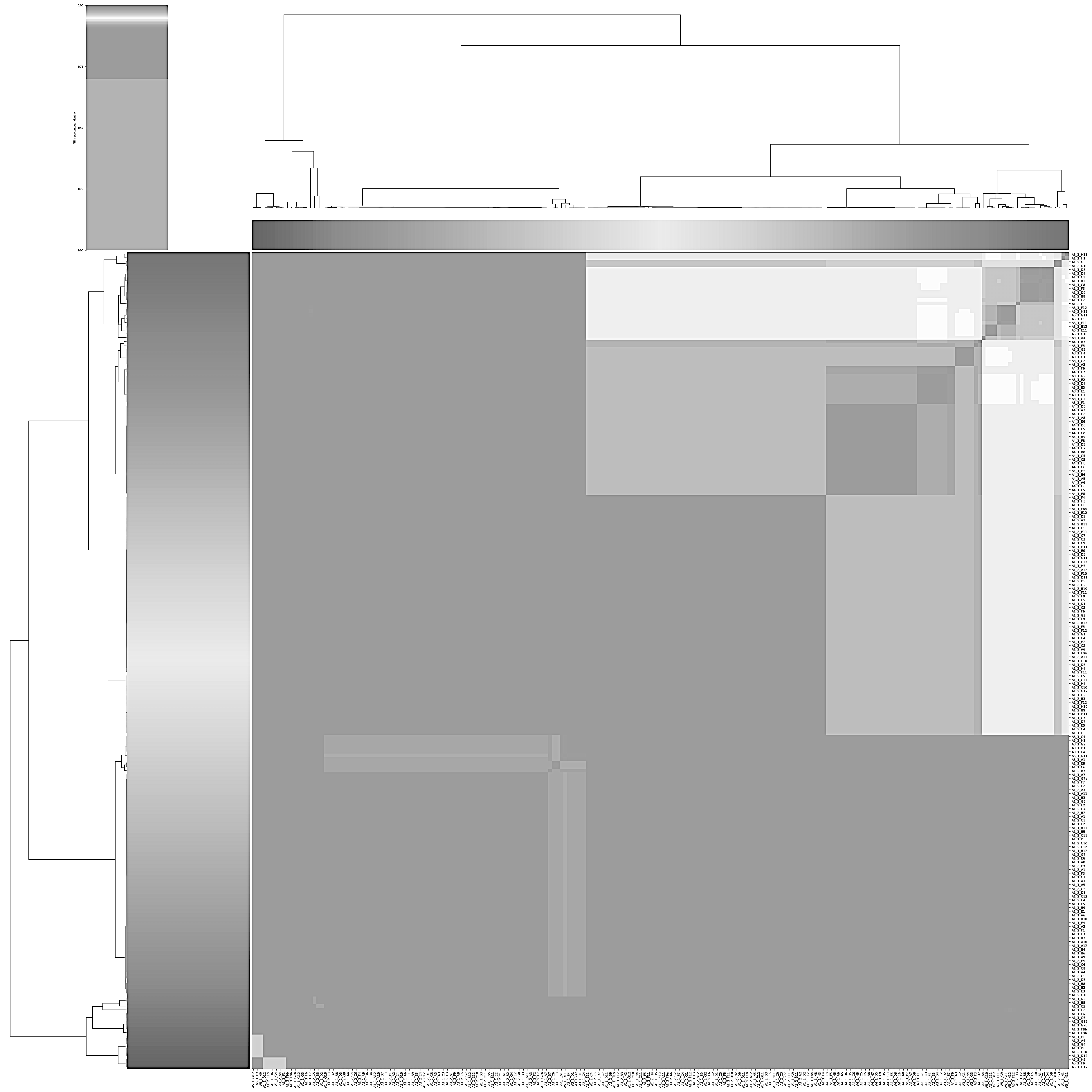

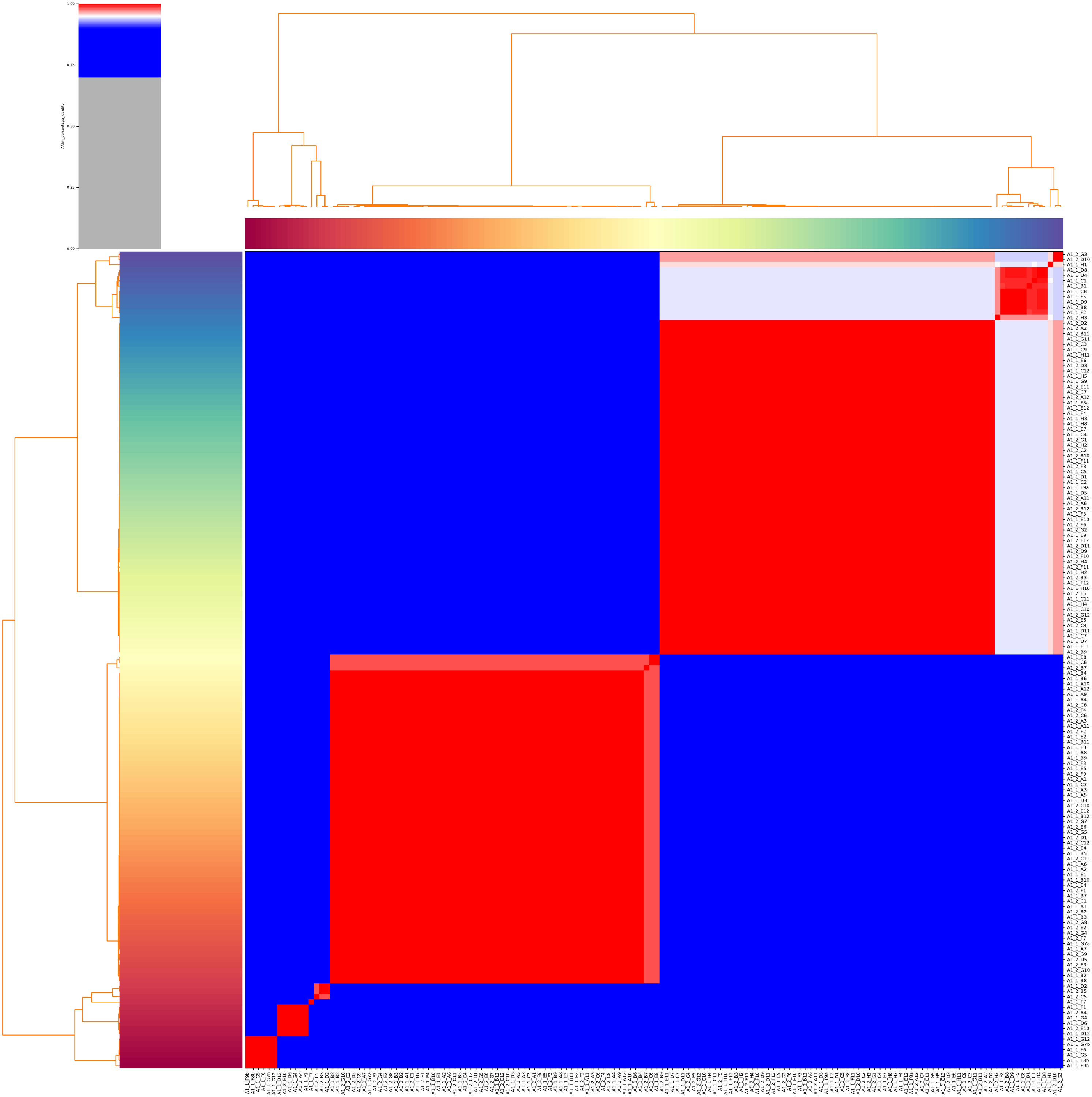

100%

≤ 90%

I

II

III

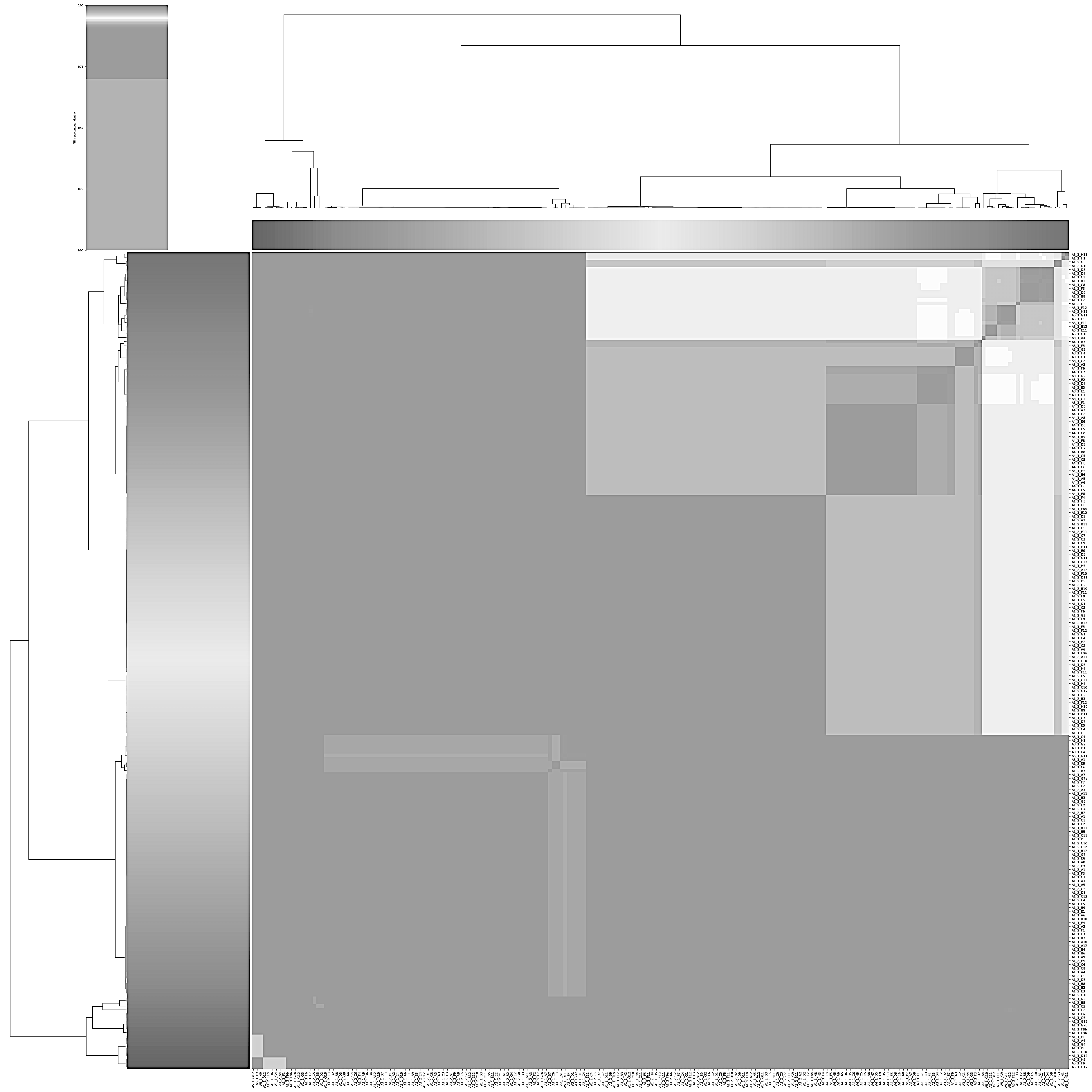

A1

A3

A4

A5

I

II

III

Legend:

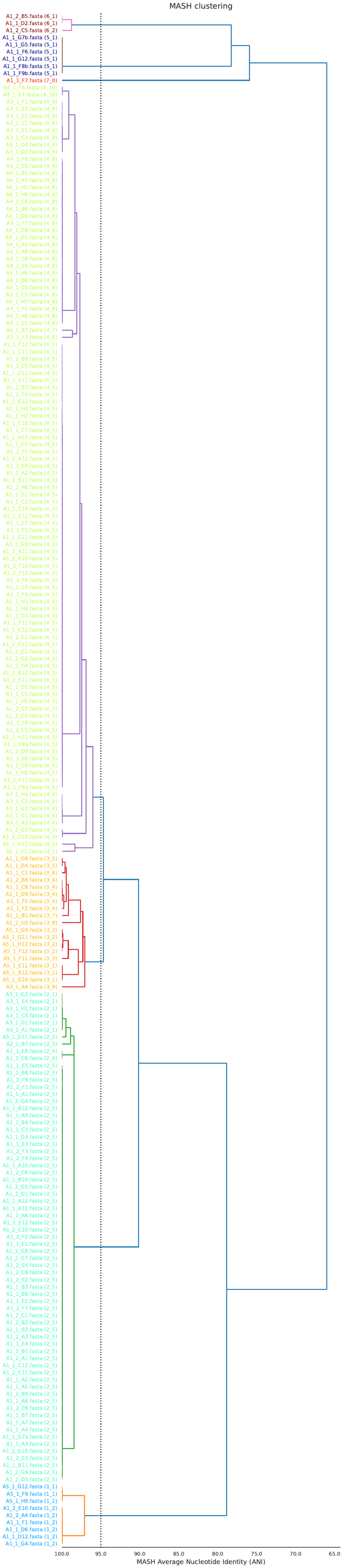

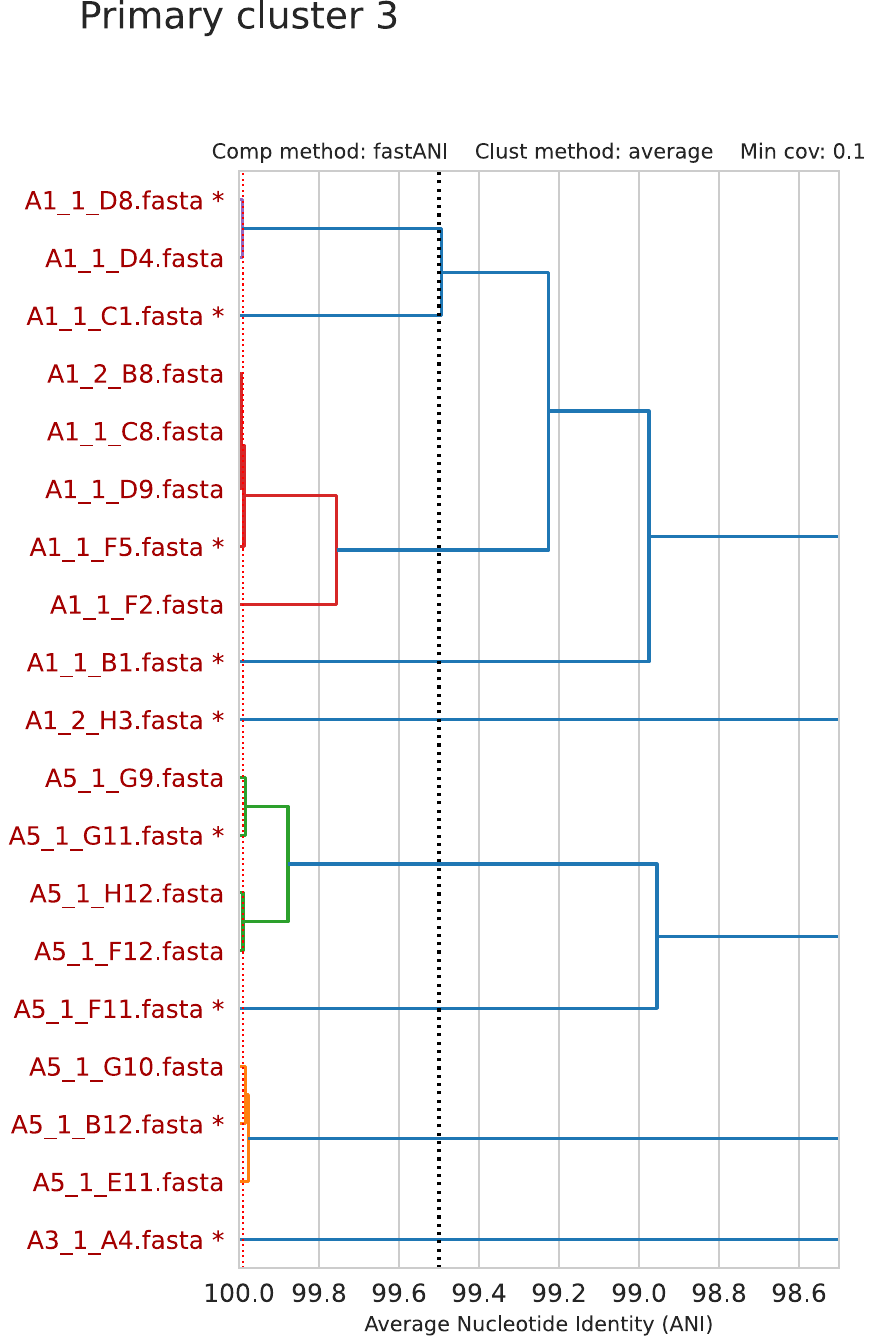

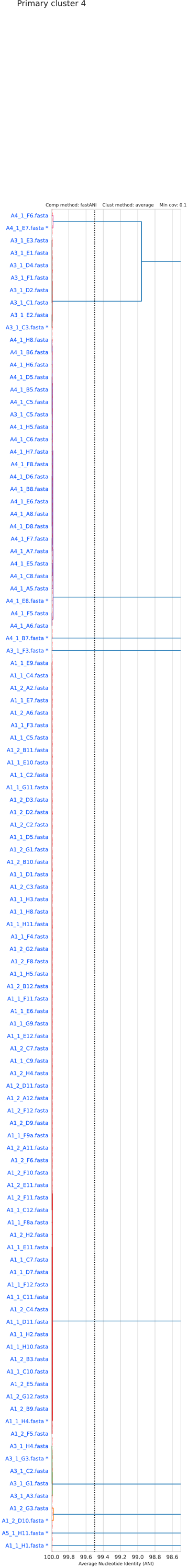

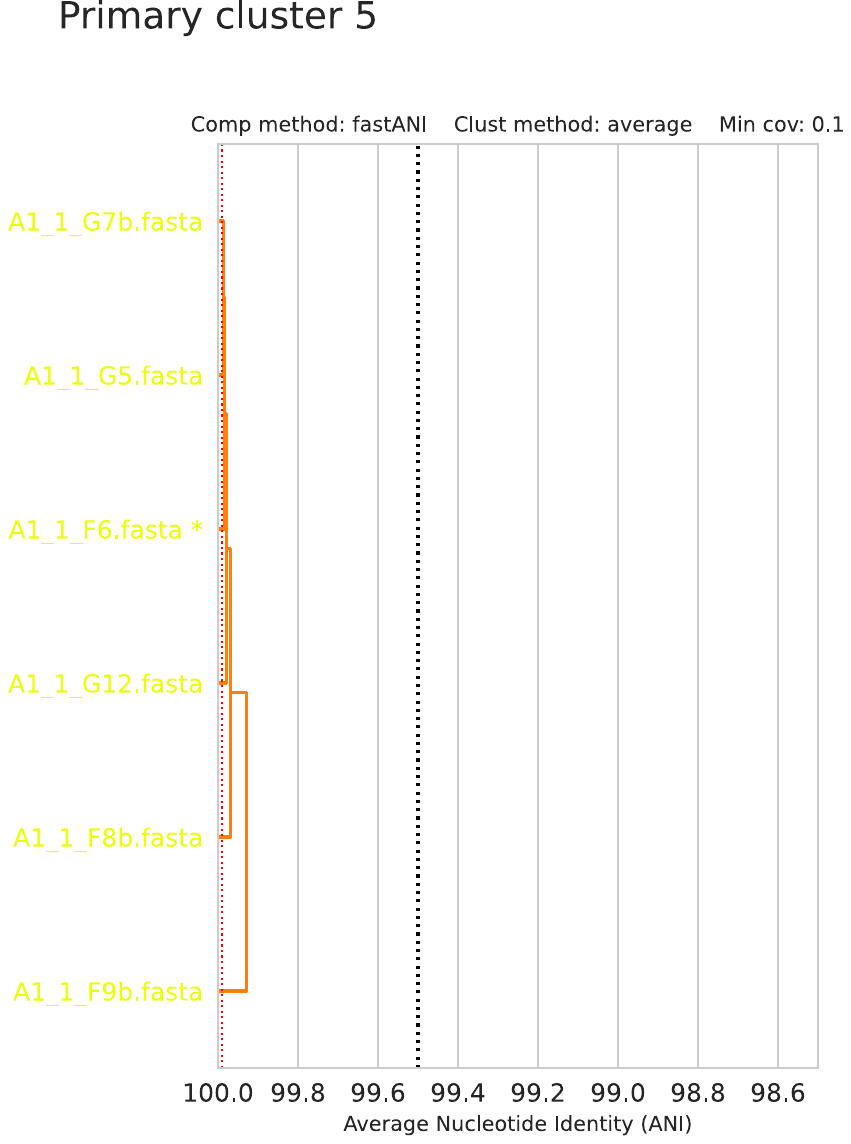

A1_1_G7b.fasta

A1_1_G5.fasta

A1_1_F6.fasta *

A1_1_G12.fasta

A1_1_F8b.fasta

A1_1_F9b.fasta

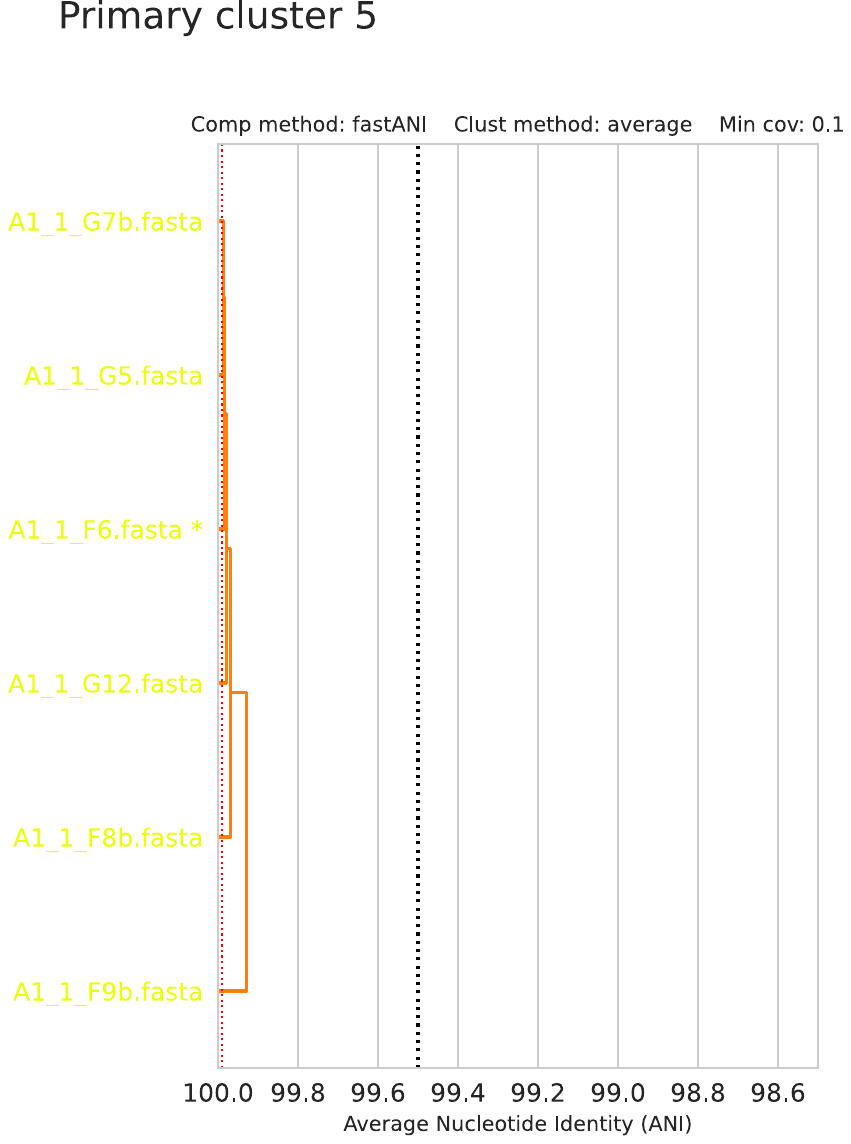

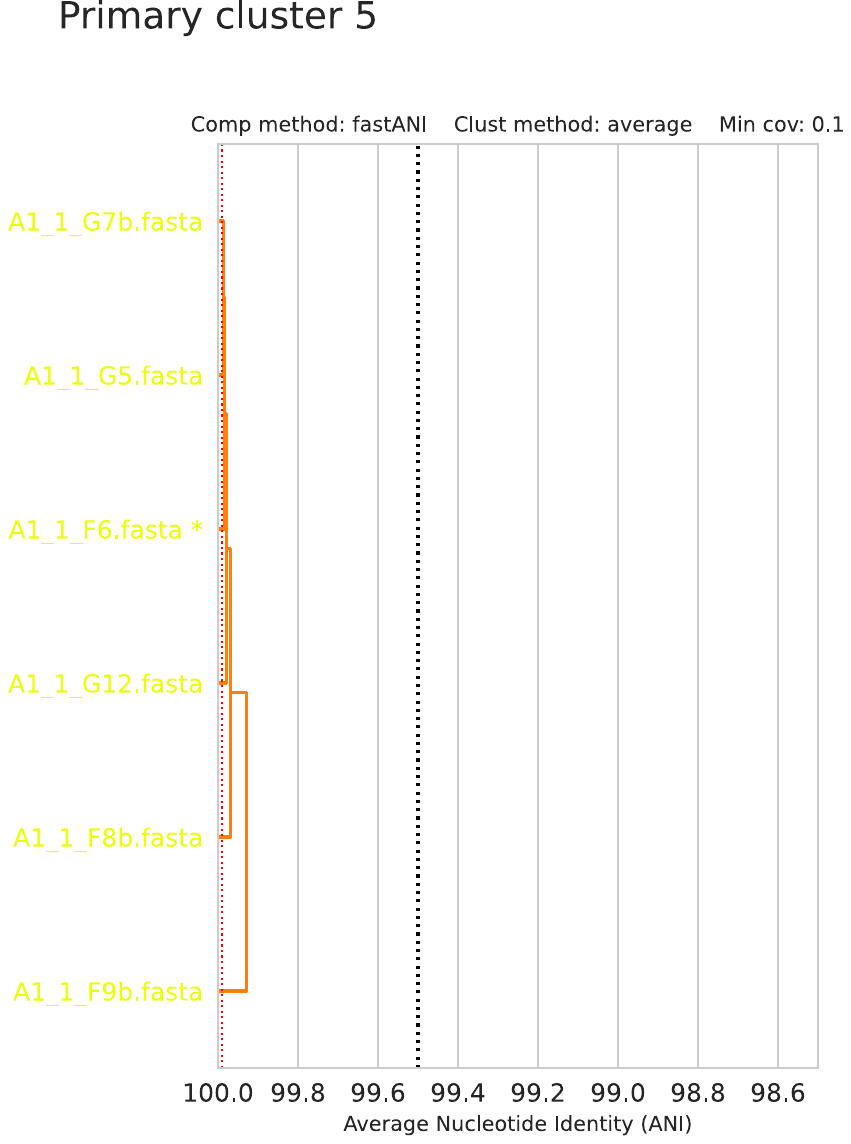

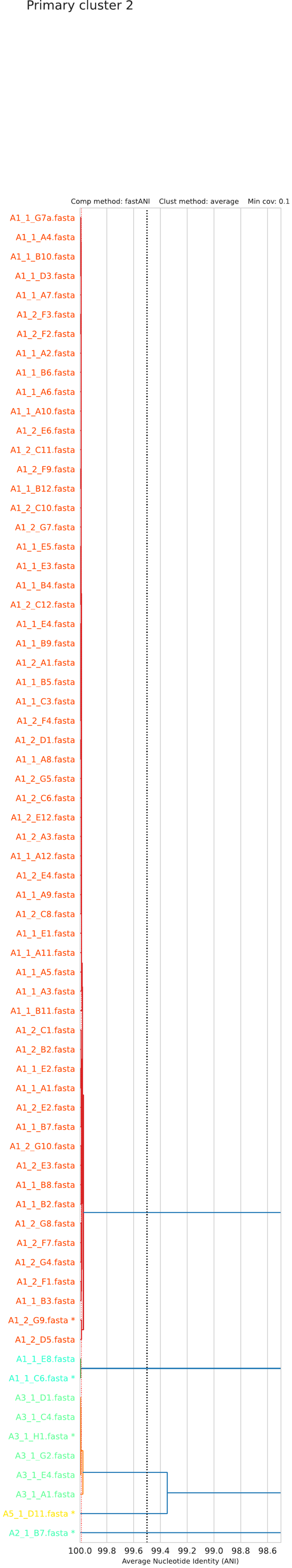

A1_1_E8.fasta

A1_1_C6.fasta *

A3_1_D1.fasta

A3_1_C4.fasta

A3_1_H1.fasta *

A3_1_G2.fasta

A3_1_E4.fasta

A3_1_A1.fasta

A3_1_D11.fasta *

A1_2_B7.fasta *

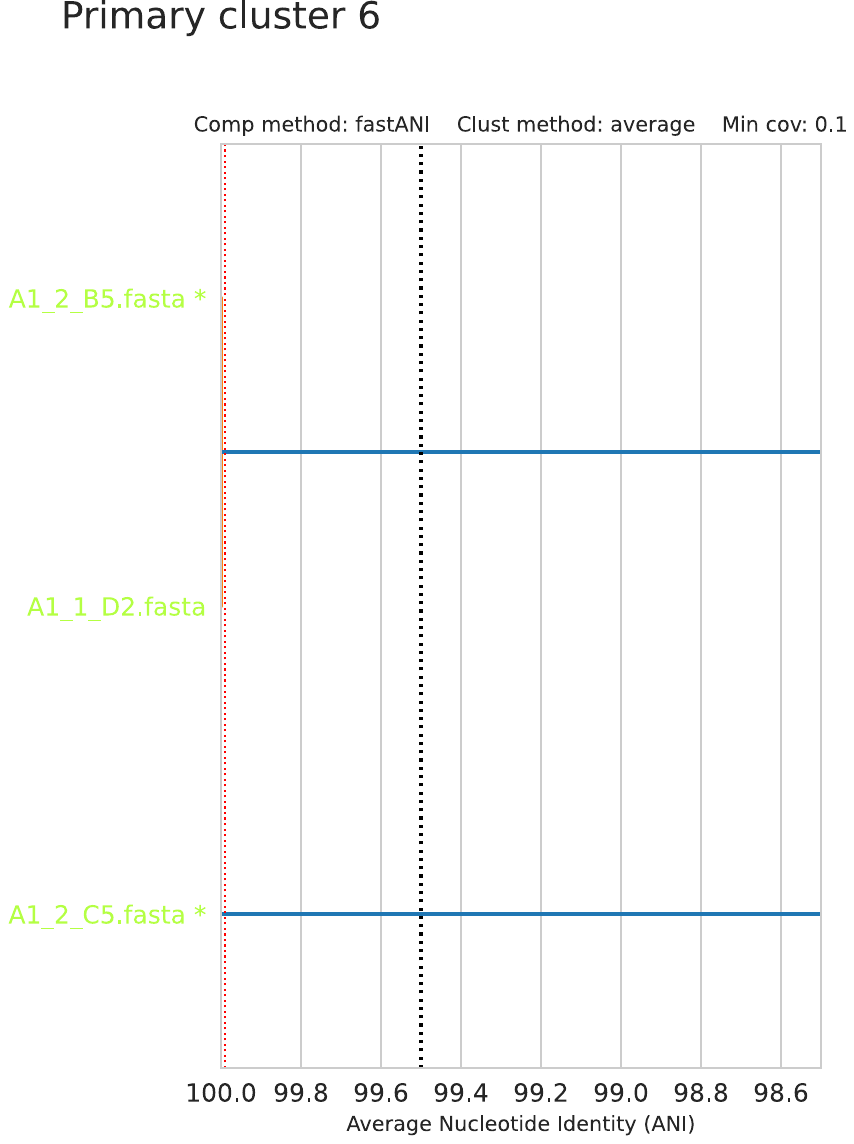

A1_2_B5.fasta *

A1_1_D2.fasta

A1_2_C5.fasta *

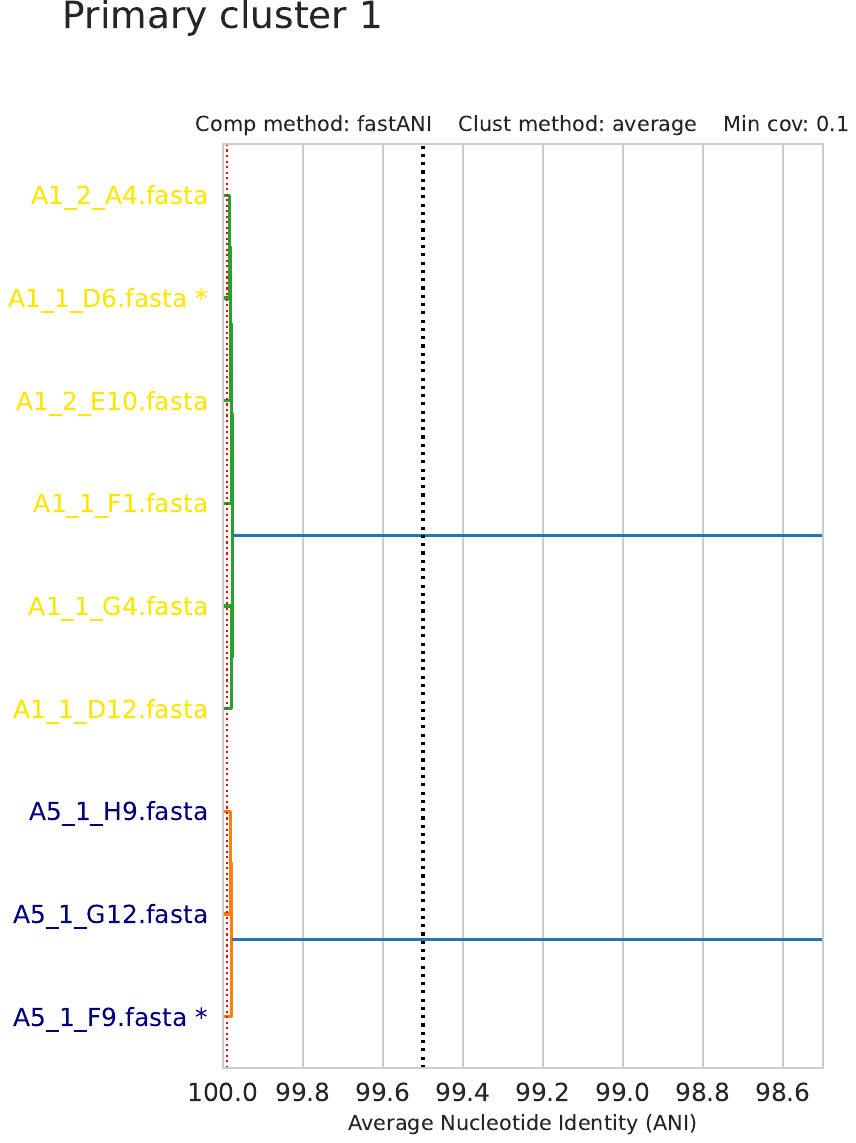

A1_2_A4.fasta

A1_1_D6.fasta *

A1_2_E10.fasta

A1_1_F1.fasta

A1_1_G4.fasta

A1_1_D12.fasta

6

5

4

3

2

1

7

*

*

*

*

*

*

*

*

*

*

*

*

*

*

*

*

*

*

*

*

*

*

*

*

*

*

*

*

*

*

Primary clustering

Secondary clustering

**Supplementary Figure 2.** Isolates were initially clustered by ANI to 7 primary clusters (species level) using MASH at a 95*%* threshold. A secondary clustering was performed on the isolates using fastANI at a 99.5% threshold to identify representatives from each cluster (labelled with *).

**A**

**B**

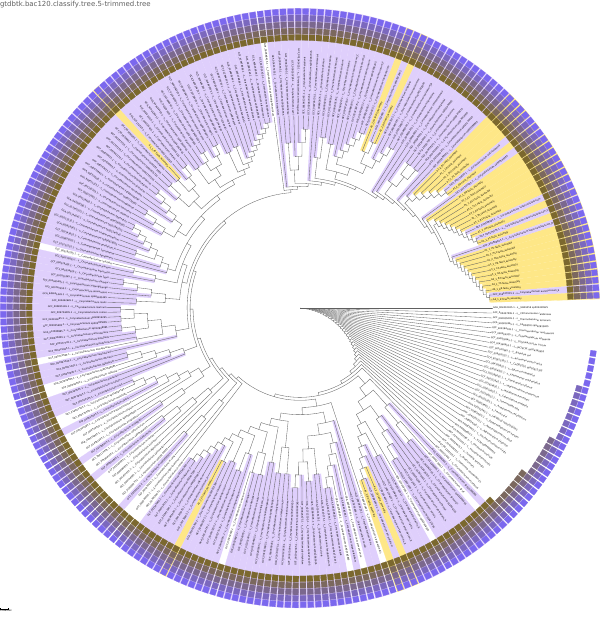

0.704

0.05

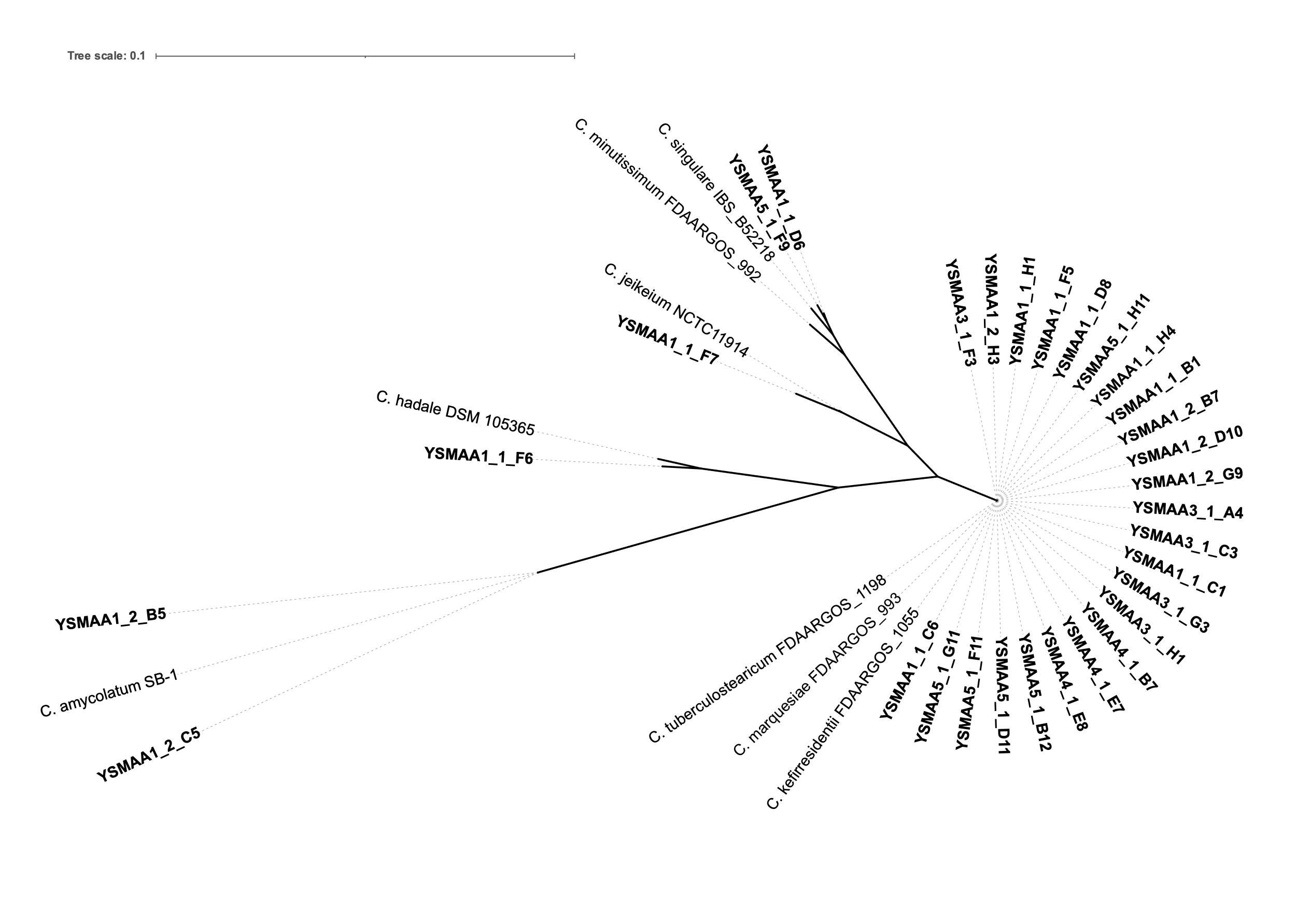

**Supplementary Figure 3. Speciating the representative isolates.** (A) The V1-V3 regions of the 16S rRNA sequences for each genome were aligned with those of the closest RefSeq genome BLASTn hits and plotted onto an unrooted tree. Branches within the red cloud share ≥ 99.8% identity. The tree was generated using PhyML and visualised on iTOL. (B) Genomes of the representative isolates (yellow) were compared to the genomes of the GTDB. 7 distinct groups of isolates were identified with coloured bars in the outermost ring.

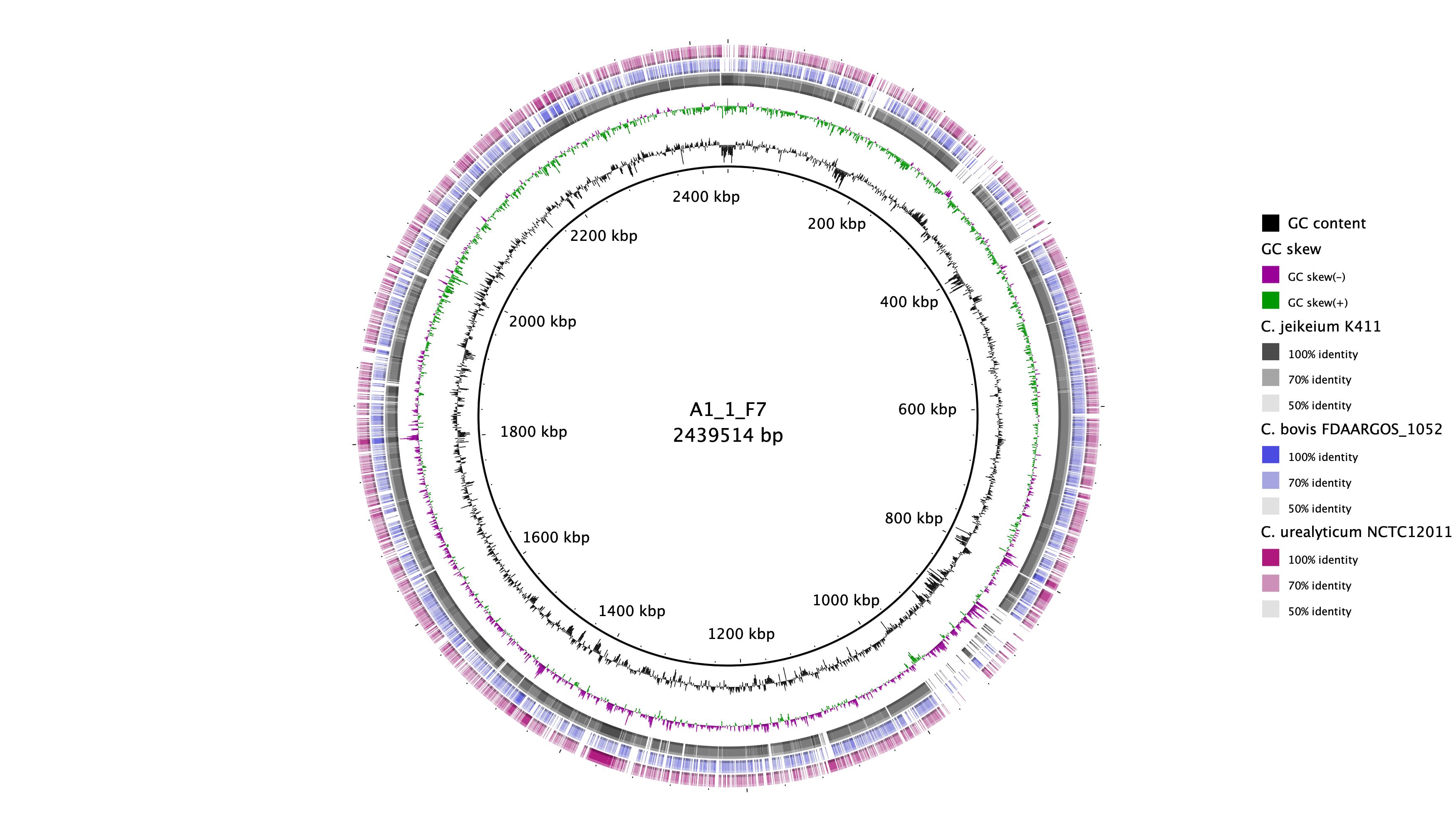

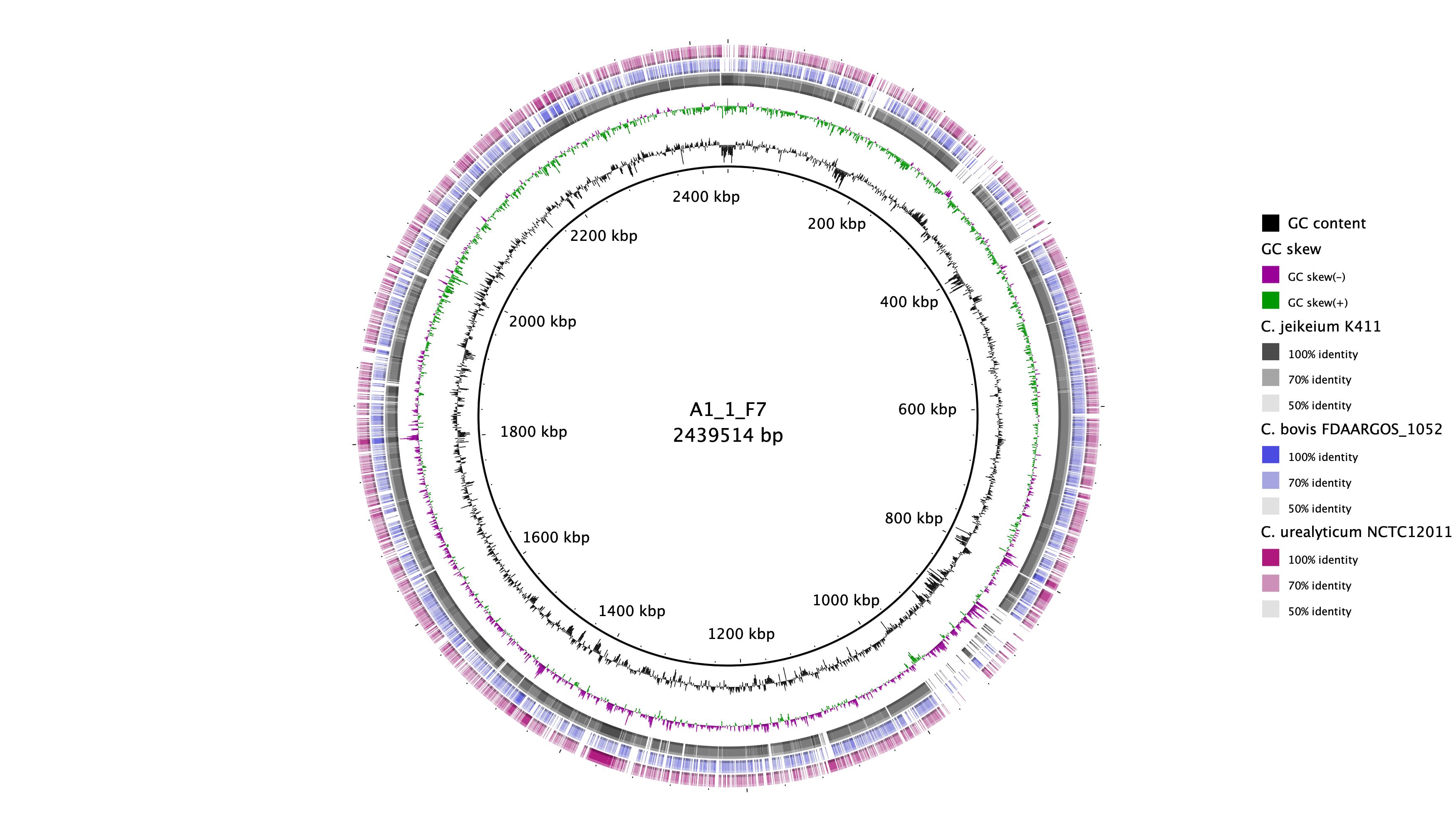

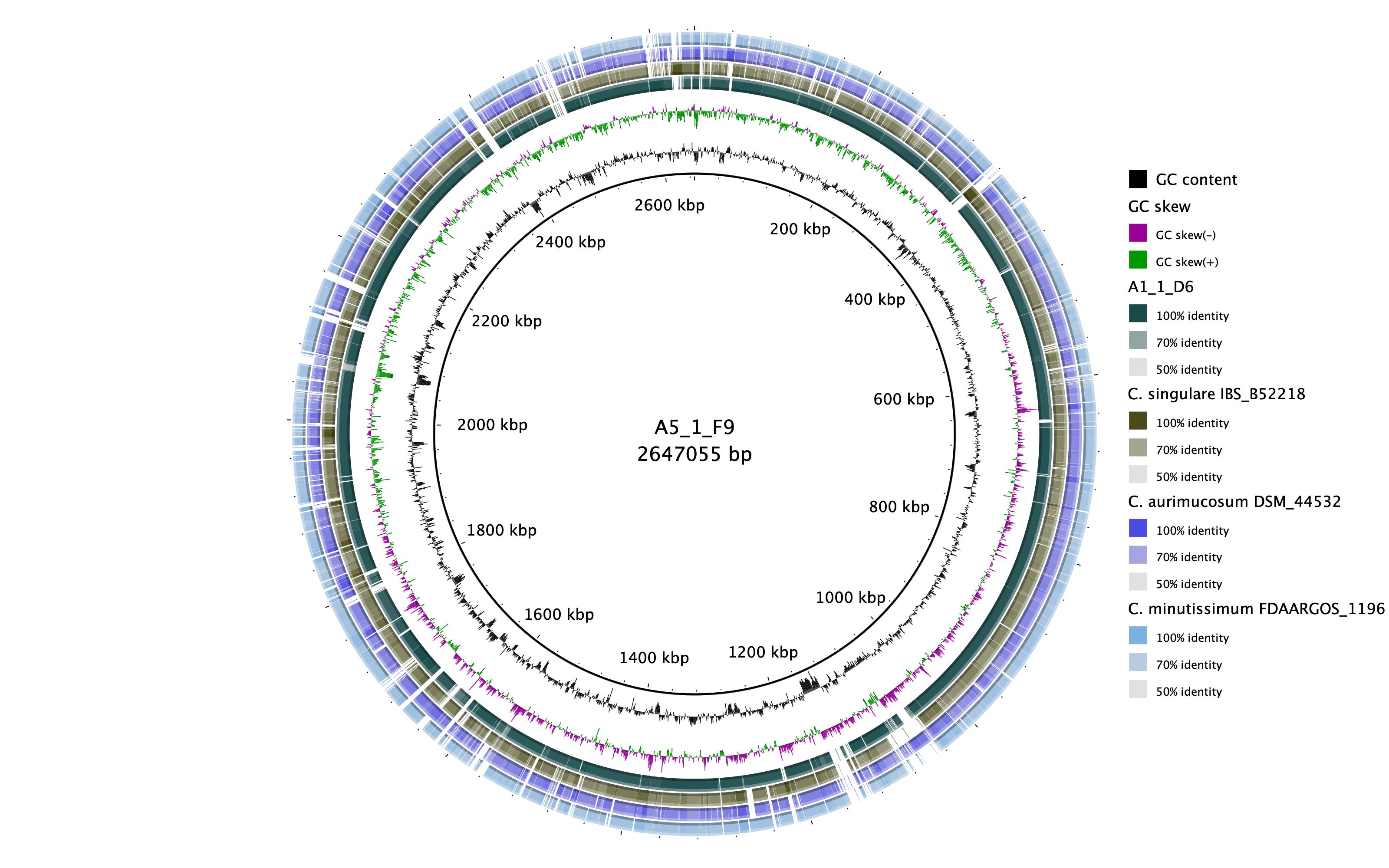

*C. axilliensis*

YSMAA5_1_F9

*C. jamesii*

YSMAA1_1_F7

**A**

**B**

**Supplementary Figure 4. Alignments of representatives from the two novel species *C. axilliensis* and *C. jamesii* against closest hits from GenBank.** *C. axilliensis* YSMAA5_1_F9 and *C. jamesii* YSMAA1_1_F7 were used as the reference genomes on the plots. *C. axilliensis* YSMAA1_1_D6 was also included in (A) for comparison. Alignments and figures were generated using BRIG.

**Supplementary Figure 5. A core genome alignment of the representative genomes (bold) and select RefSeq genomes was generated using Roary with a 80% BLASTp cutoff.** The 7 species were identified using coloured wedges and the species within the *C. tuberculostearicum* species complex (grey, brown and blue wedges) were also distinguished. The tree was generated using PhyML and visualised on iTOL.

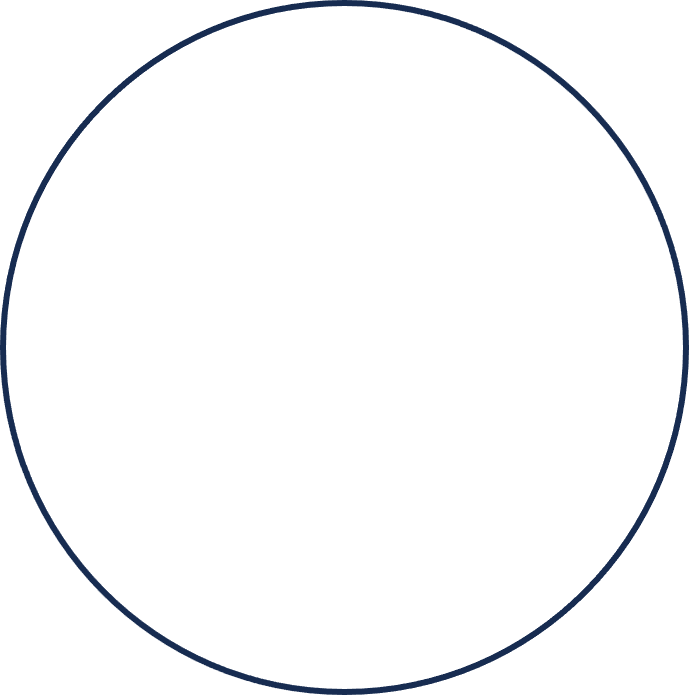

***C. tuberculostearicum species complex***

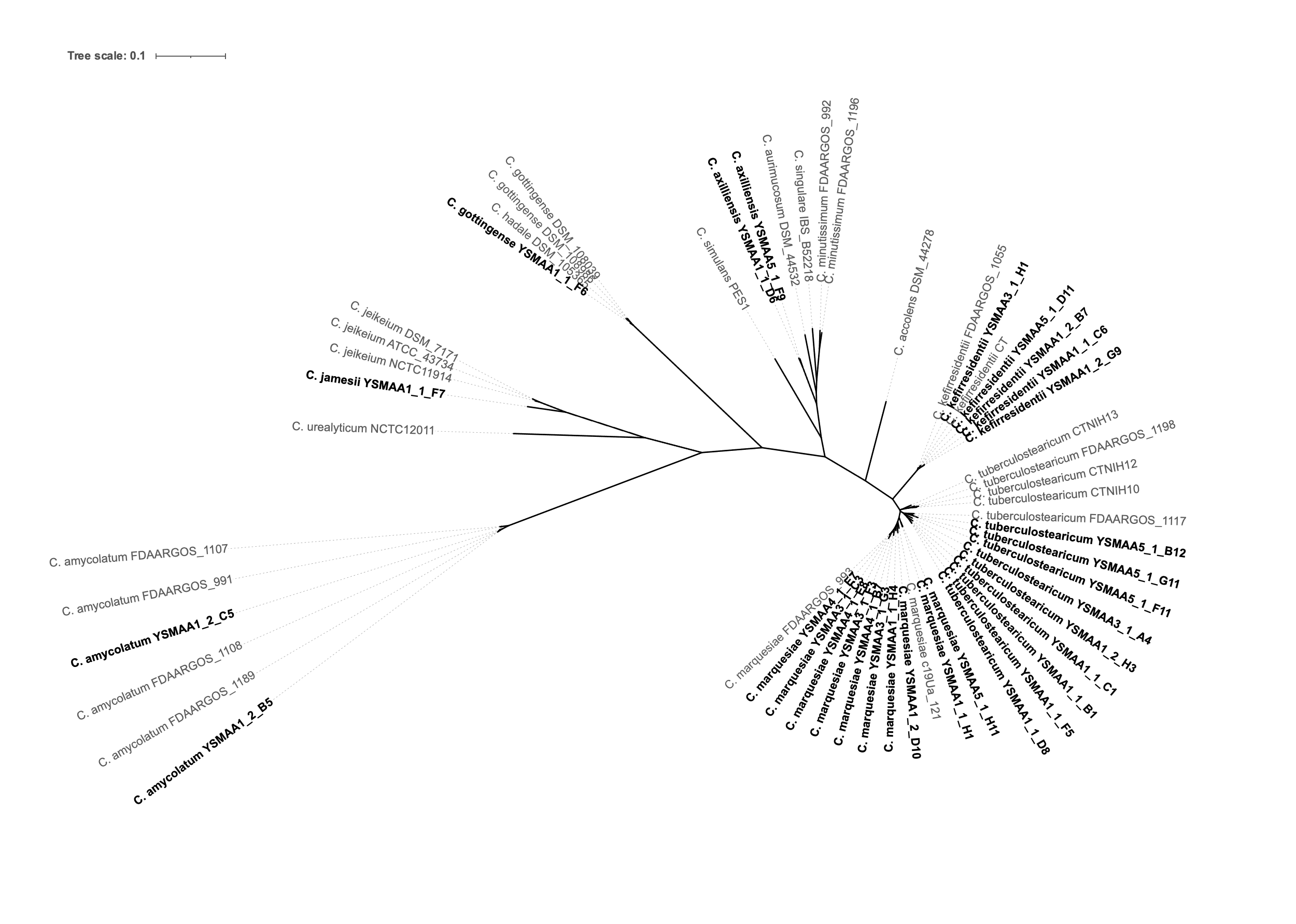

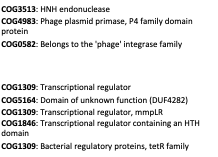

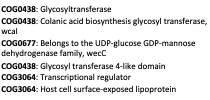

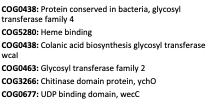

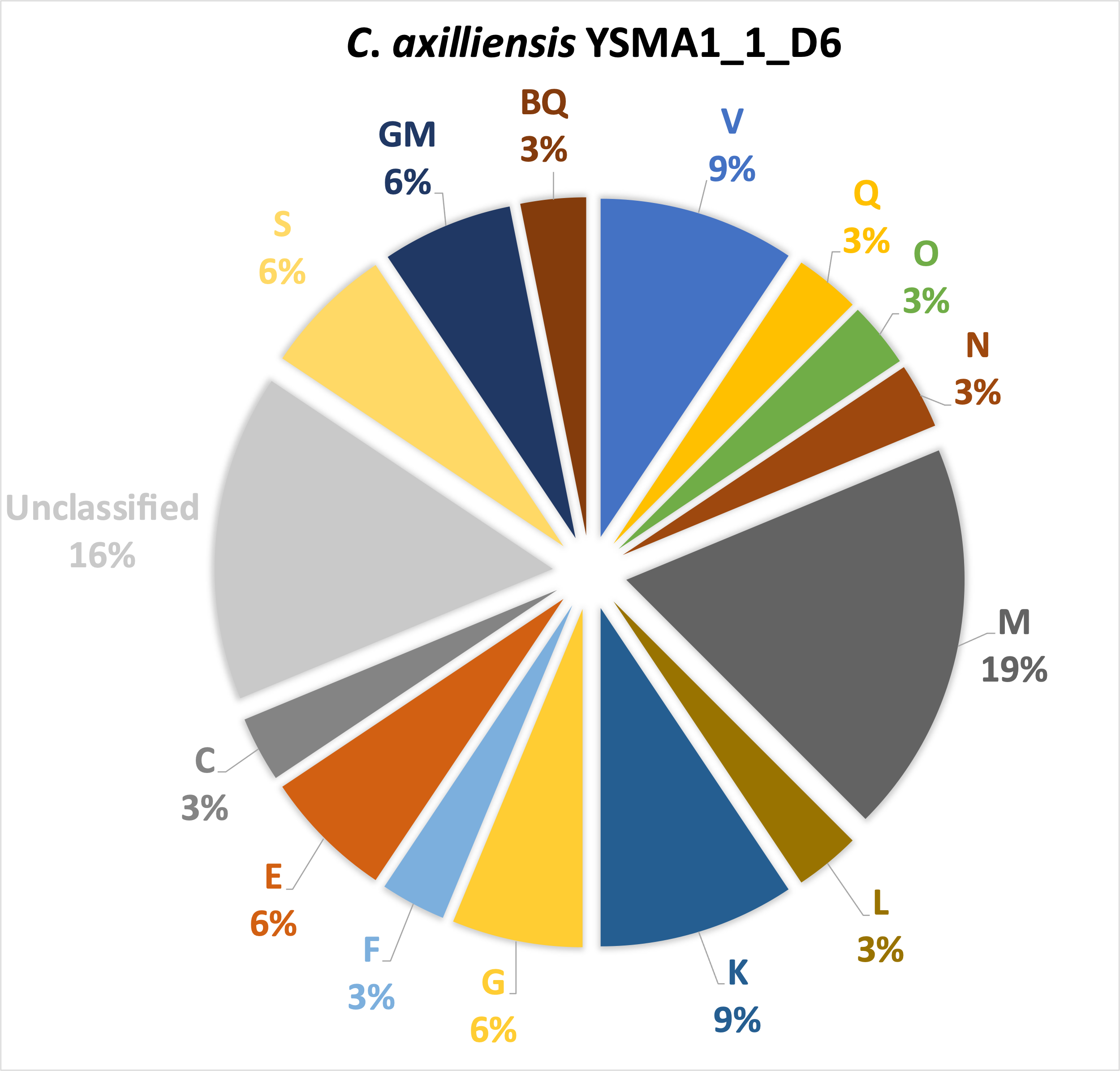

(32)

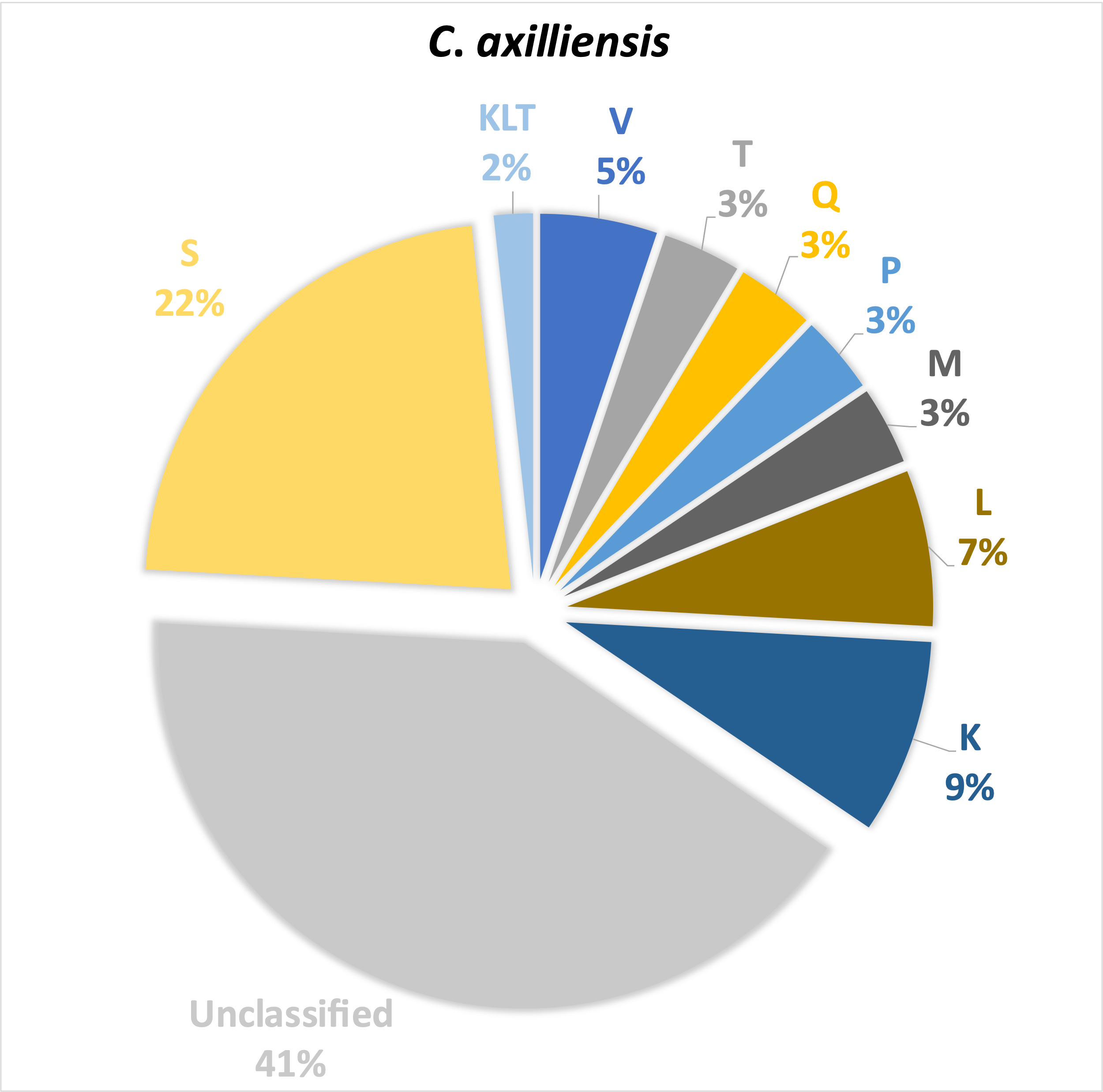

(59)

**A**

**C**

**L – Replication, recombination & repair**

**K – Transcription**

**M – Cell wall/membrane/envelope biogenesis**

**M – Cell wall/membrane/envelope biogenesis**

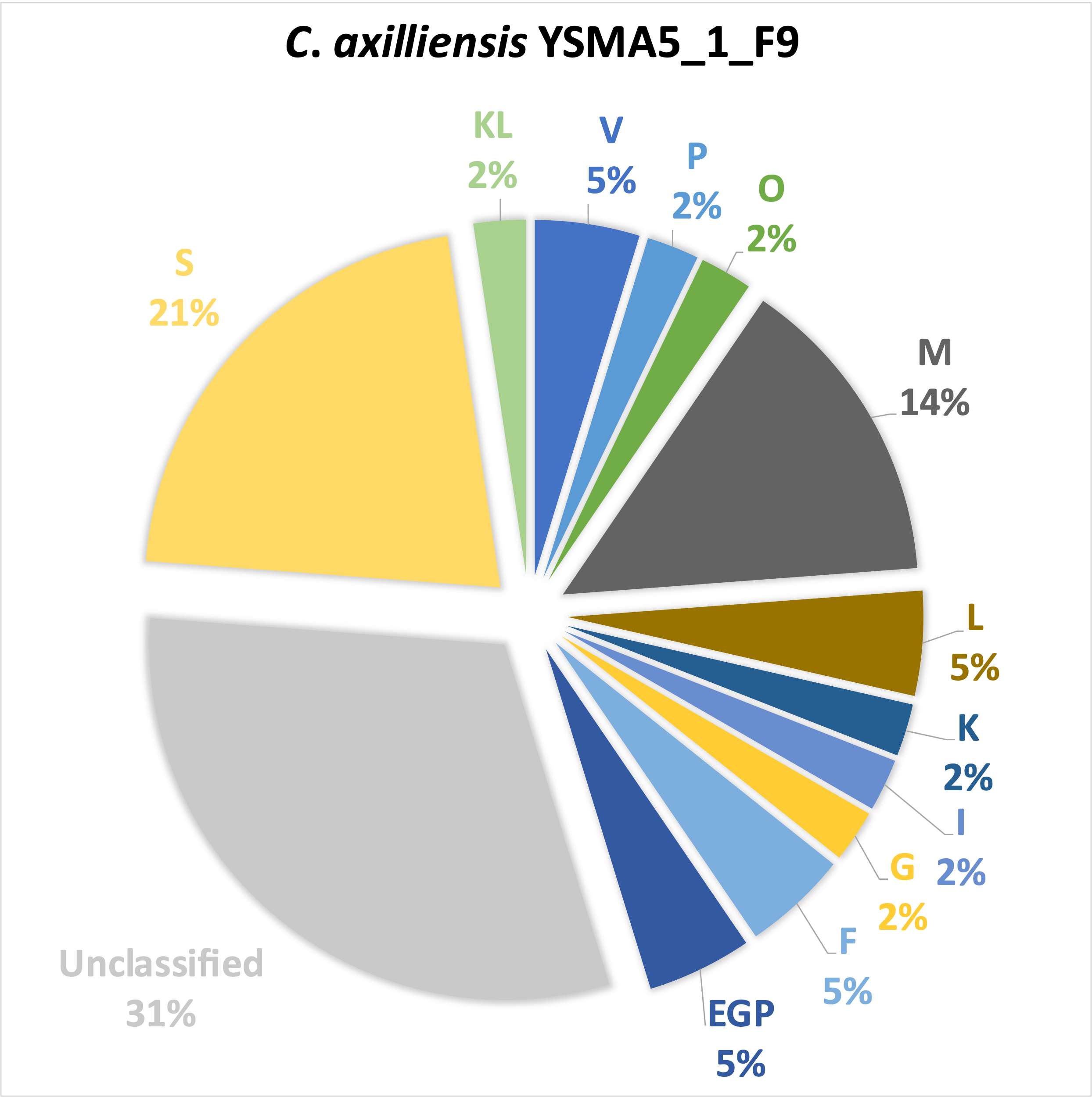

(42)

*C. axilliensis YSMAA5_1_F9*

*C. axilliensis YSMAA1_1_D6*

**B**

**Supplementary Figure 6. COG classification of genes of the *C. axilliensis*** (A) core genome and genes specific to (B) YSMAA5_1_F9 and (C) YSMAA1_1_D6. The percentage of genes within each COG classification is calculated against the total number of genes for each analysis. The genes of COG categories with the highest percentage and not unclassified or unknown (S) were shown.

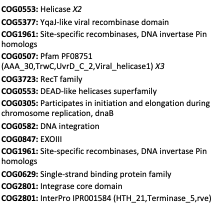

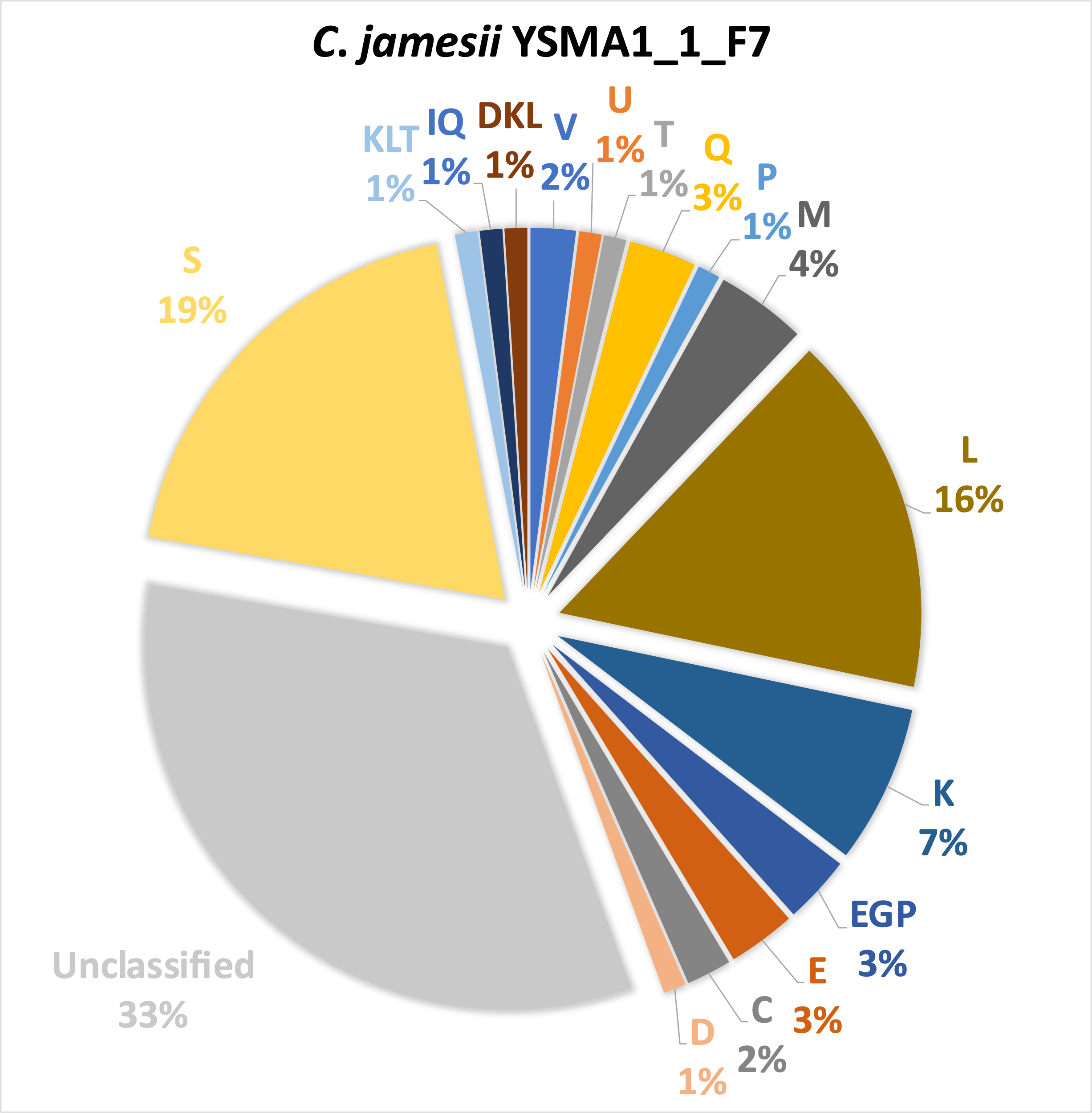

(100)

**L – Replication, recombination and repair**

**Supplementary Figure 7. COG classification of genes of the *C. jamesii* YSMAA1_1_F7.** The percentage of genes within each COG classification is calculated against the total number of genes for each analysis. The genes of COG categories with the highest percentage and not unclassified or unknown (S) were shown.

*C. jamesii YSMAA1_1_F7*

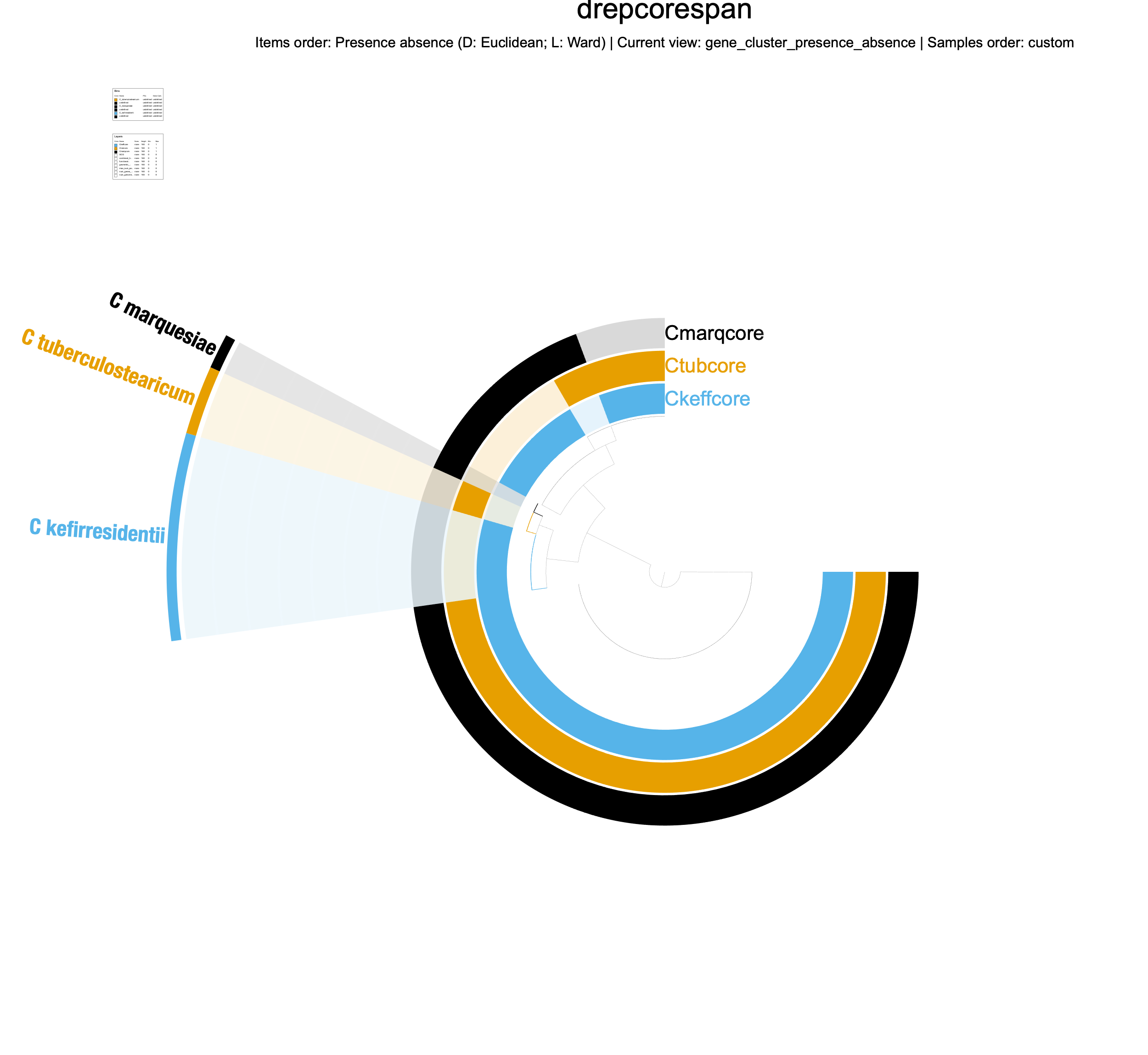

**Supplementary Figure 8. Comparisons of the core genomes of species *C. marquesiae*, *C. tuberculostearicum* and *C. kefirresidentii*** derived from the respective isolate genomes using anvi’o. Gene clusters specific to each species were highlighted using the coloured wedge.

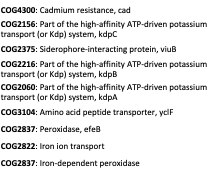

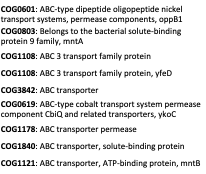

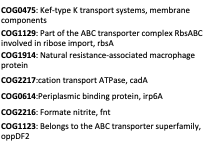

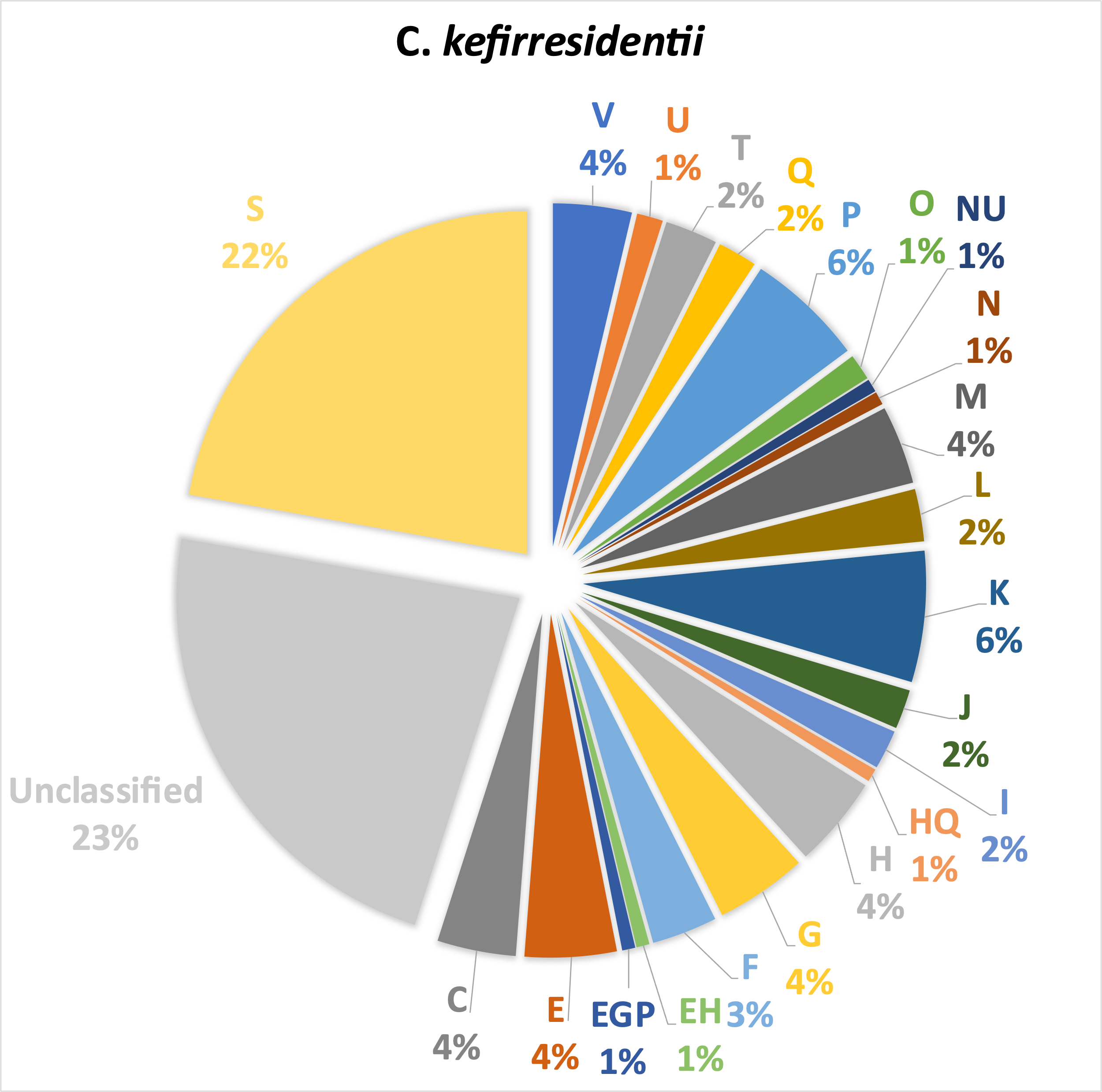

(162)

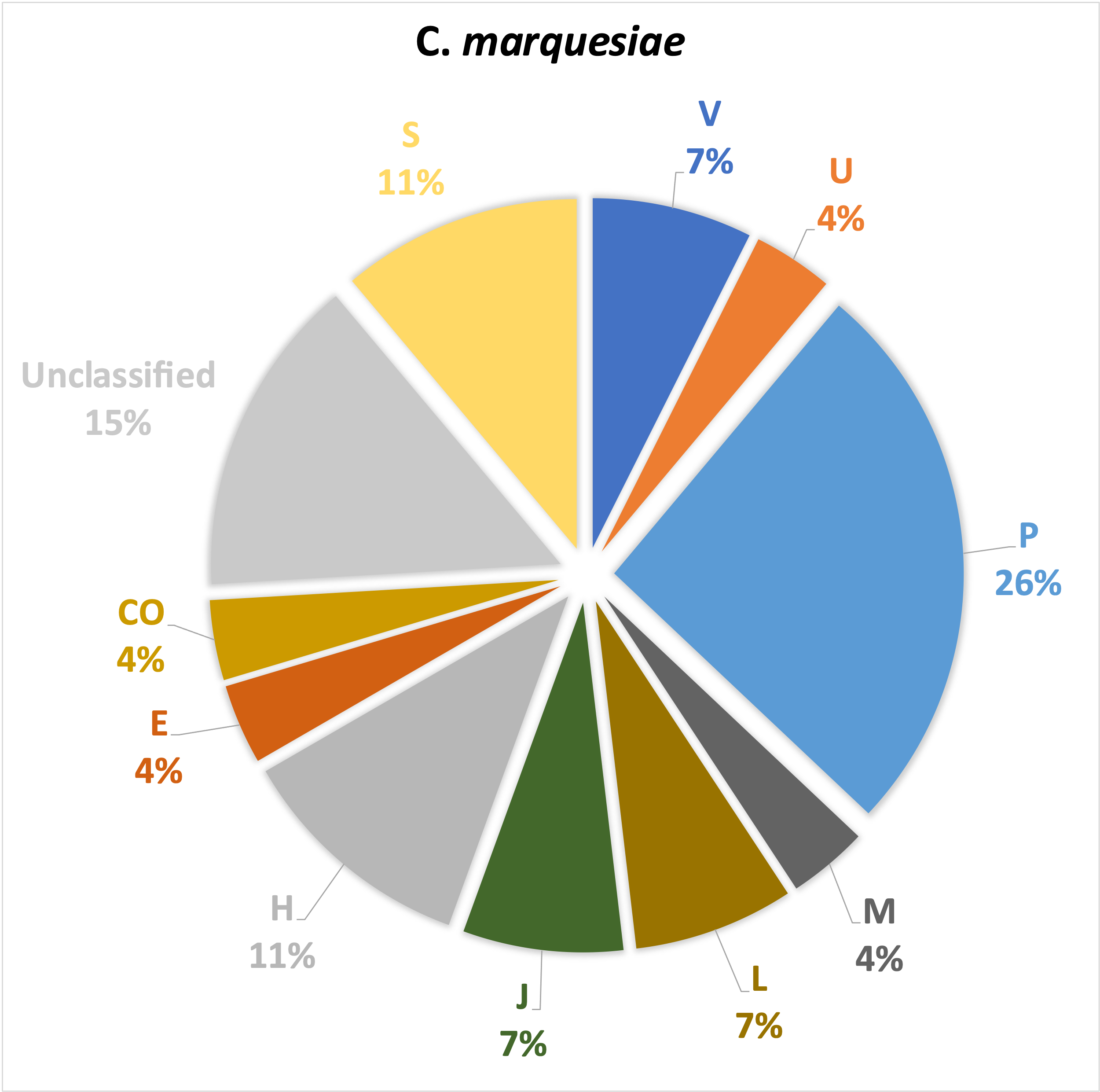

(27)

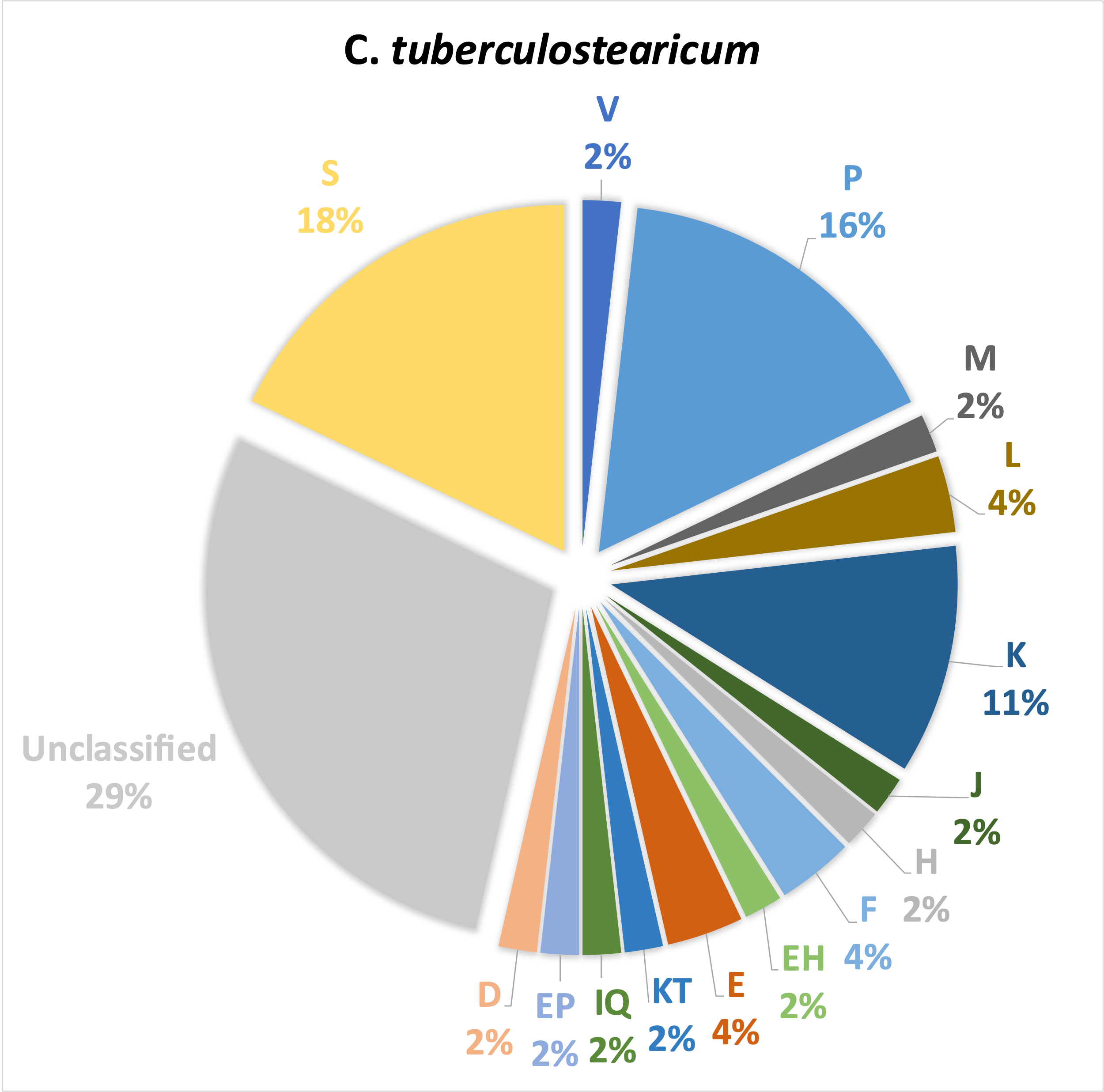

(56)

**A**

**B**

**C**

**P – Inorganic ion transport and metabolism**

**Supplementary Figure 9. COG classification of genes specific to (A) *C. kefirresidentii*, (B) *C. tuberculostearicum* and (C) *C. marquesiae*.** The percentage of genes within each COG classification is calculated against the total number of genes for each analysis. The genes of COG categories with the highest percentage and not unclassified or unknown (S) were shown.

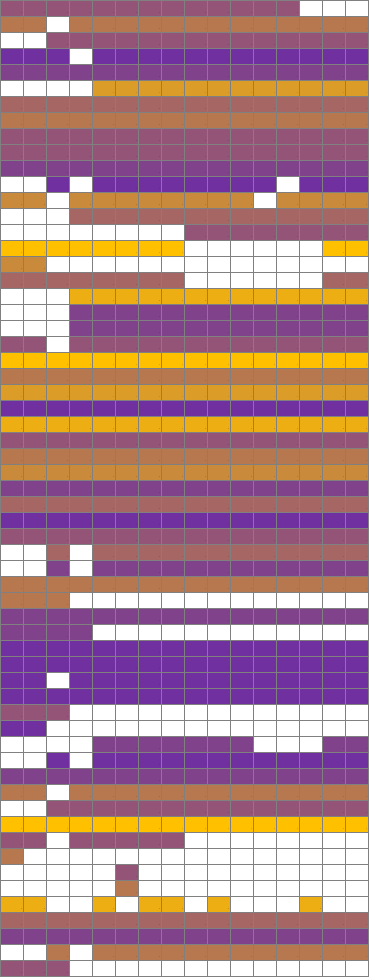

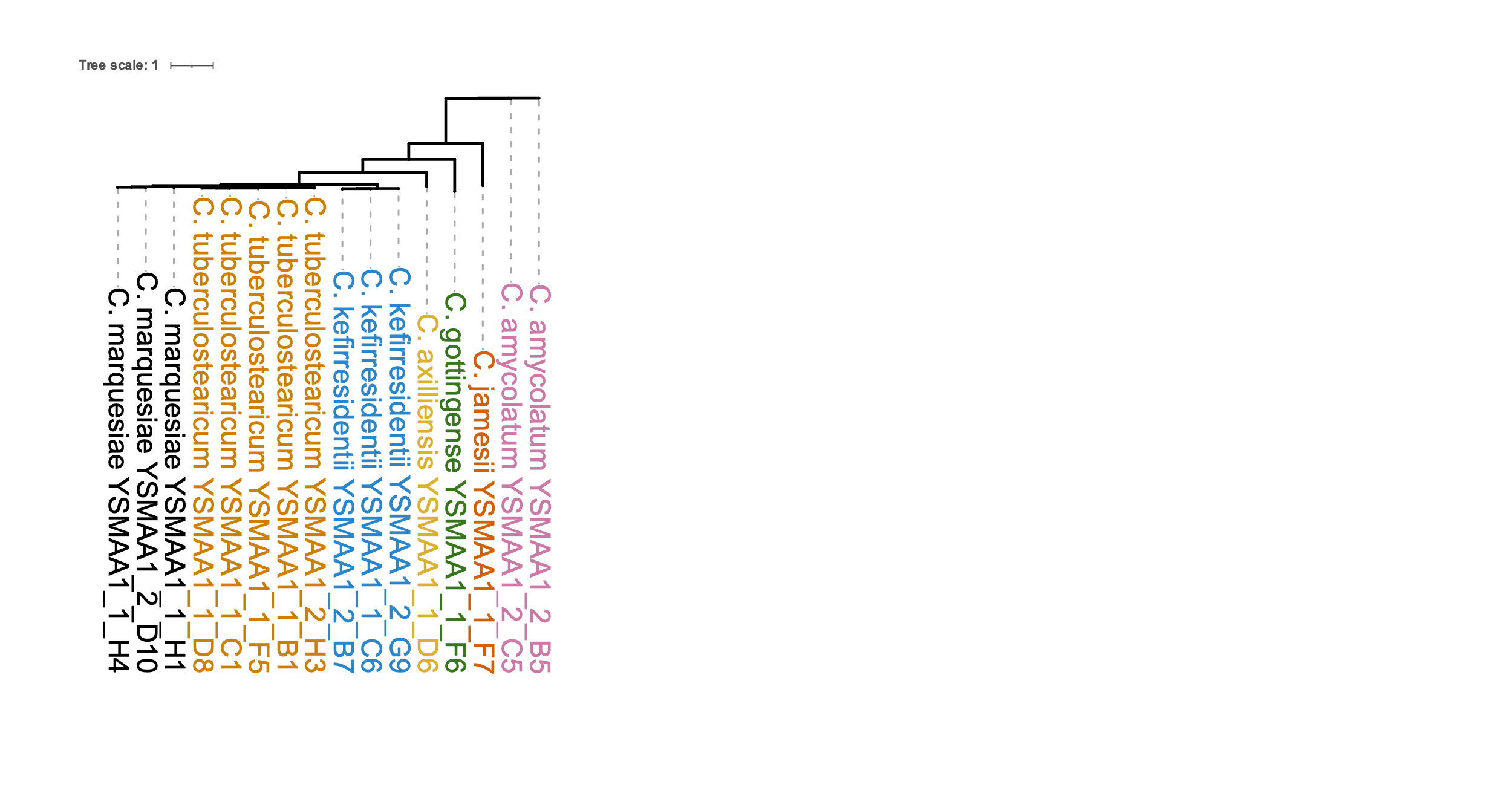

**Amino acid metabolism**

Biosynthesis of terpenoids and polyketides

Carbohydrate metabolism

Energy metabolism

**Lipid metabolism**

**Metabolism of cofactors and vitamins**

Nucleotide metabolism

M00020 Serine biosynthesis

M00018 Threonine biosynthesis

M00023 Tryptophan biosynthesis

M00016 Lysine biosynthesis

M00028 Ornithine biosynthesis

M00526 Lysine biosynthesis

M00083 Fatty acid biosynthesis

M00916 Pyridoxal-P biosynthesis

M00119 Pantothenate biosynthesis

M00123 Biotin biosynthesis

M00577 Biotin biosynthesis

**Supplementary Figure 10. Metabolic pathways in axillary corynebacteria of volunteer A1.** Each KEGG module for each isolate were placed on a heatmap where the modules with the largest number of reactions are in yellow and lowest in purple with absent modules in white. Modules of interest were described on the right of the heatmap. KEGG modules were grouped according to the categories on the left of the heatmap. Isolates were grouped by species with a core genome tree depicting the variation.

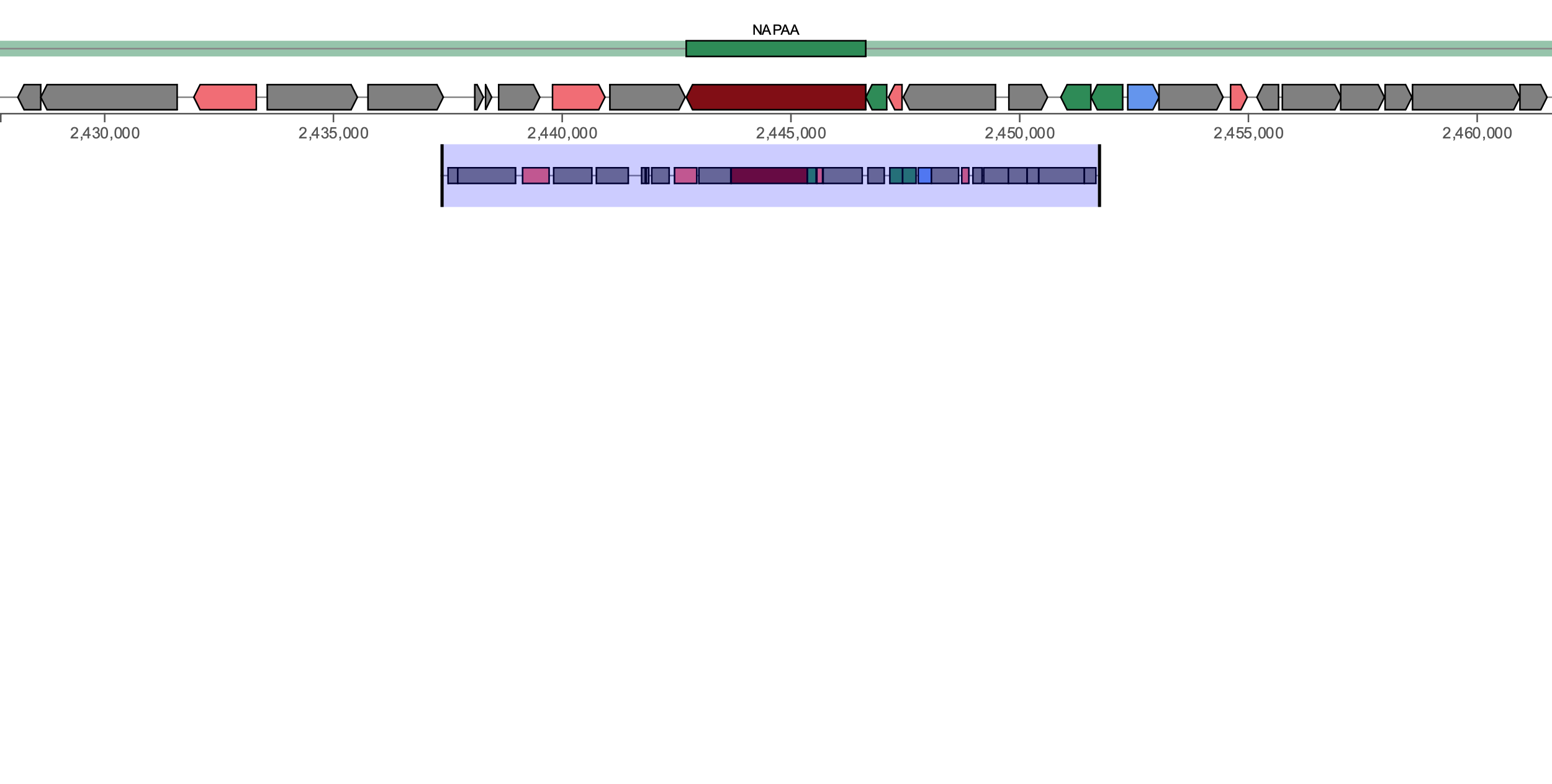

NAPAA

***C. tuberculosteariucm***

**YSMAA1_1_C1**

(cyclofaulknamycin, 16%)

dapE

**dltA**

ytrE

degS

lnrK

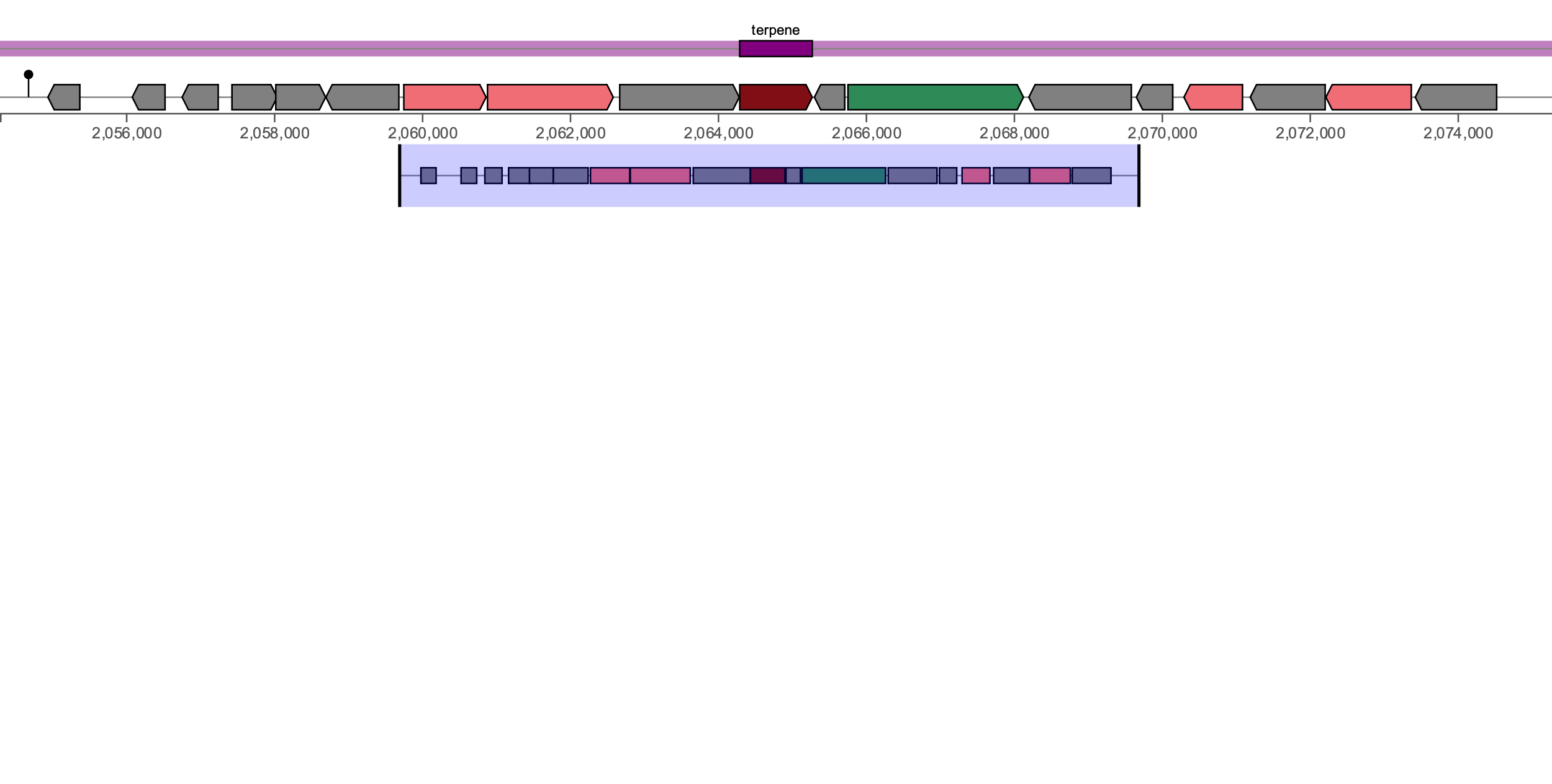

Terpene (4)

***C. amycolatum***

**YSMAA1_2_B5**

(no similar cluster)

pknD

**crtB**

carA2

pimB

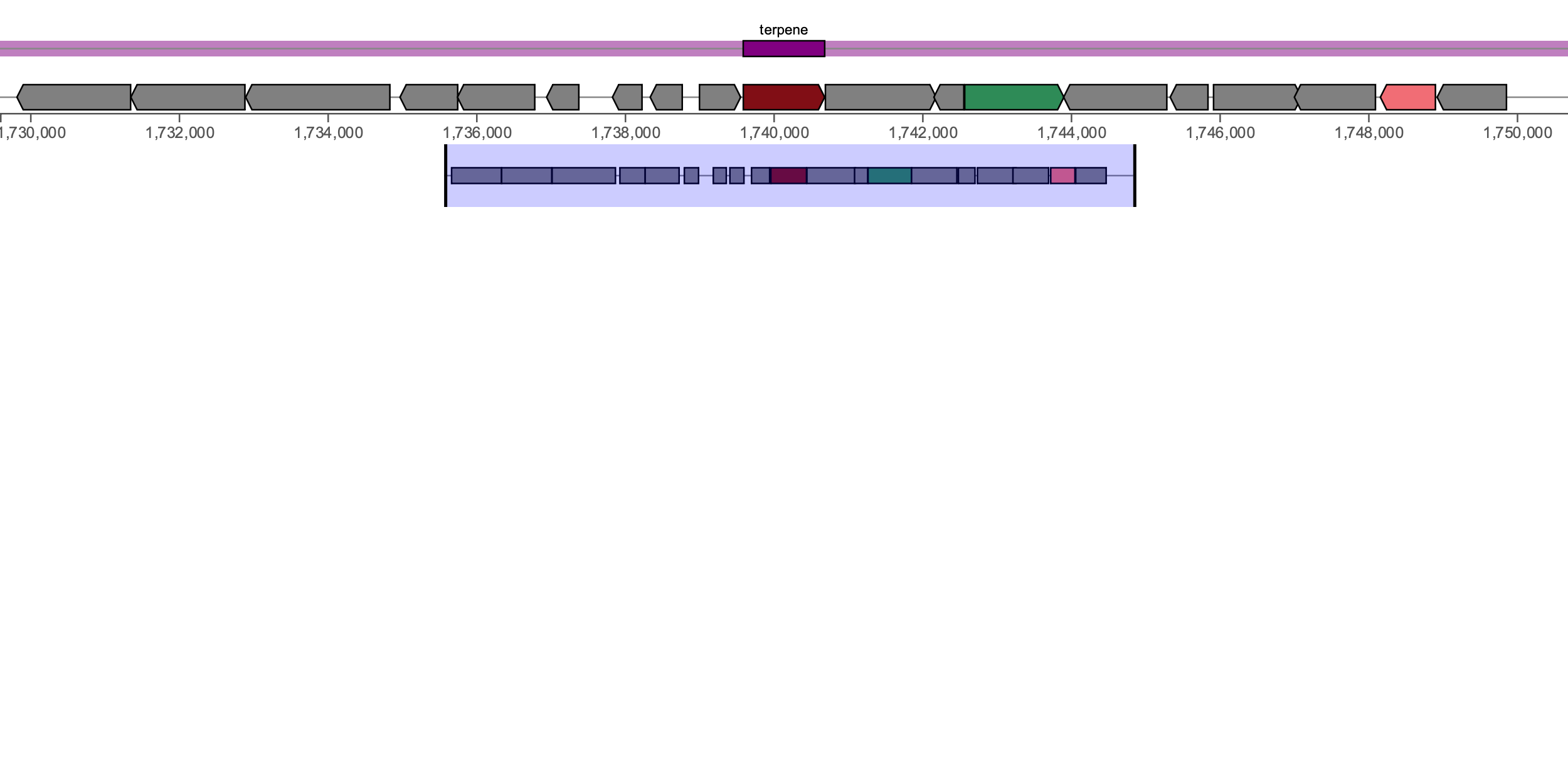

Terpene (6)

***C. kefirresidentii***

**YSMAA1_1_C6**

(no similar cluster)

pknD

**napT7**

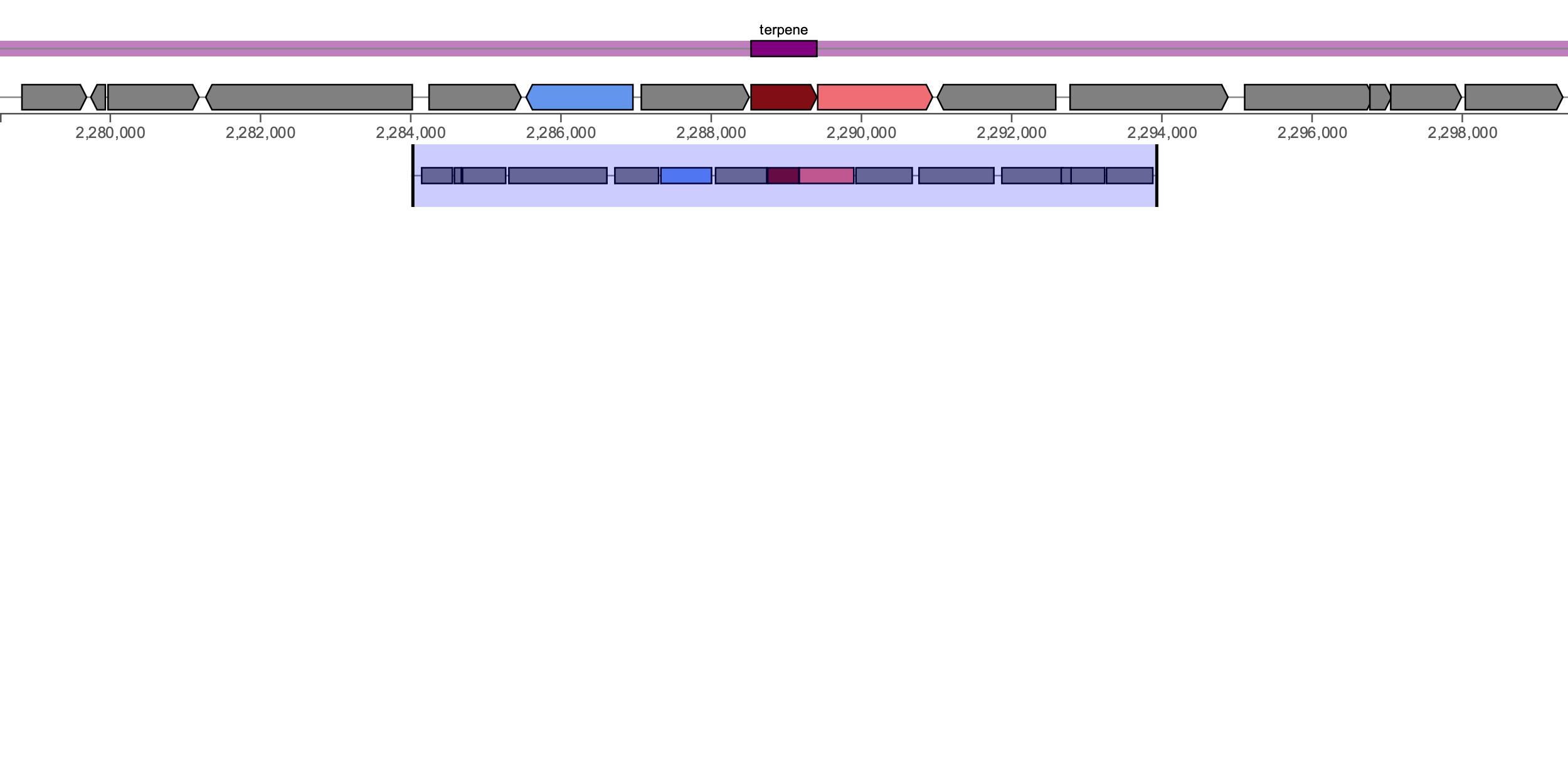

Terpene (3)

***C. kefirresidentii***

**YSMAA1_1_C6**

(carotenoid, 25%)

**crtN**

lysS1

ribZ

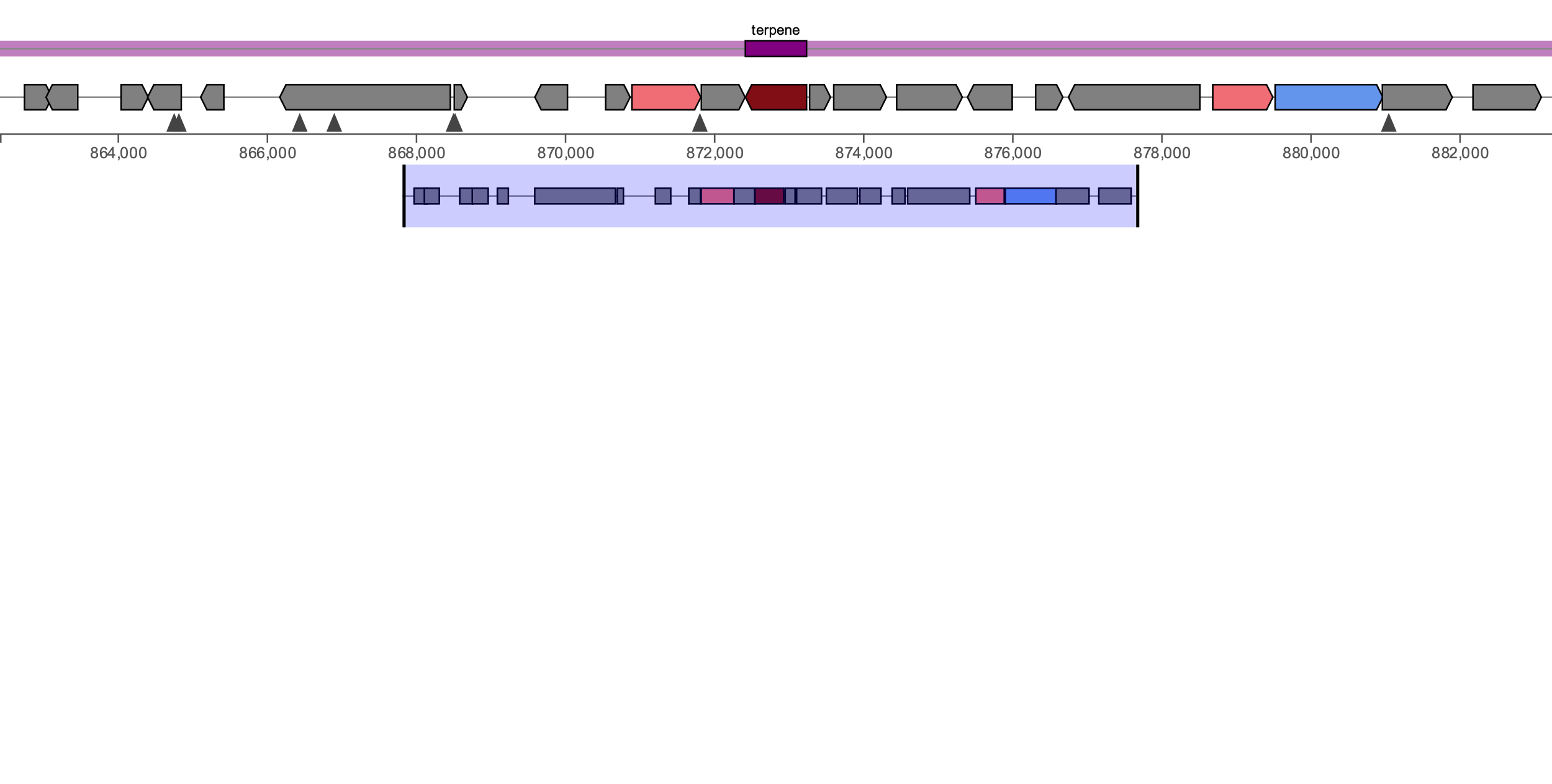

Terpene (1)

***C. gottingense***

**YSMAA1_1_F6**

(oxalomycin B, 6%)

**crtB**

arcB

qacA

bla

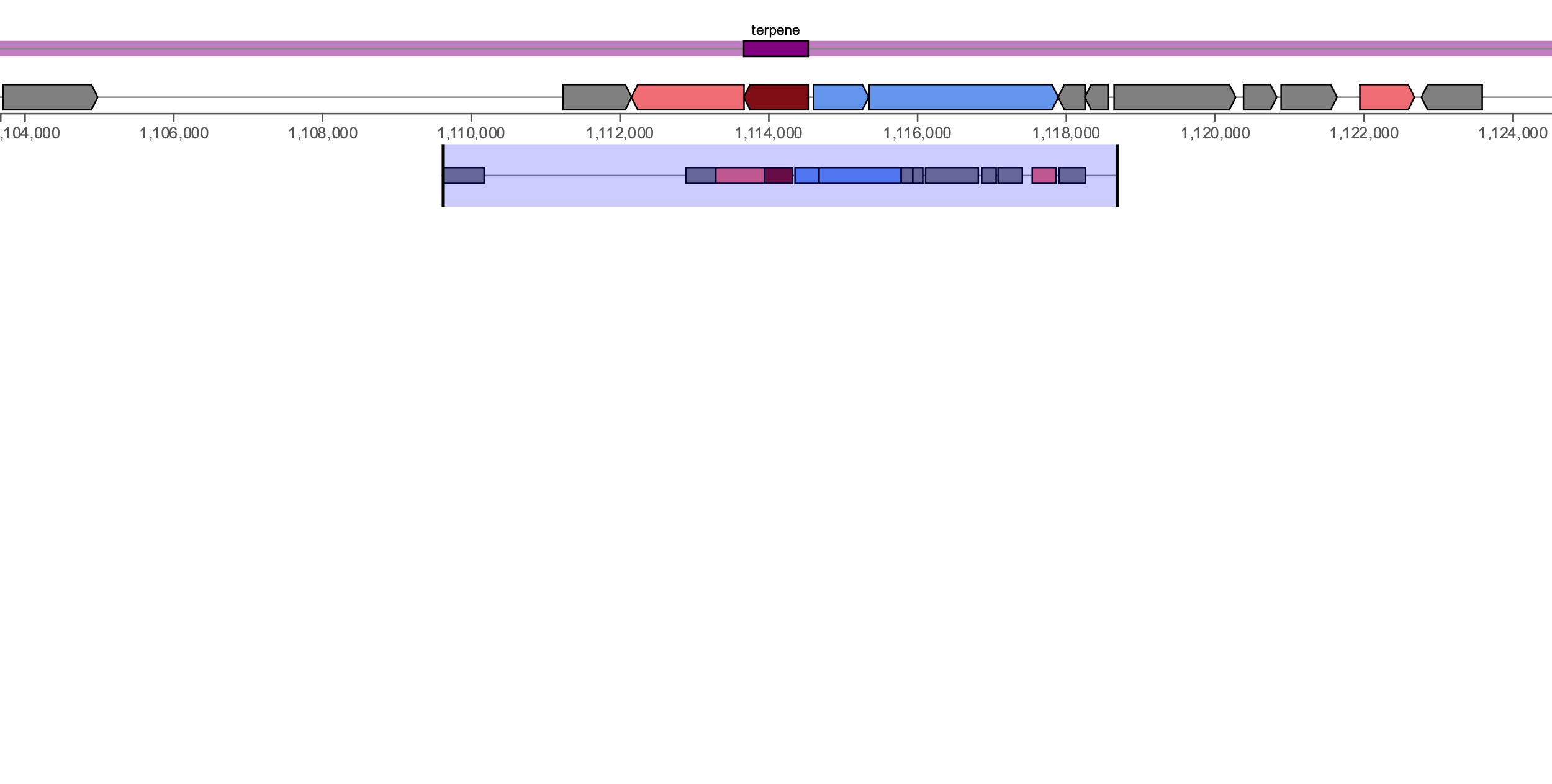

Terpene (2)

***C. axilliensis***

**YSMAA1_1_D6**

(carotenoid, 25%)

crtN

ABC Transporter

**crtB**

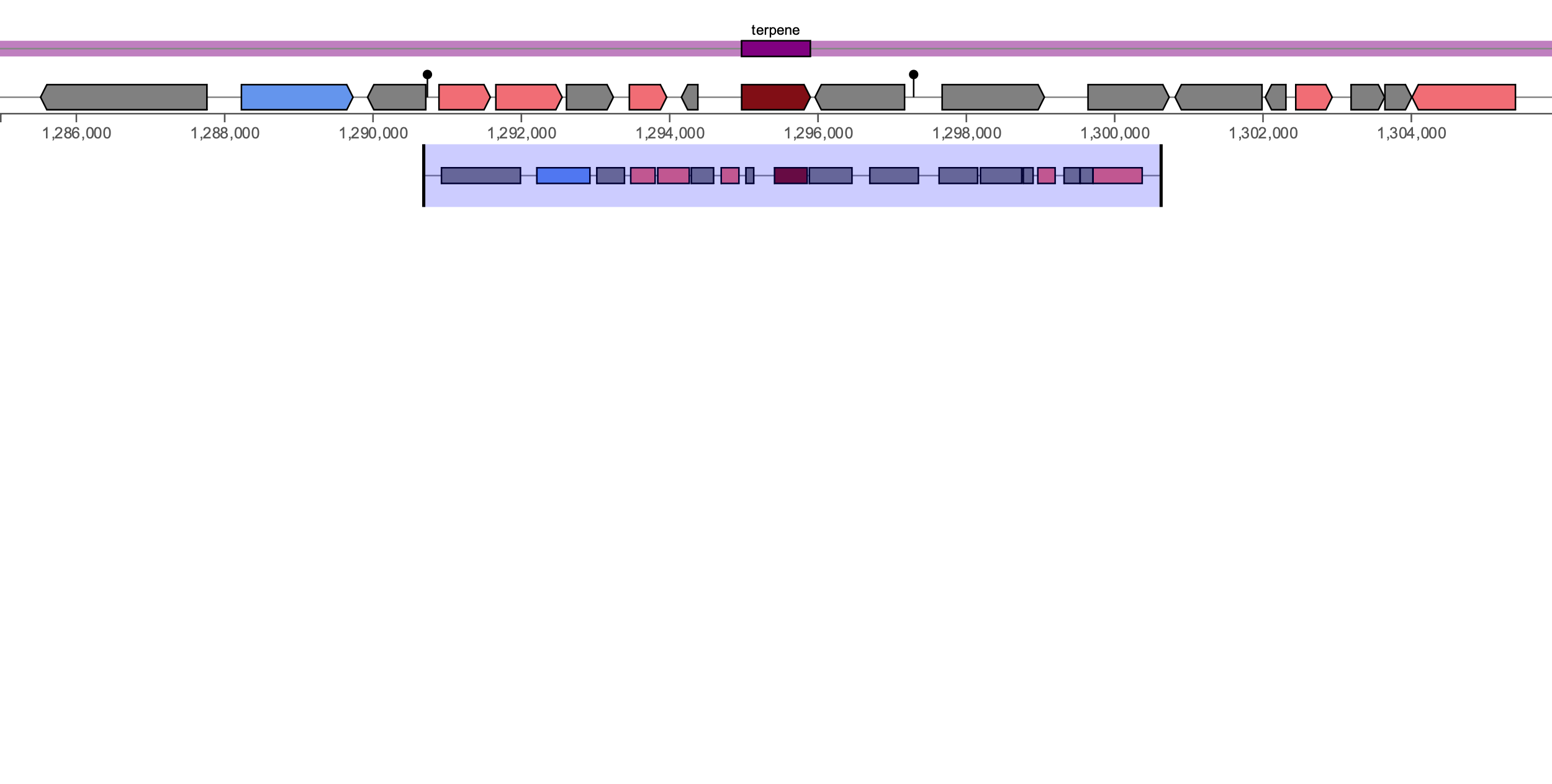

Terpene (5)

***C. jamesii***

**YSMAA1_1_F7**

(no similar cluster)

**crtB**

npcB

sdr

abi

bcp

ybiT

amiC

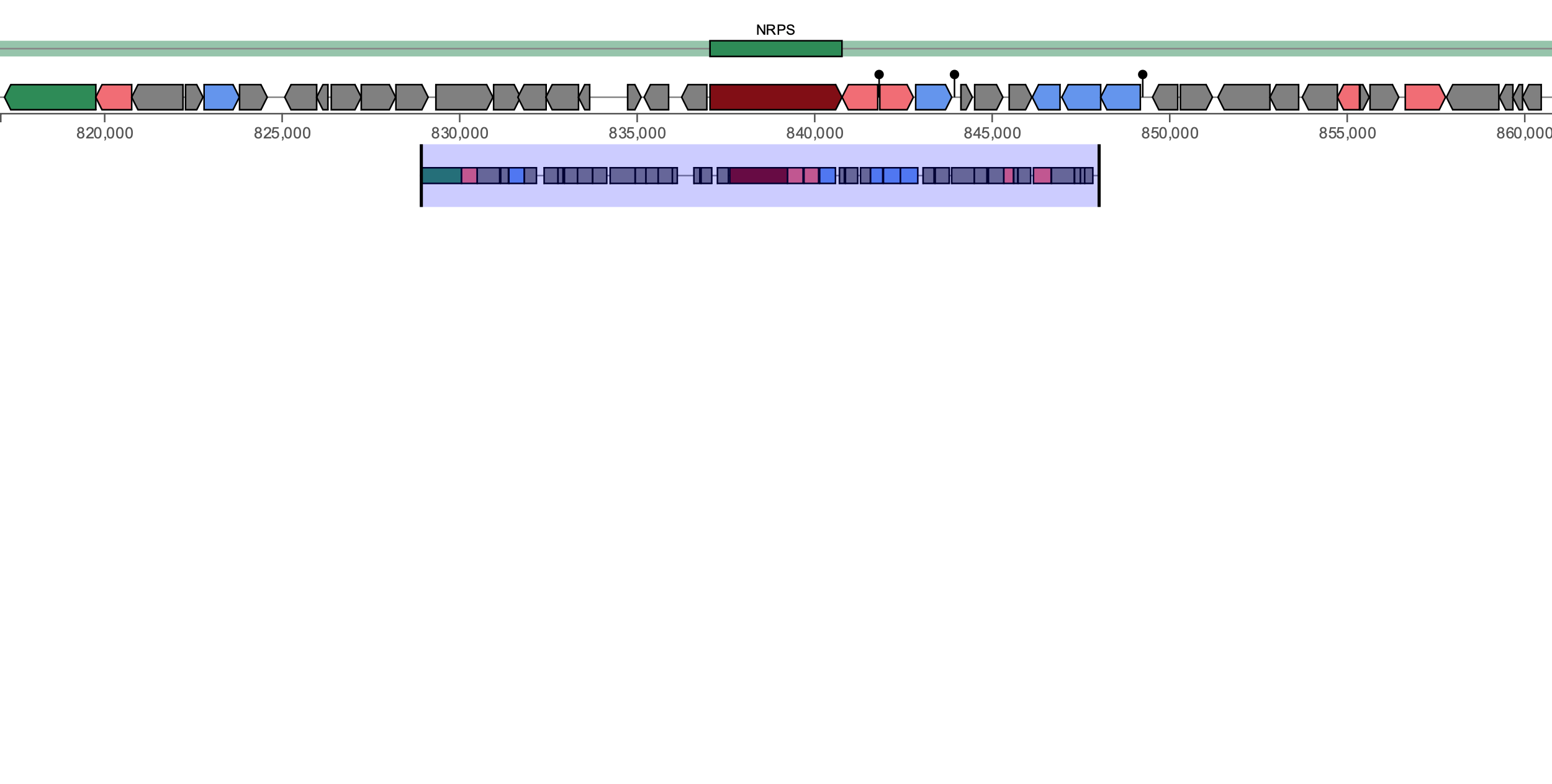

NRPS (1)

***C. tuberculostearicum***

**YSMAA5_1_B12**

(coelichelin, 36%)

pknD

glnH

bcrA

**igrD**

yfmC

fmt

ABC Transporter

fes

def

ybdK

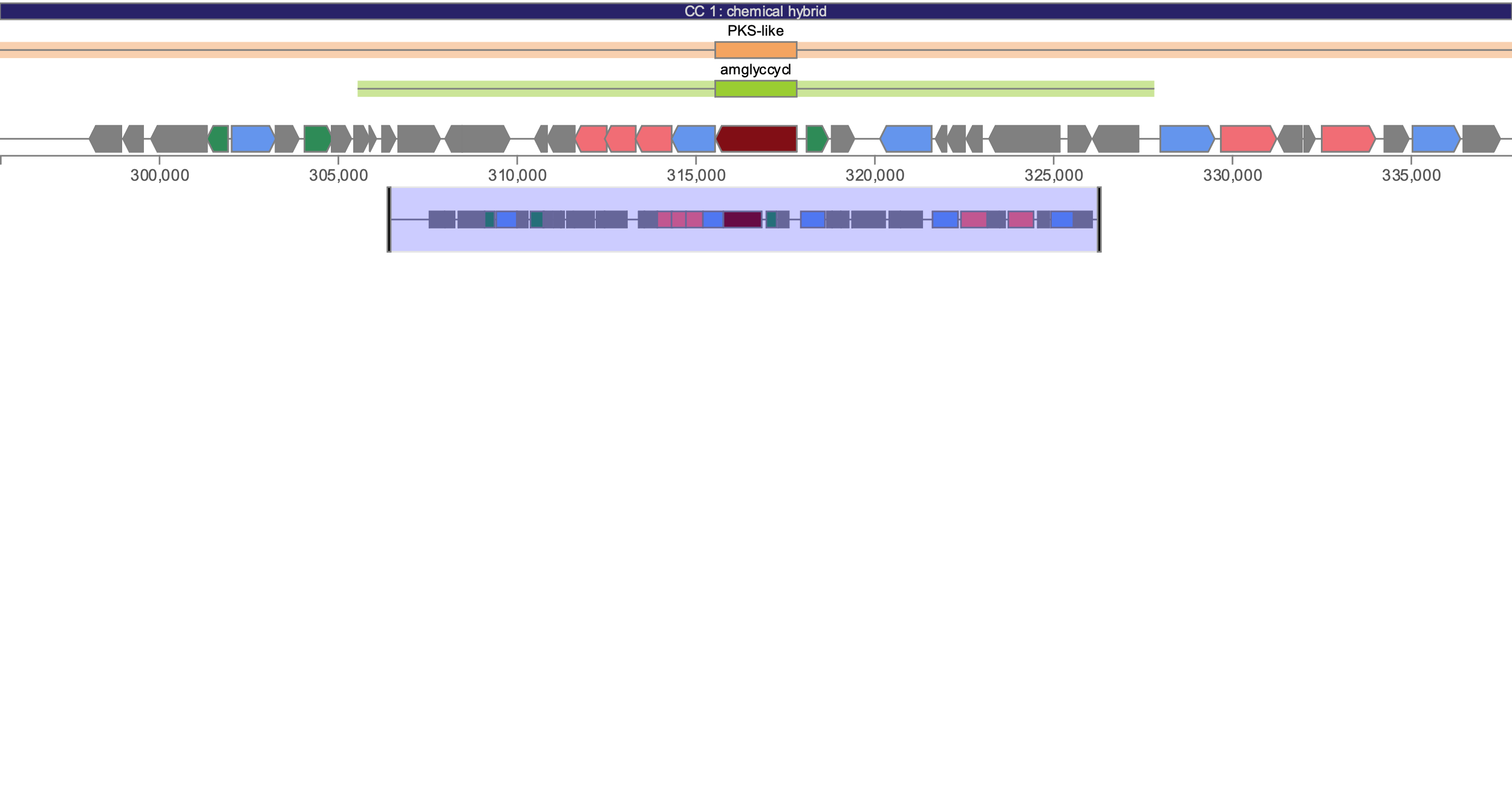

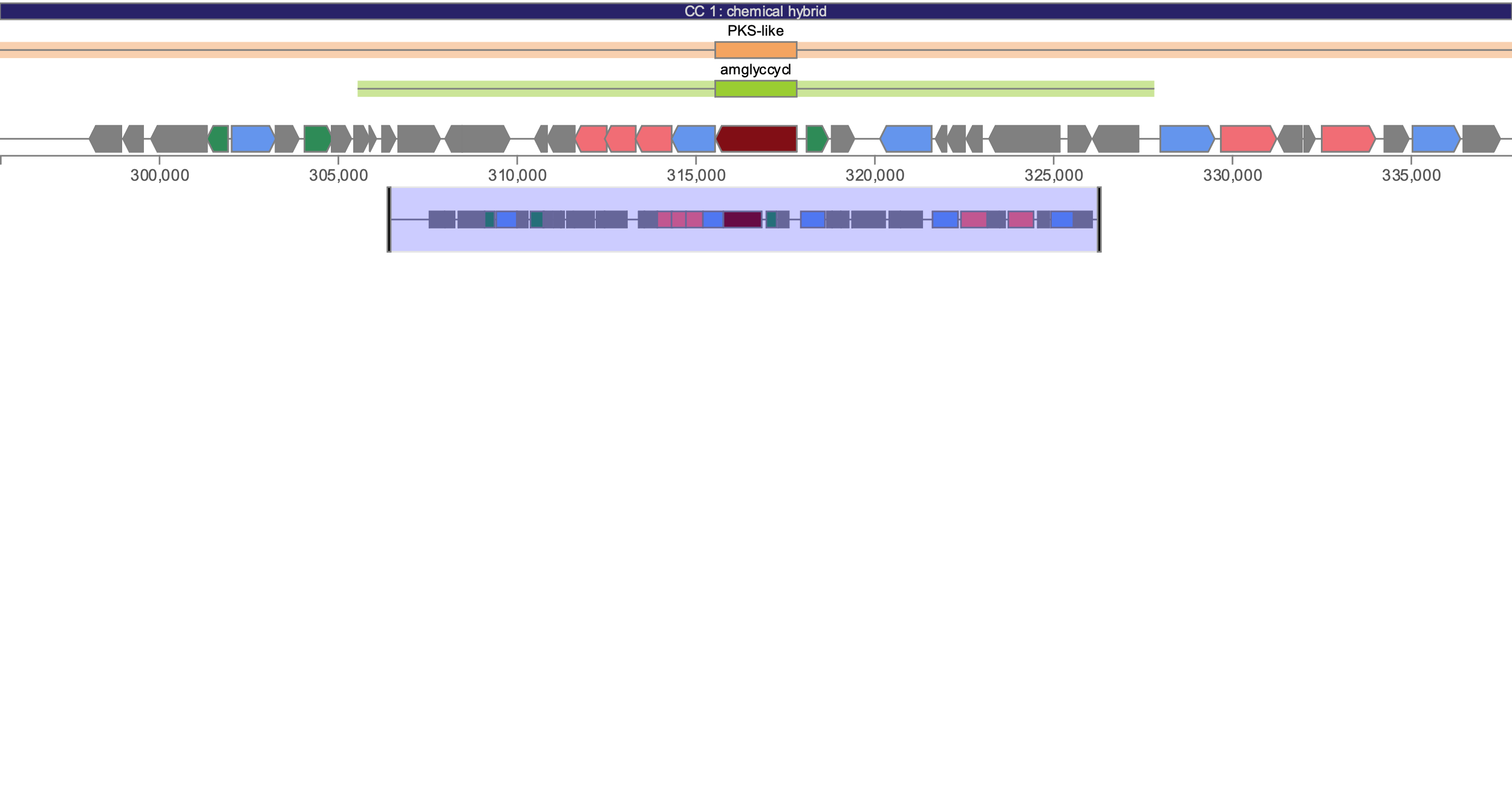

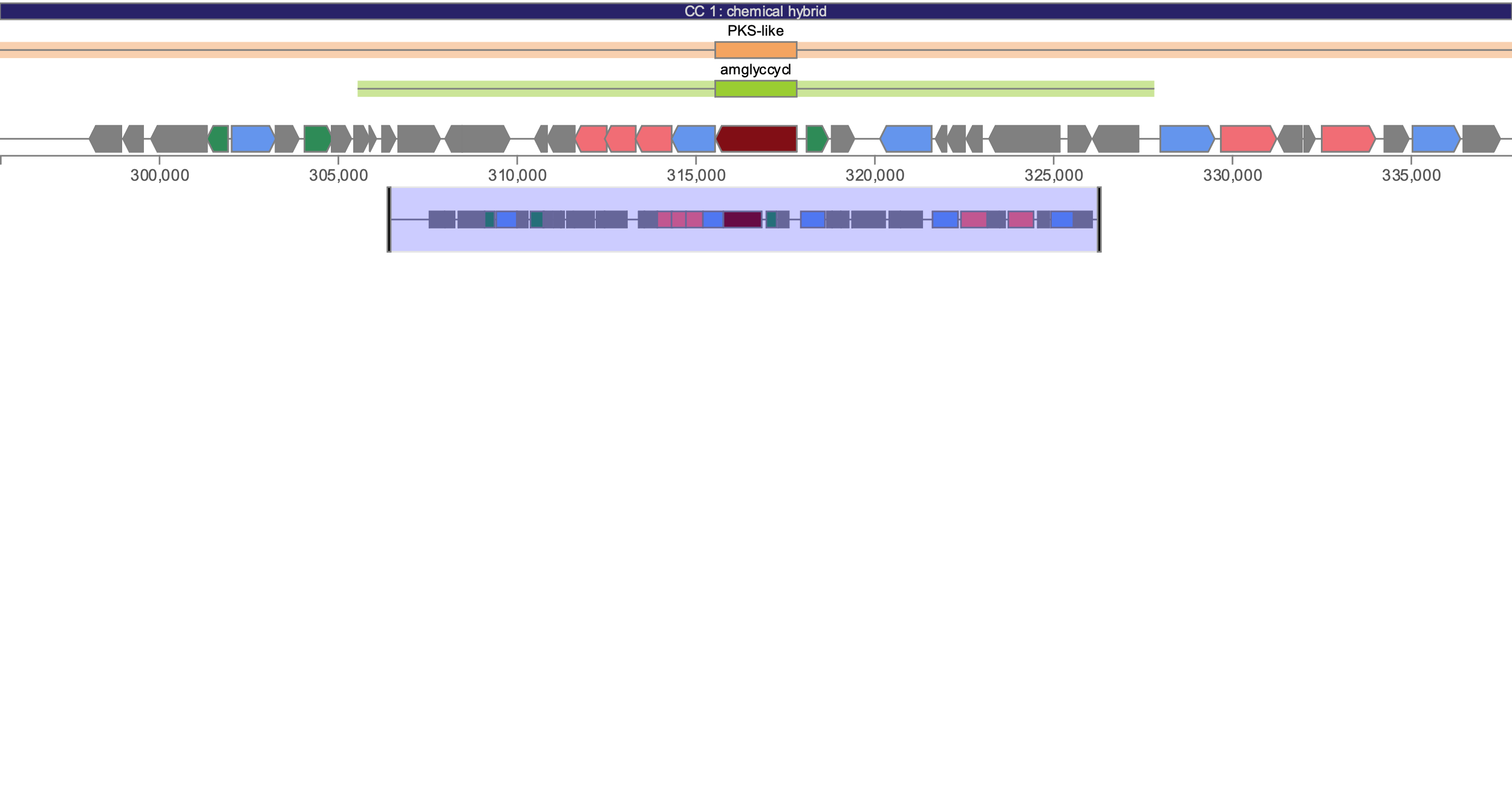

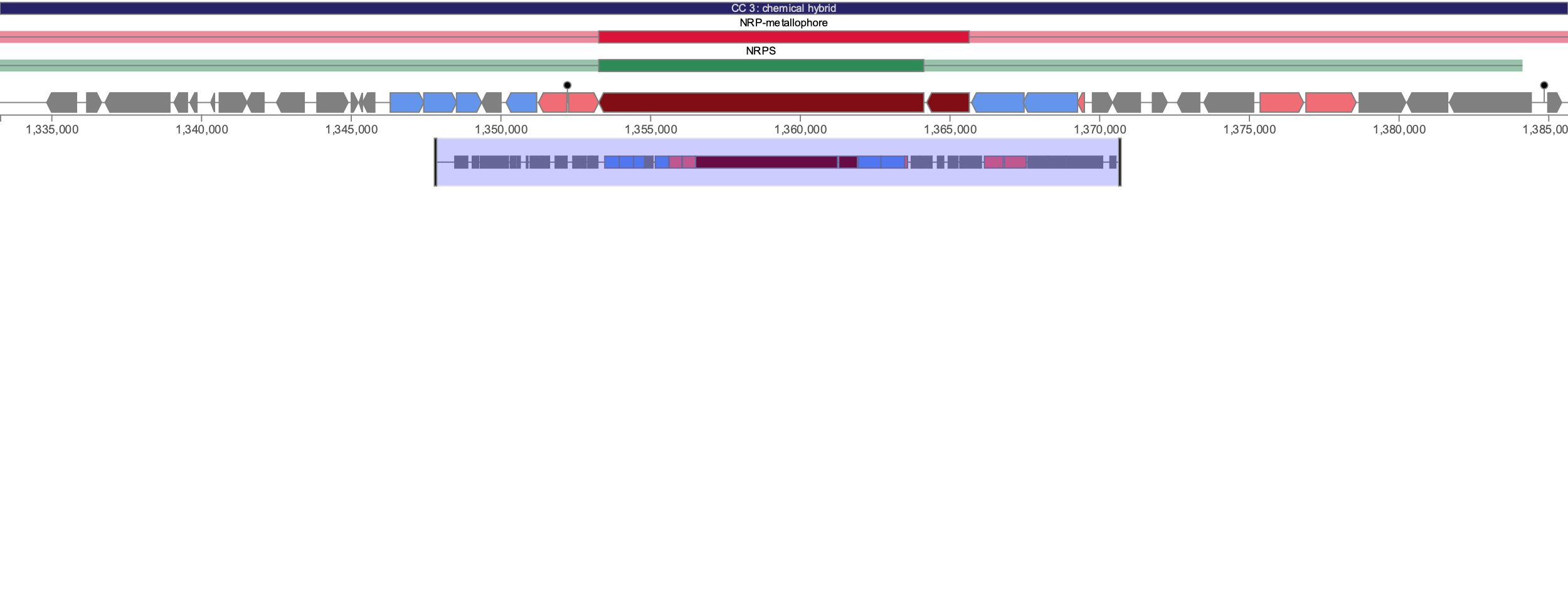

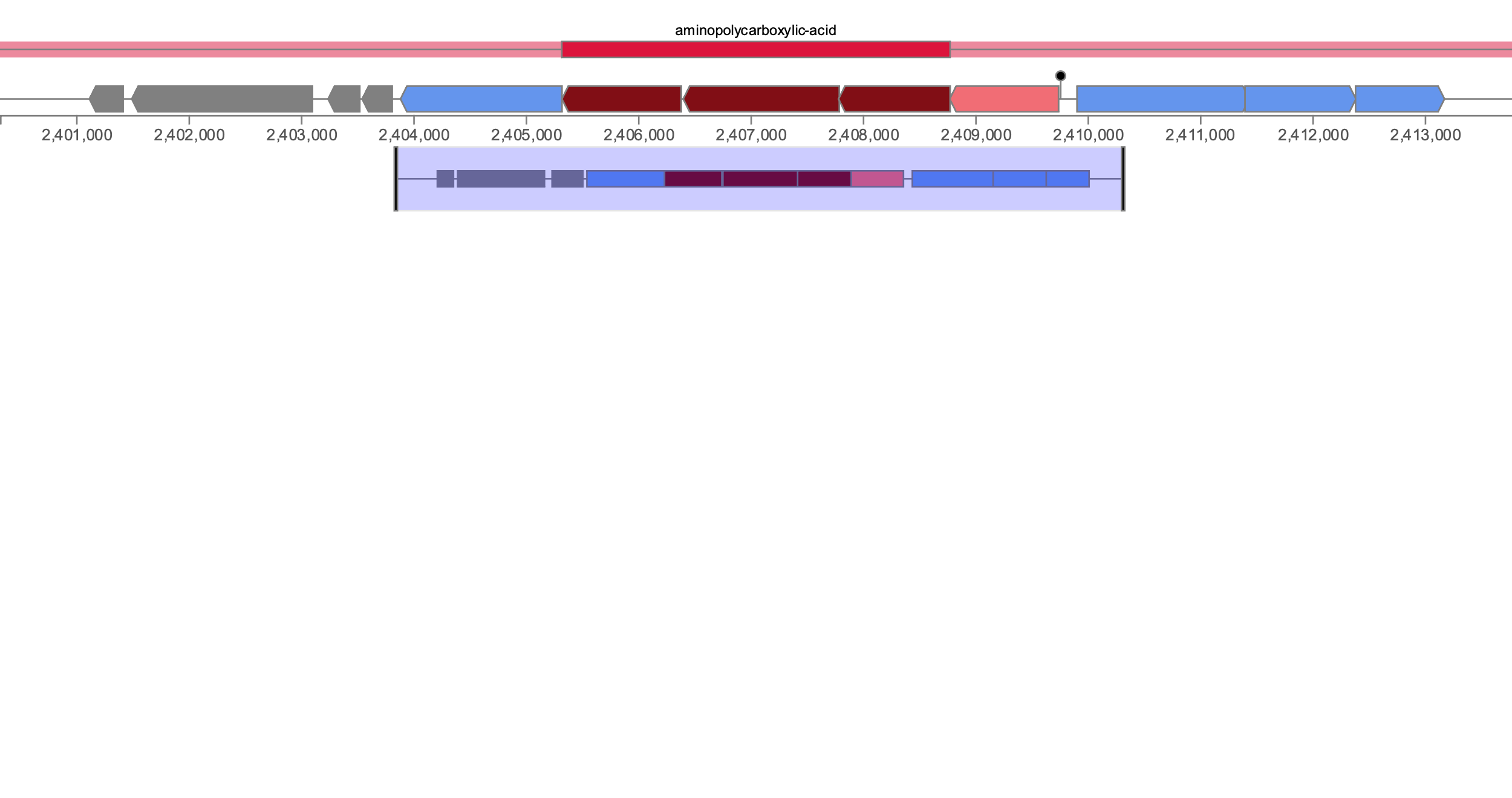

Other genes

Core biosynthetic

Regulatory

Transport

Additional biosynthetic

Legend:

**Supplementary Figure 11**. Architectures of the 8 distinct putative biosynthetic gene clusters predicted using antiSMASH 7.

NRPS-like

***C. tuberculostearicum***

**YSMAA5_1_F11**

(HTTPCA/picibactin/

prepicibactin, 4%)

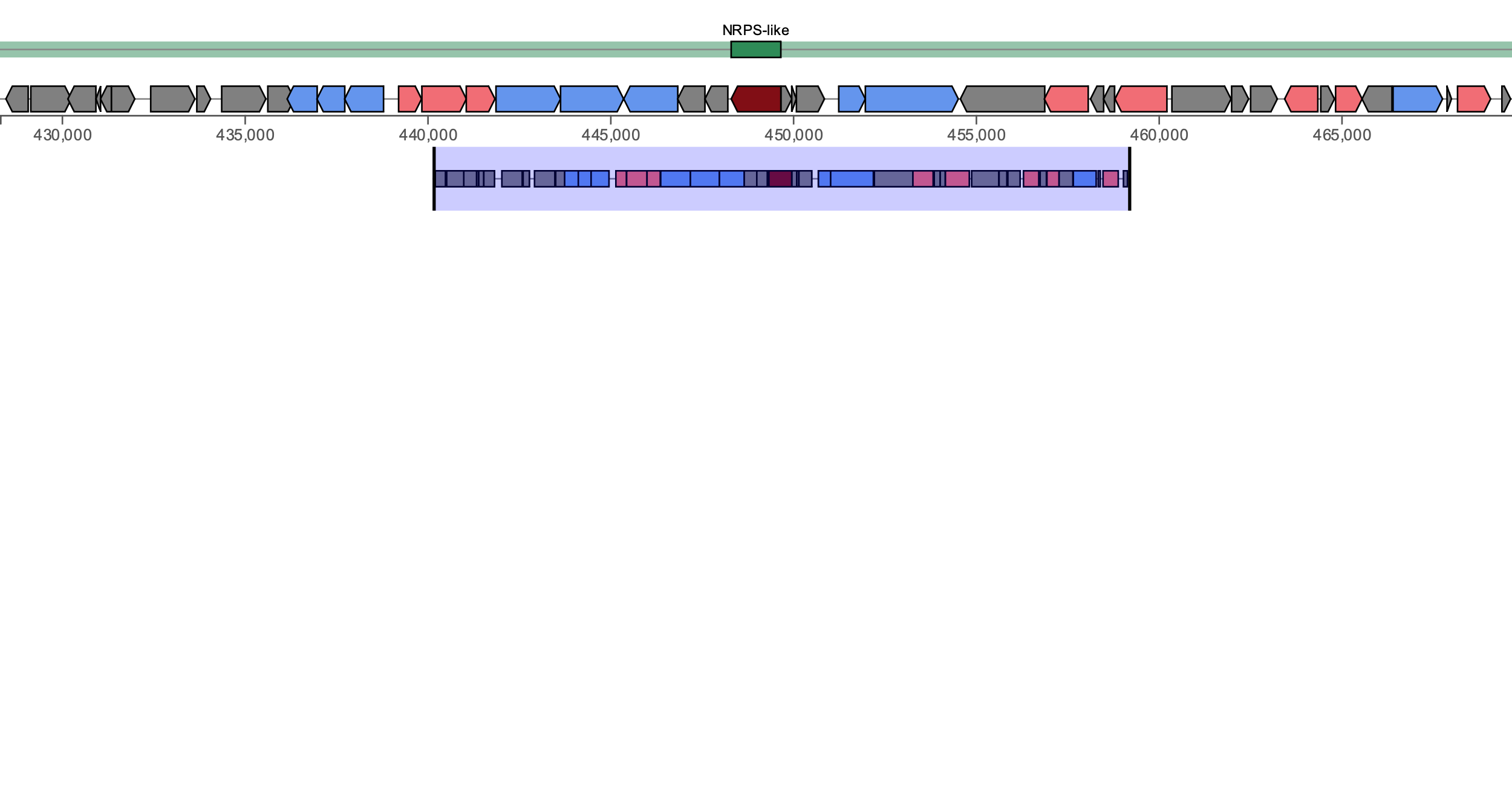

**angR**

Mac

tylN

pikAV

ABC-2 Transporter

ABC Transporter

iolT

alsT

trxB

aph

ABC Transporter

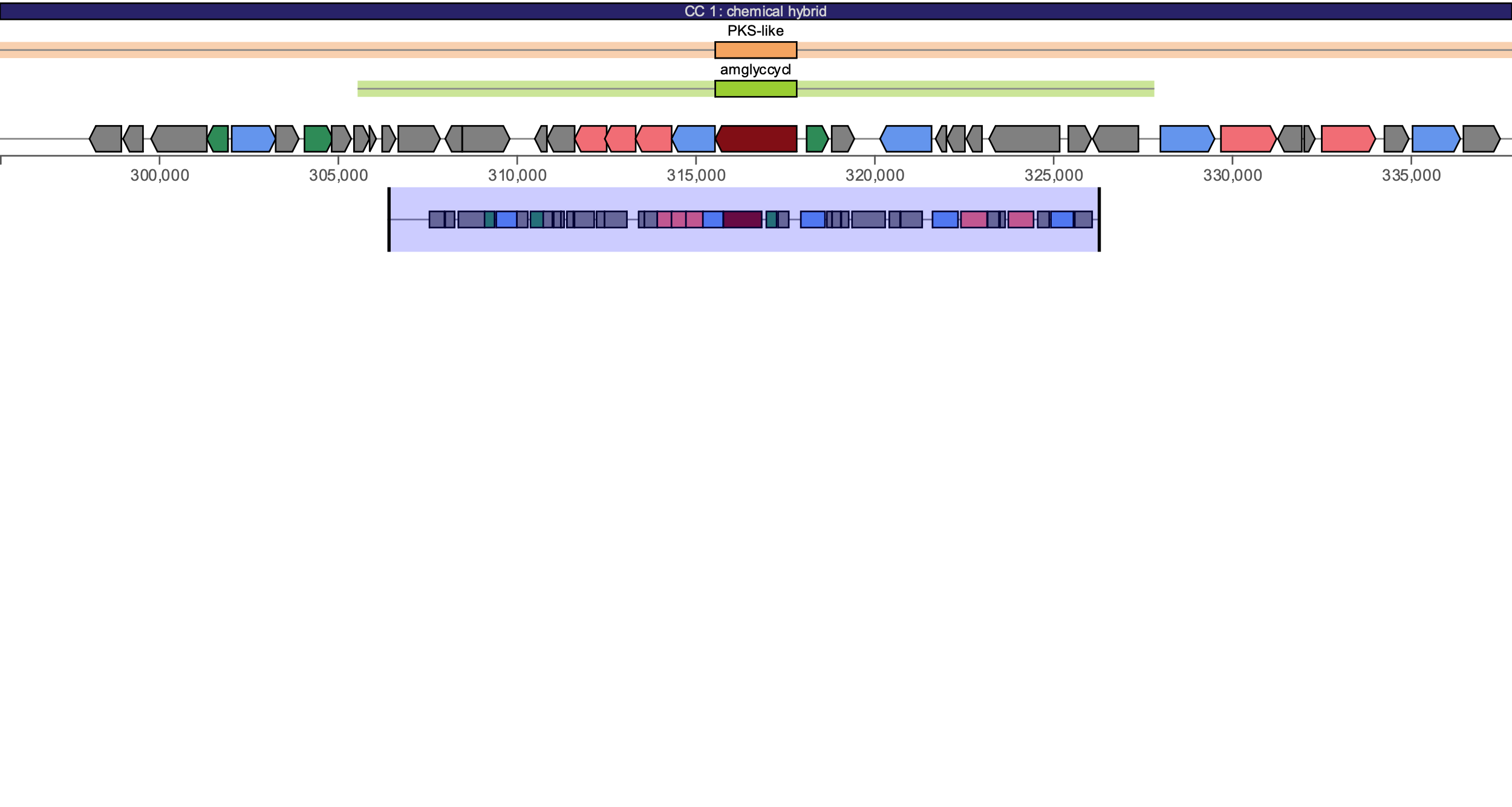

**salQ/**

**ksIII**

comR

ydhP

lgoD

ycdF

tdh

ynfM

dctA

ilvA

gatA

ribZ

emrB

PKS-like, aminoglycoside

***C. jamesii***

**YSMAA1_1_F7**

(no similar cluster)

NRP-metallophore

***C. jamesii***

**YSMAA1_1_F7**

(coelichelin, 45%)

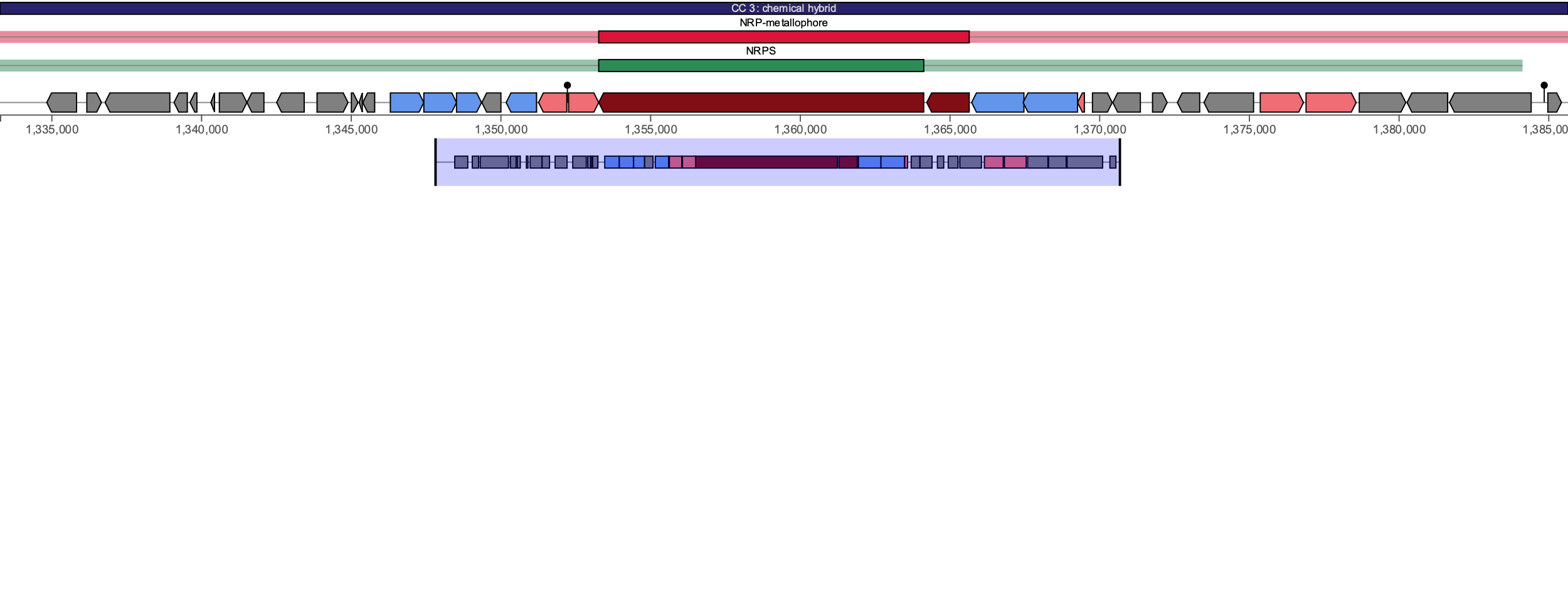

**dltA**

**pvdA**

ABC Transporter

mbtH

alsT

fmt

ABC Transporter

yfmC

fes

fadD

T1PKS

***C. gottingense YSMAA1_1_F6***

(corynecin I/II/III, 13%)

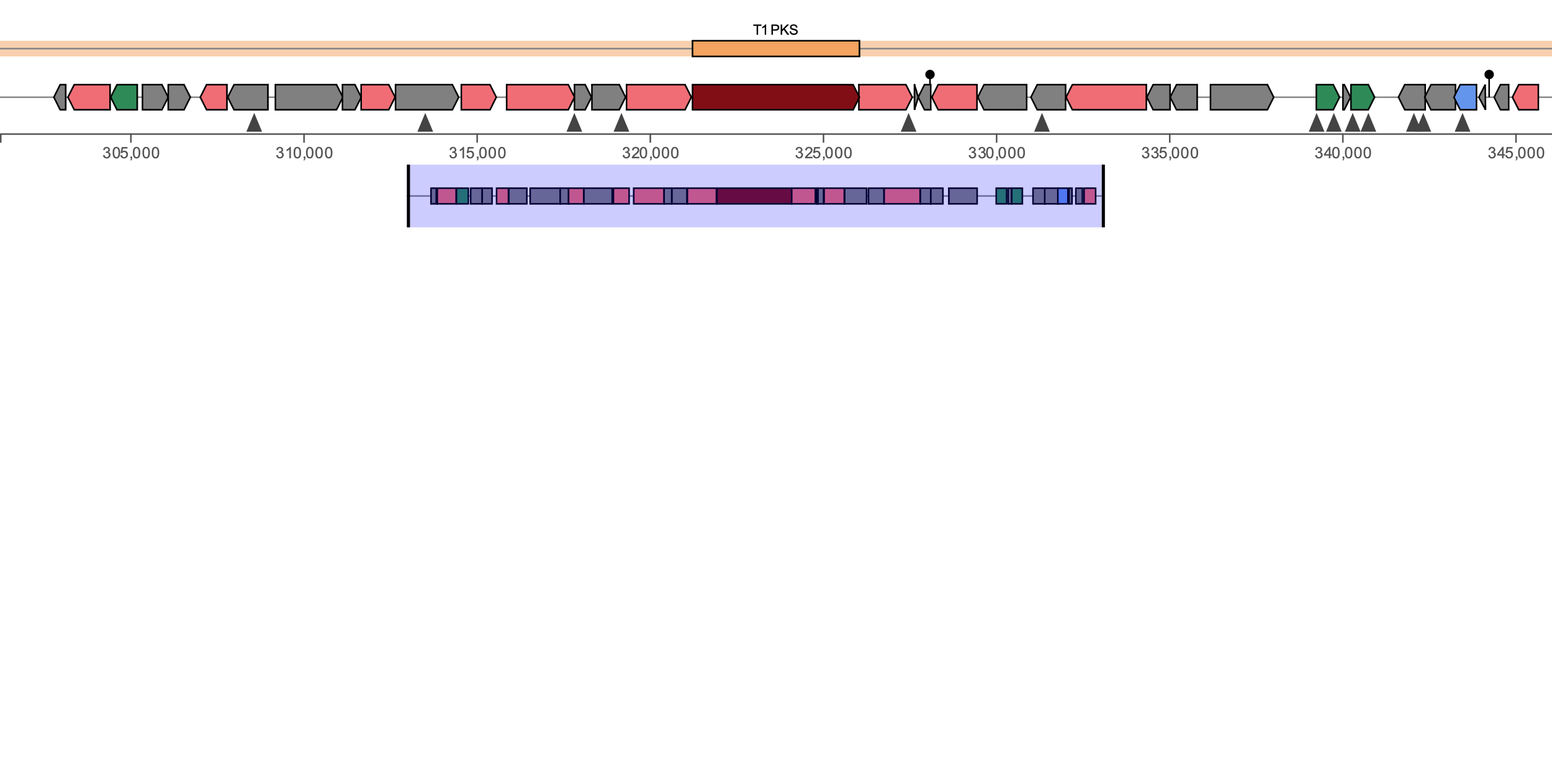

pcaR

catD

ubiA

fbpC

accD5

thlA

**ppsA**

mmpL3

cmtC

cma

NRPS (2)

***C. amycolatum***

**YSMAA1_2_B5**

(phthoxazolin, 4%)

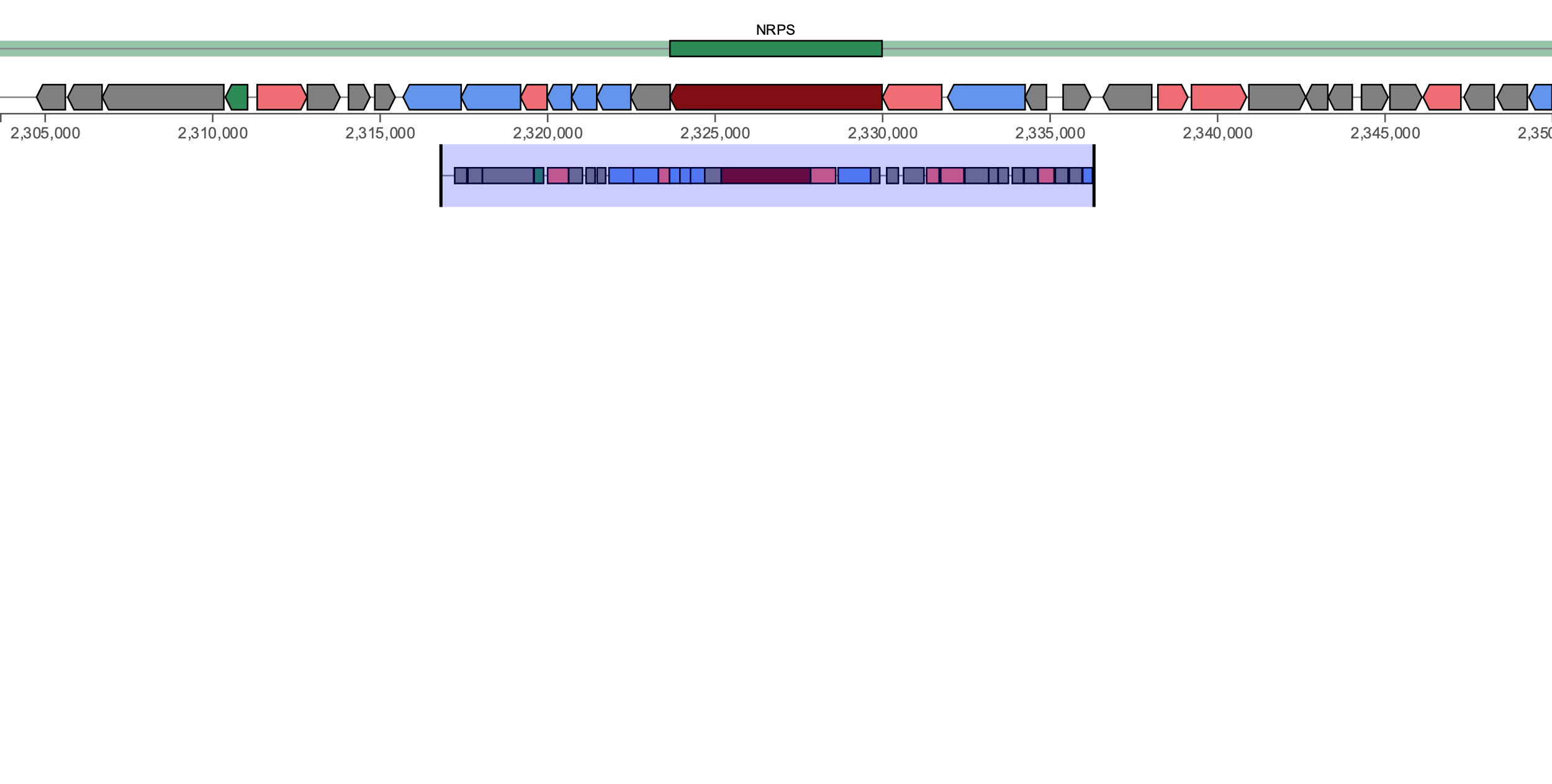

**dltA**

pksJ

btuD

ybiV

prfC

ftsE

caiD

glmU

btuD

irtB

tam

ABC-2 Transporter

**aesC**

**aesB**

**aesA**

mppO

mtdK

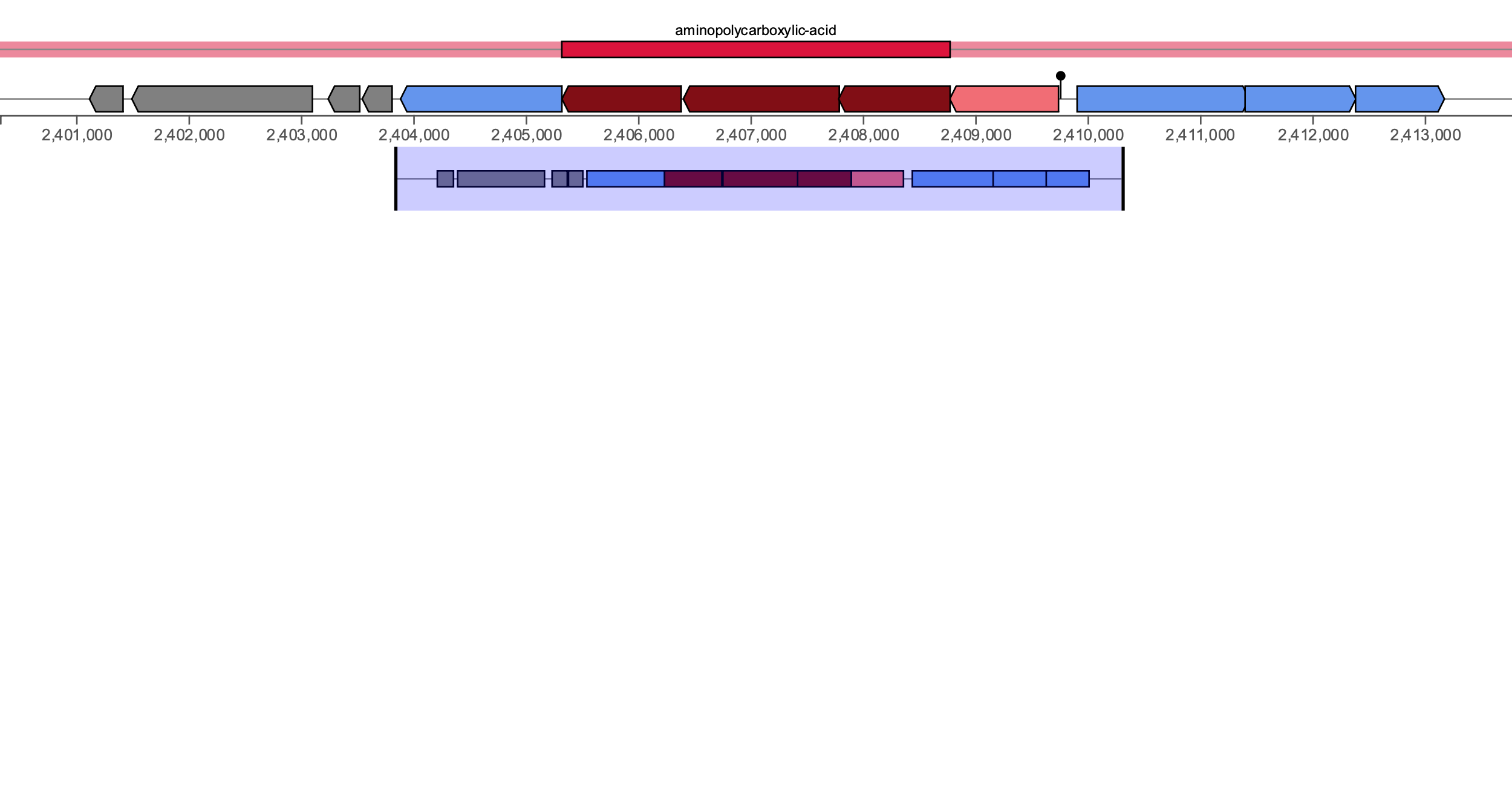

ABC Transporter

Aminopolycarboxylic acid

***C. tuberculostearicum***

**YSMAA1_1_F5**

(EDHA, 33%)

Type VII SS

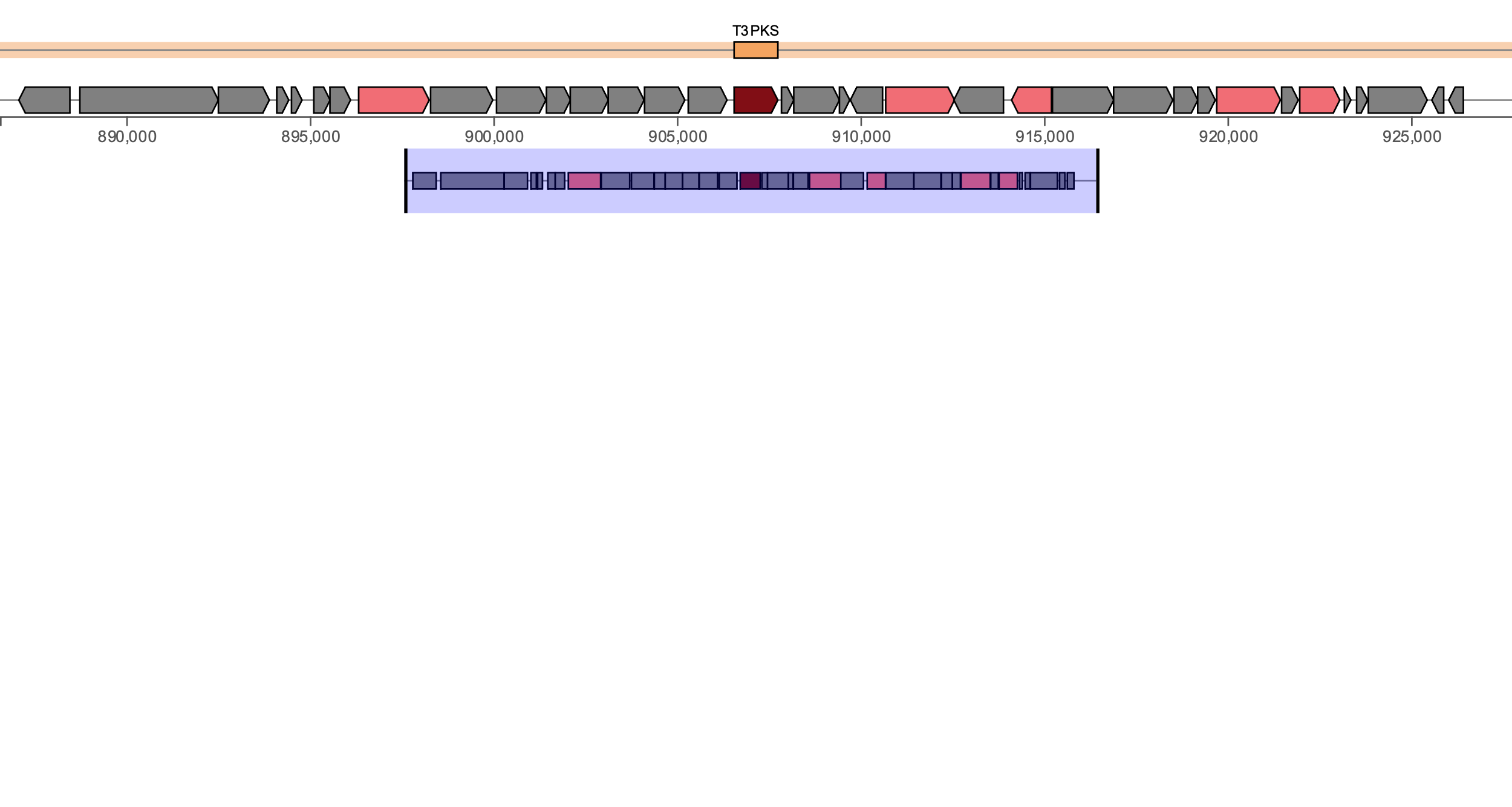

**mvaS**

oatA

glmS

poxB

tsaD

T3PKS

***C. amycolatum***

**YSMAA1_2_B5**

(phenazine SA/SB/SC, 18%)

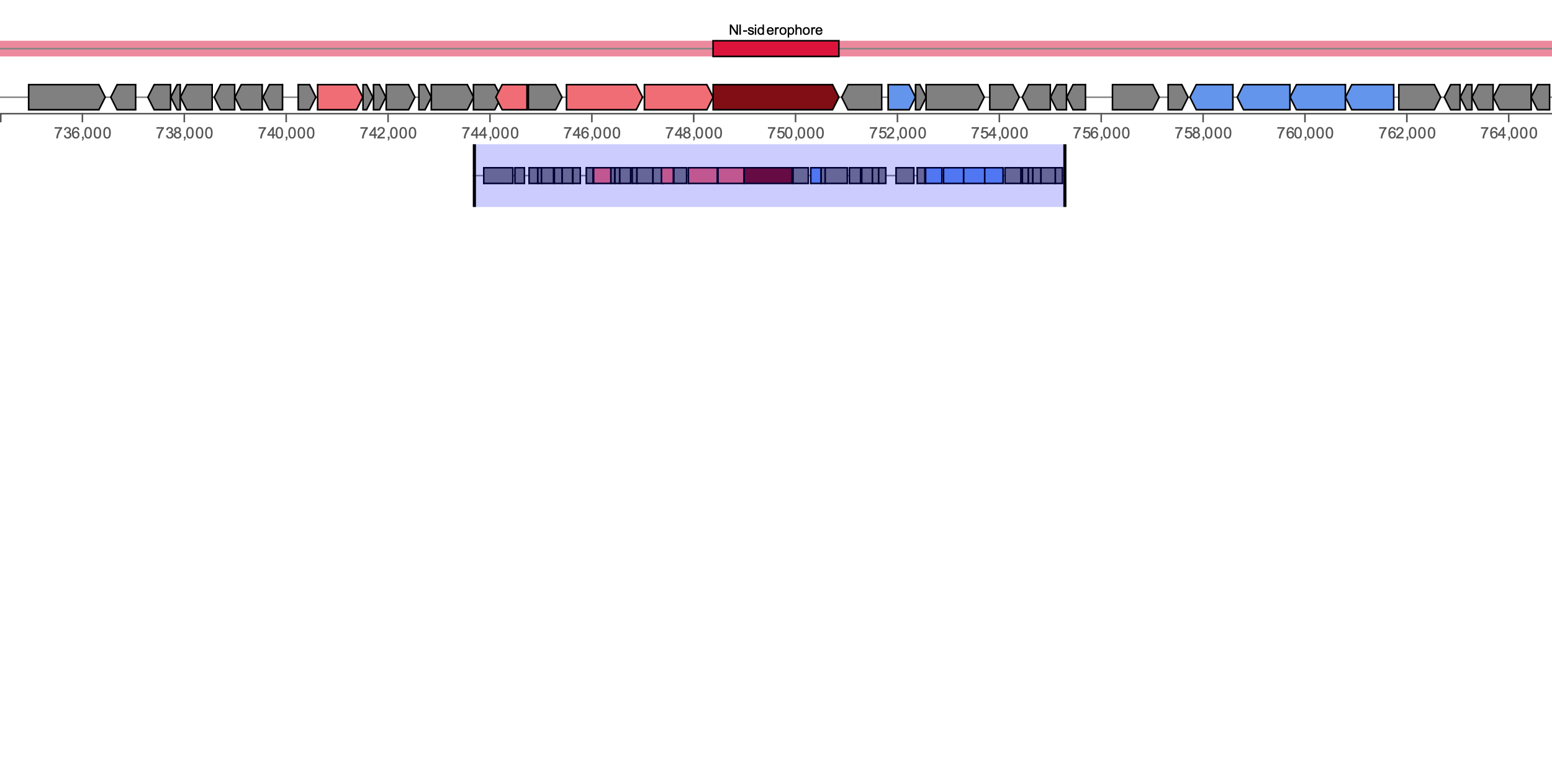

ABC Transporter

lnrL

lcdH

ddc

iucD

NI-siderophore

***C. kefirresidentii***

**YSMAA3_1_H1**

(dehydroxynocardamine, 28%)

**iucC**

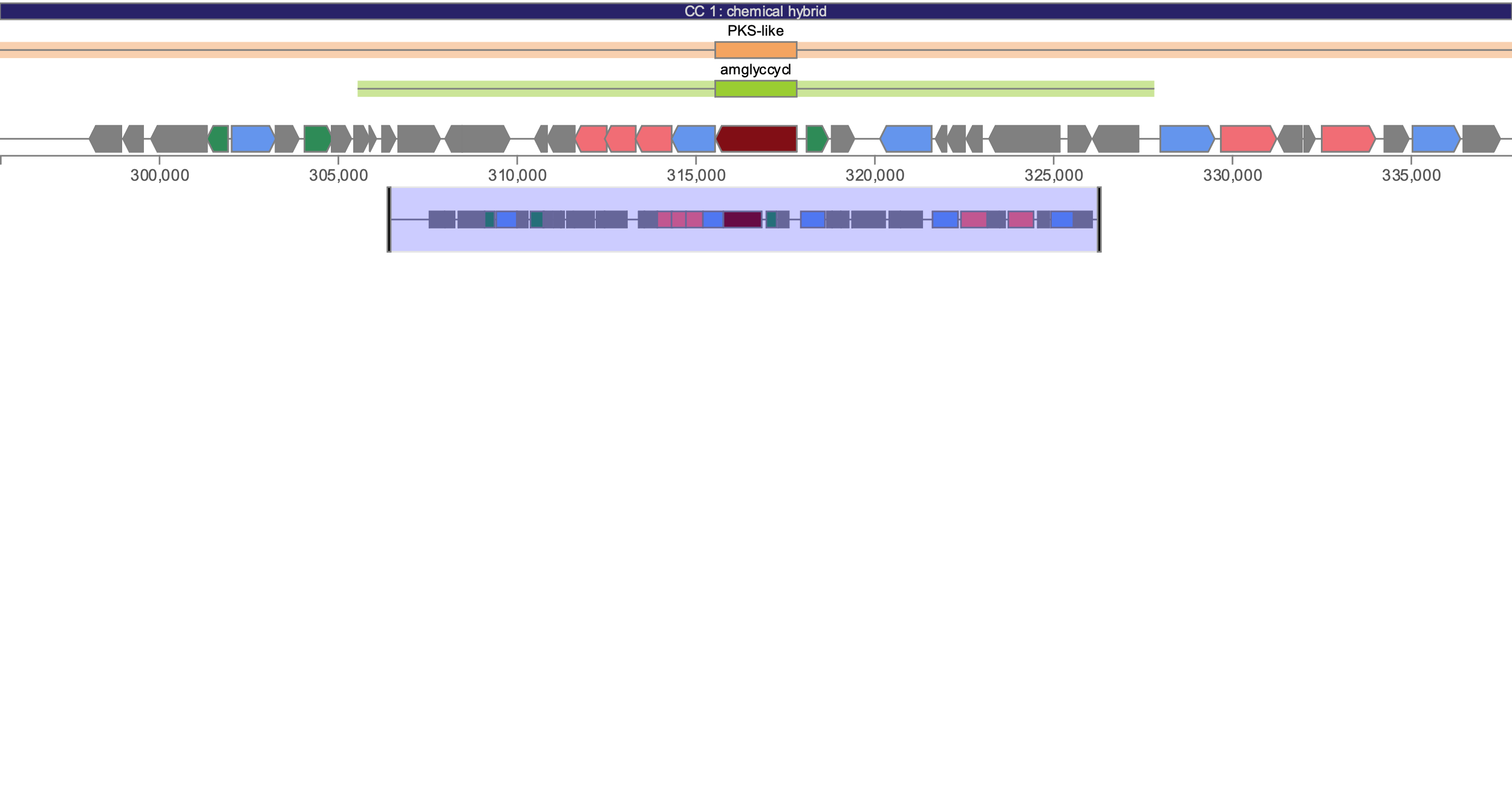

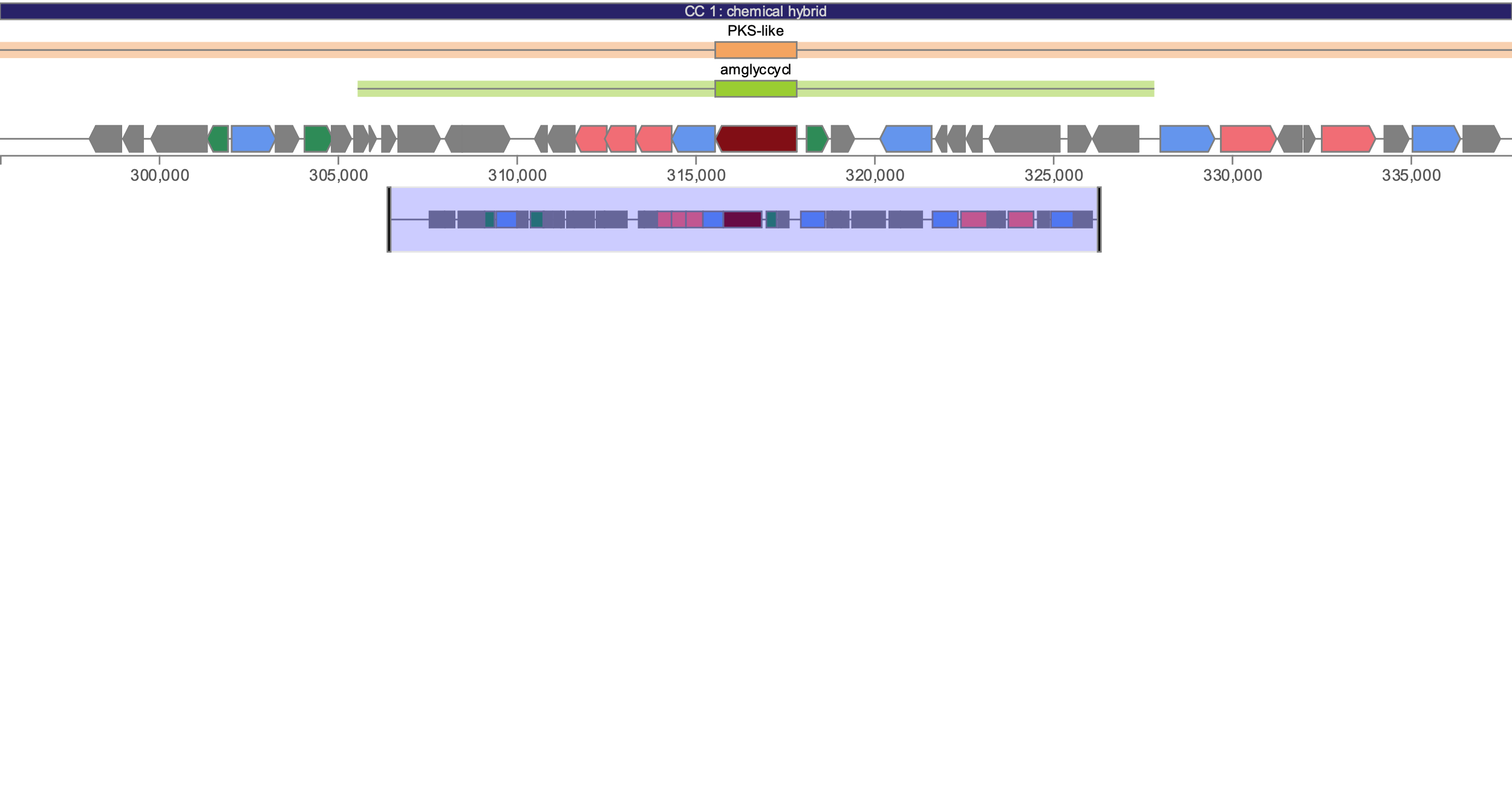

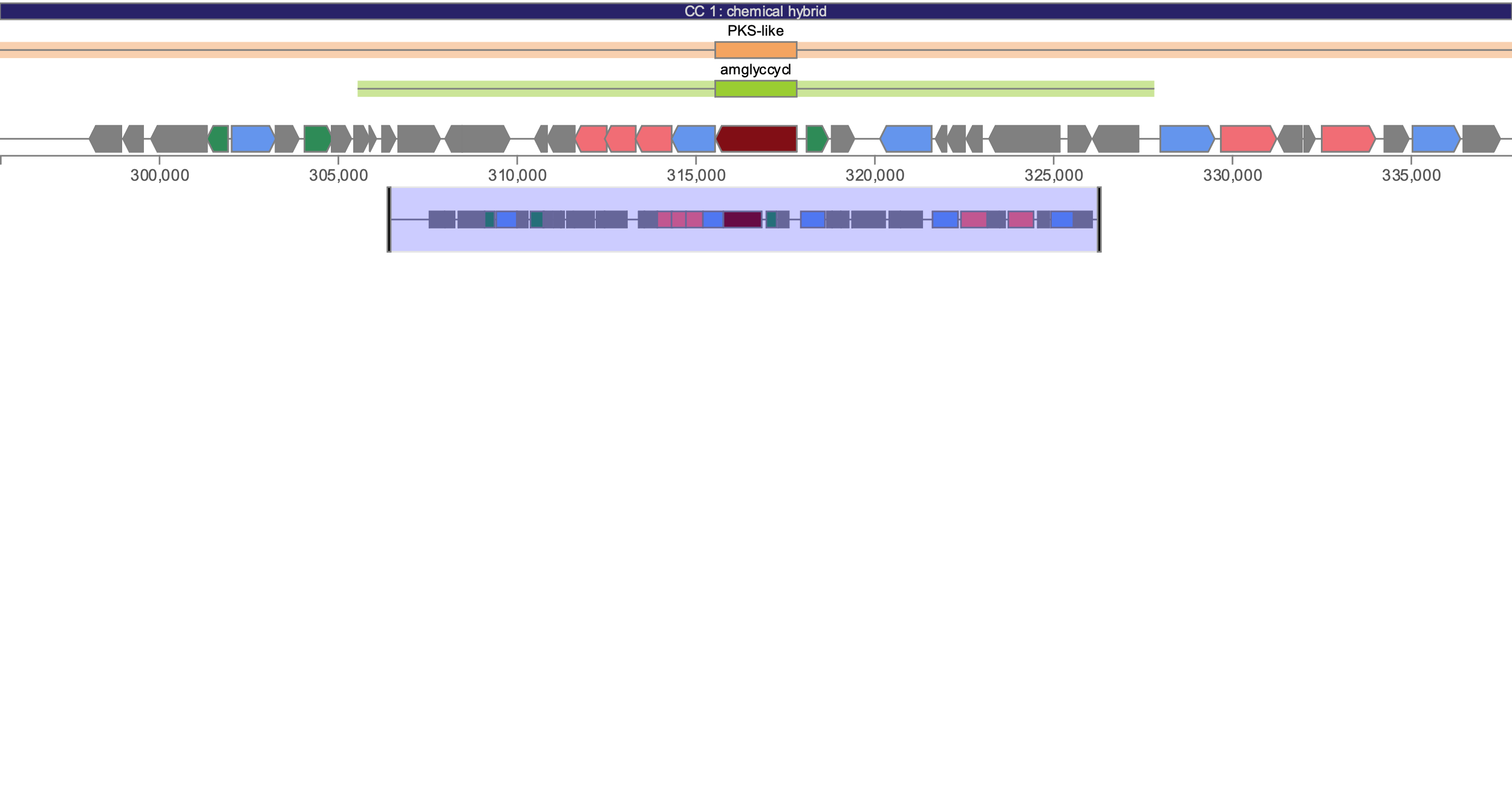

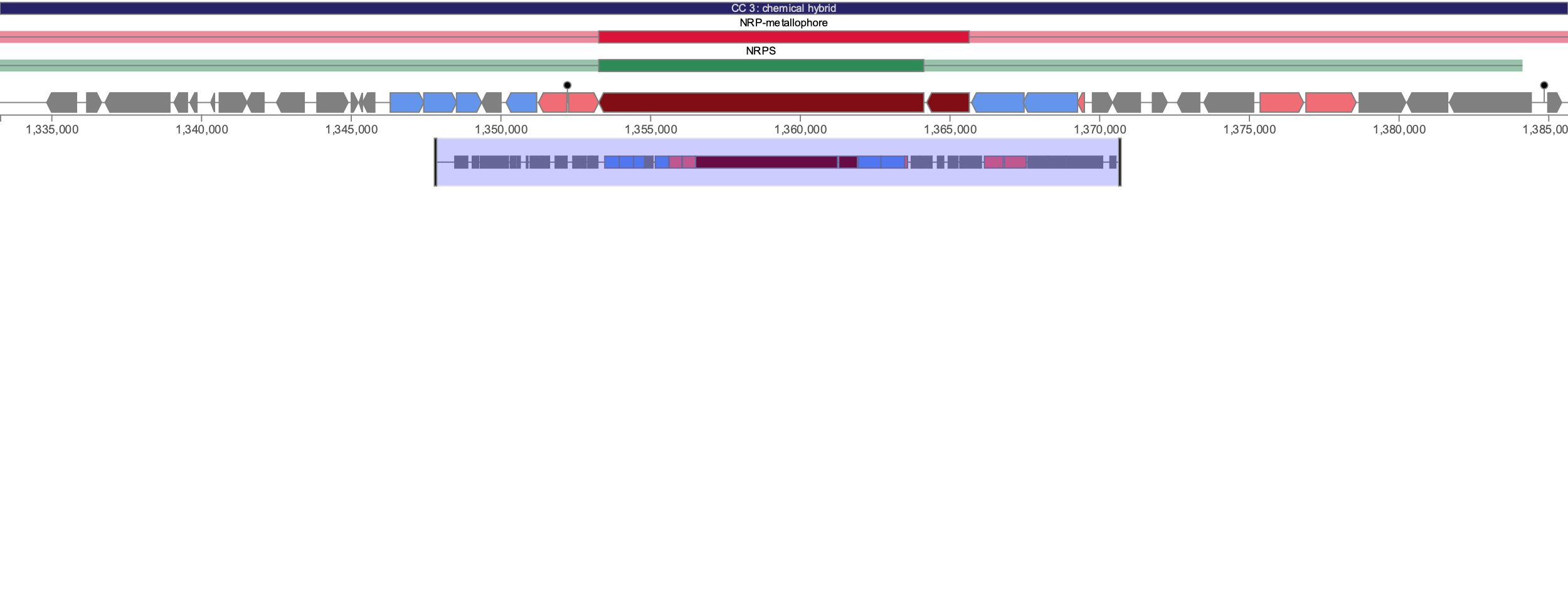

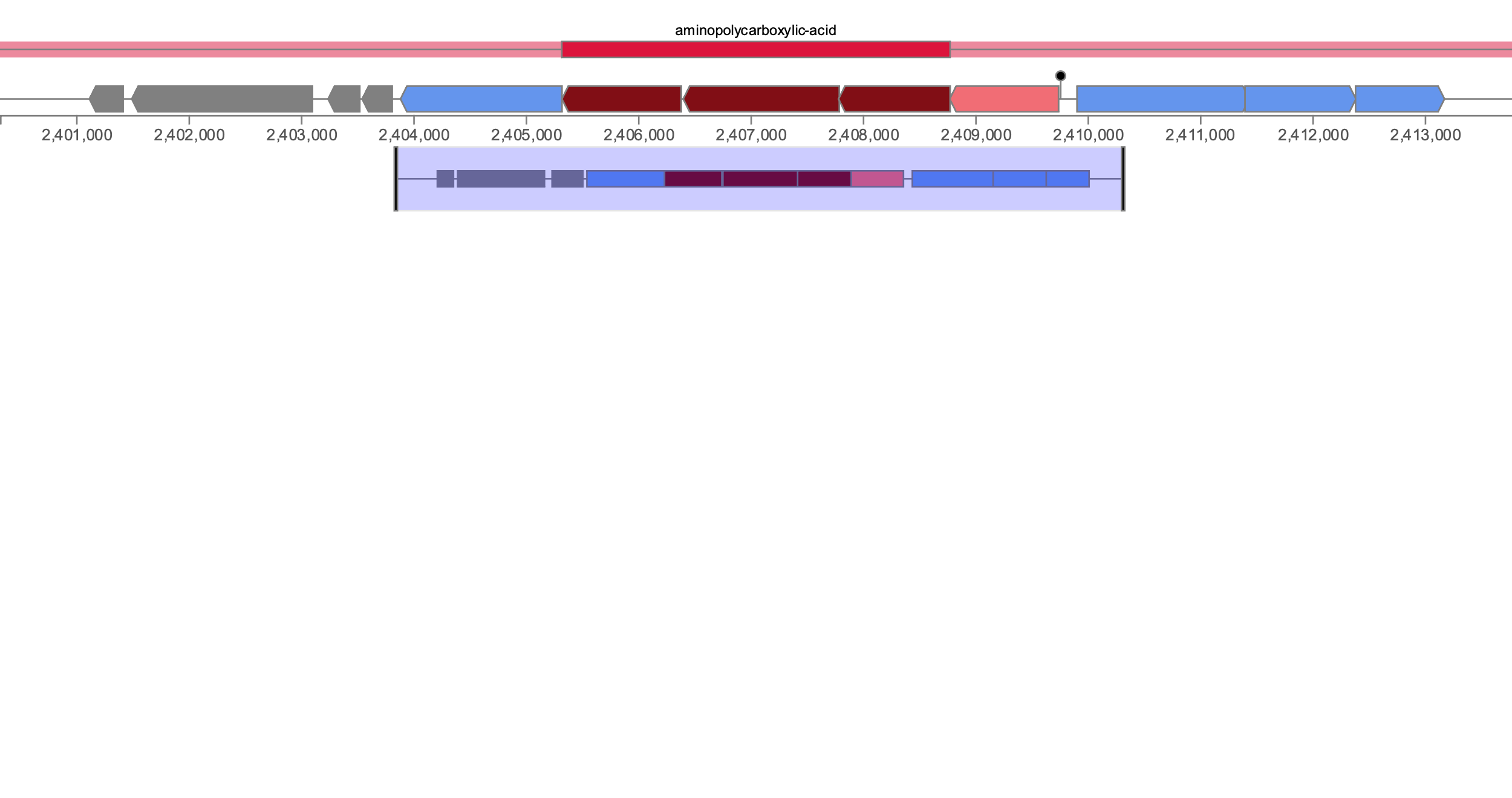

Other genes

Core biosynthetic

Regulatory

Transport

Additional biosynthetic

Legend:

**Supplementary Figure 12**. Architectures of the 8 distinct putative biosynthetic gene clusters predicted using antiSMASH 7.

***C. marquesiae***

A1_1_H1, A1_1_H4, A1_2_D10, A3_1_C3, A3_1_F3, A3_1_G3, A4_1_B7, A4_1_E7, A4_1_E8, A5_1_H11

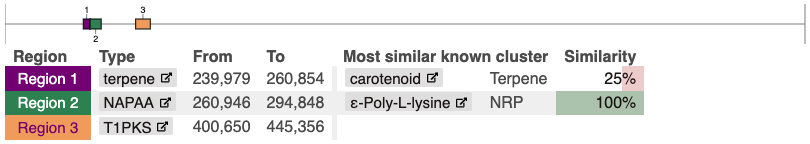

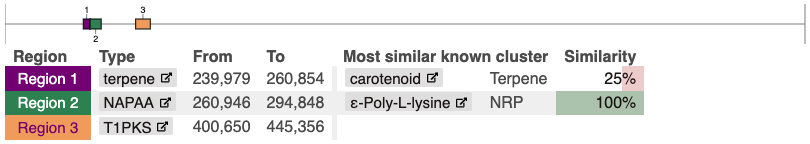

*C. marquesiae A1_1_H4 also includes an aminopolycarboxylic acid cluster (most similar known cluster: EDHA, 33%)*

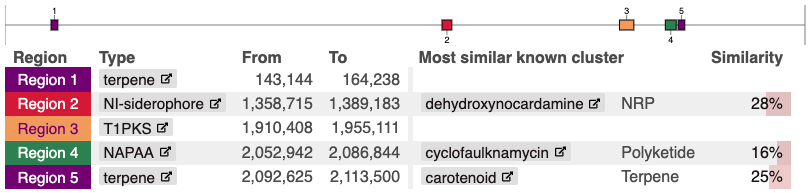

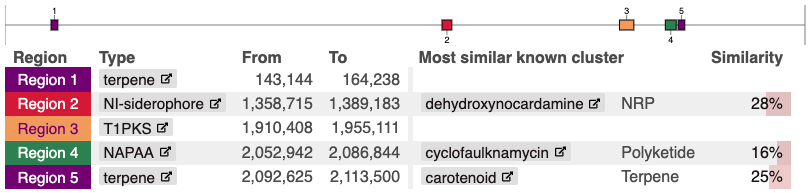

***C. kefirresidentii***

A1_1_C6, A1_2_B7, A3_1_H1, A5_1_D11

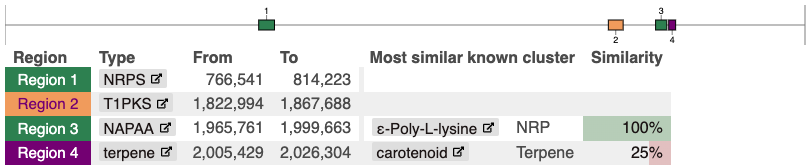

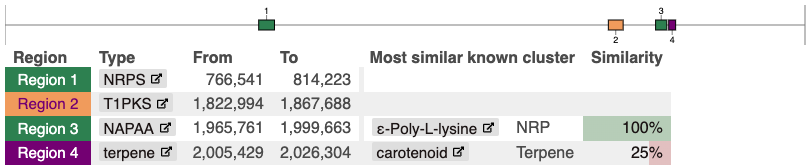

***C. kefirresidentii***

A1_2_G9

***C. tuberculostearicum***

A3_1_A4

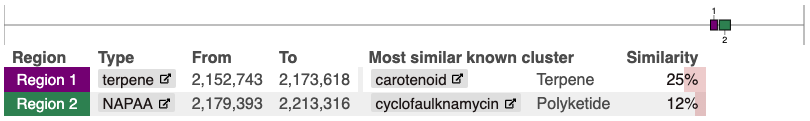

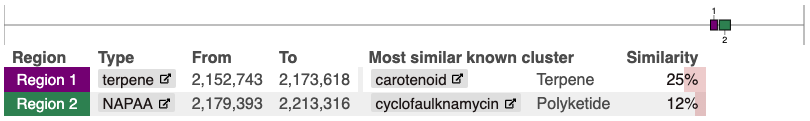

***C. tuberculostearicum***

A5_1_B12, A5_1_F11

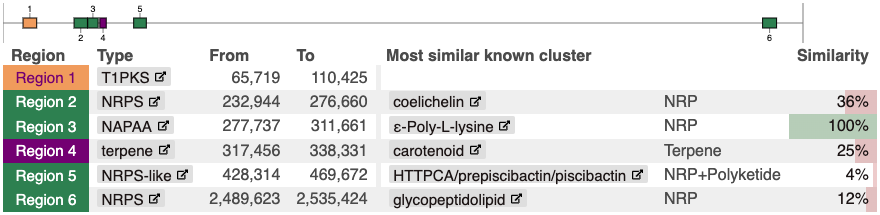

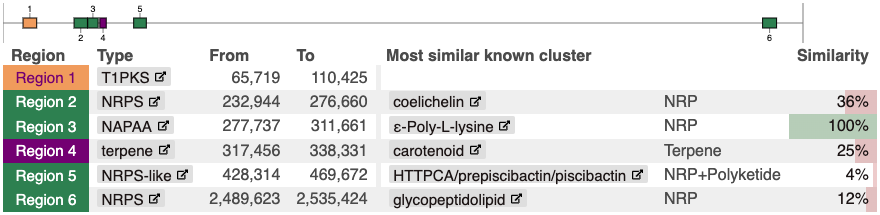

***C. tuberculostearicum***

A1_2_H3, A5_1_G11

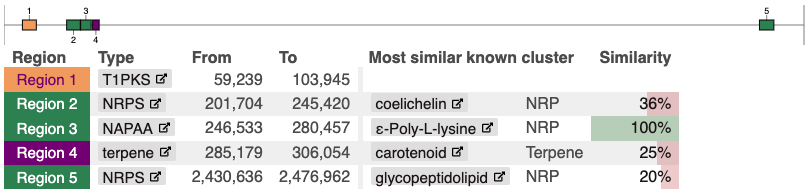

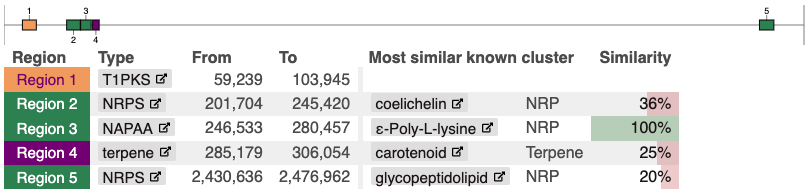

*C. tuberculostearicum A1_1_F5 also includes an aminopolycarboxylic acid cluster (most similar known cluster: EDHA, 33%)*

***C. tuberculostearicum***

A1_1_B1, A1_1_C1, A1_1_D8, A1_1_F5

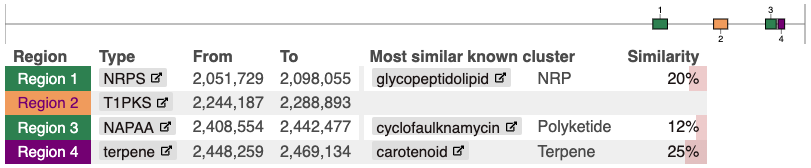

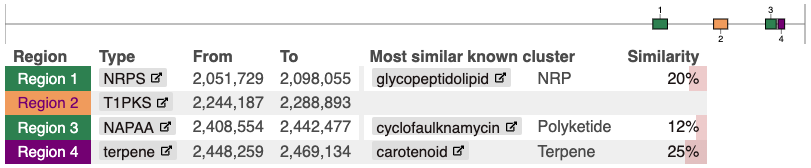

**Supplementary Figure 13.** The distribution of putative biosynthetic gene clusters in the *C. tuberculostearicum* species complex predicted using antiSMASH 7.

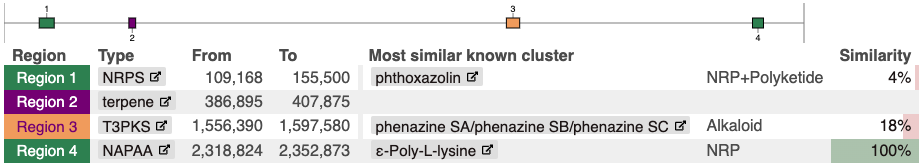

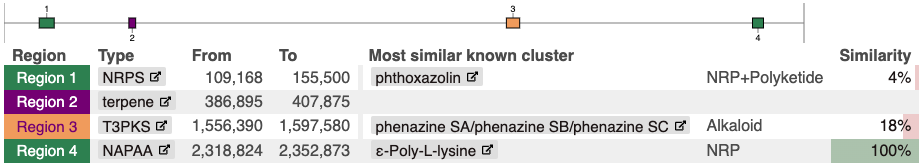

***C. amycolatum***

A1_2_B5, A1_2_C5

***C. gottingense***

A1_1_F6

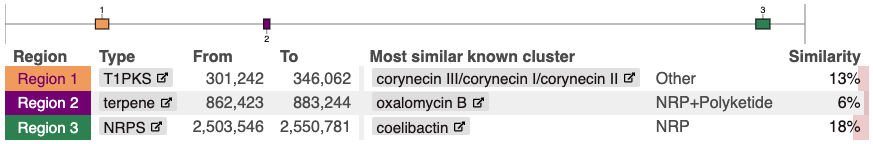

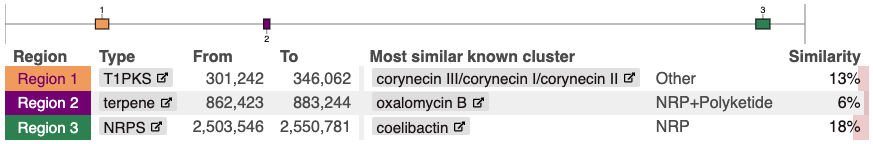

***C. axilliensis***

A1_1_D6, A5_1_F9

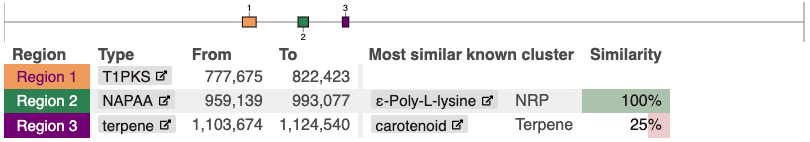

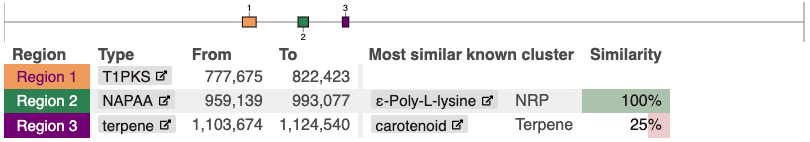

***C. jamesii***

A1_1_F7

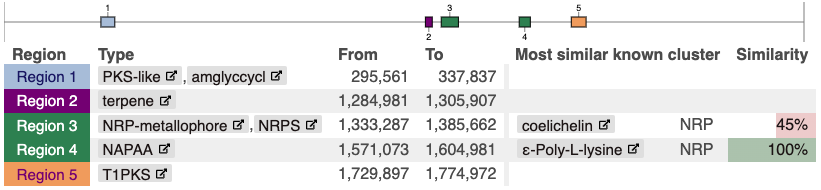

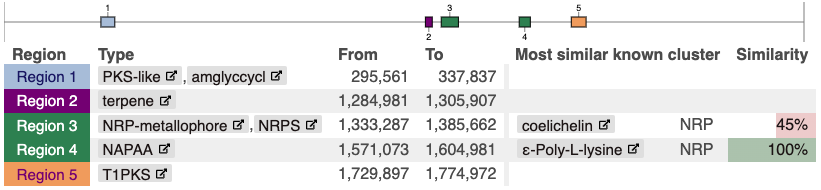

**Supplementary Figure 14.** The distribution of putative biosynthetic gene clusters in the *C. amycolatum, C. gottingense, C. axilliensis and C jamesii* isolates predicted using antiSMASH 7.

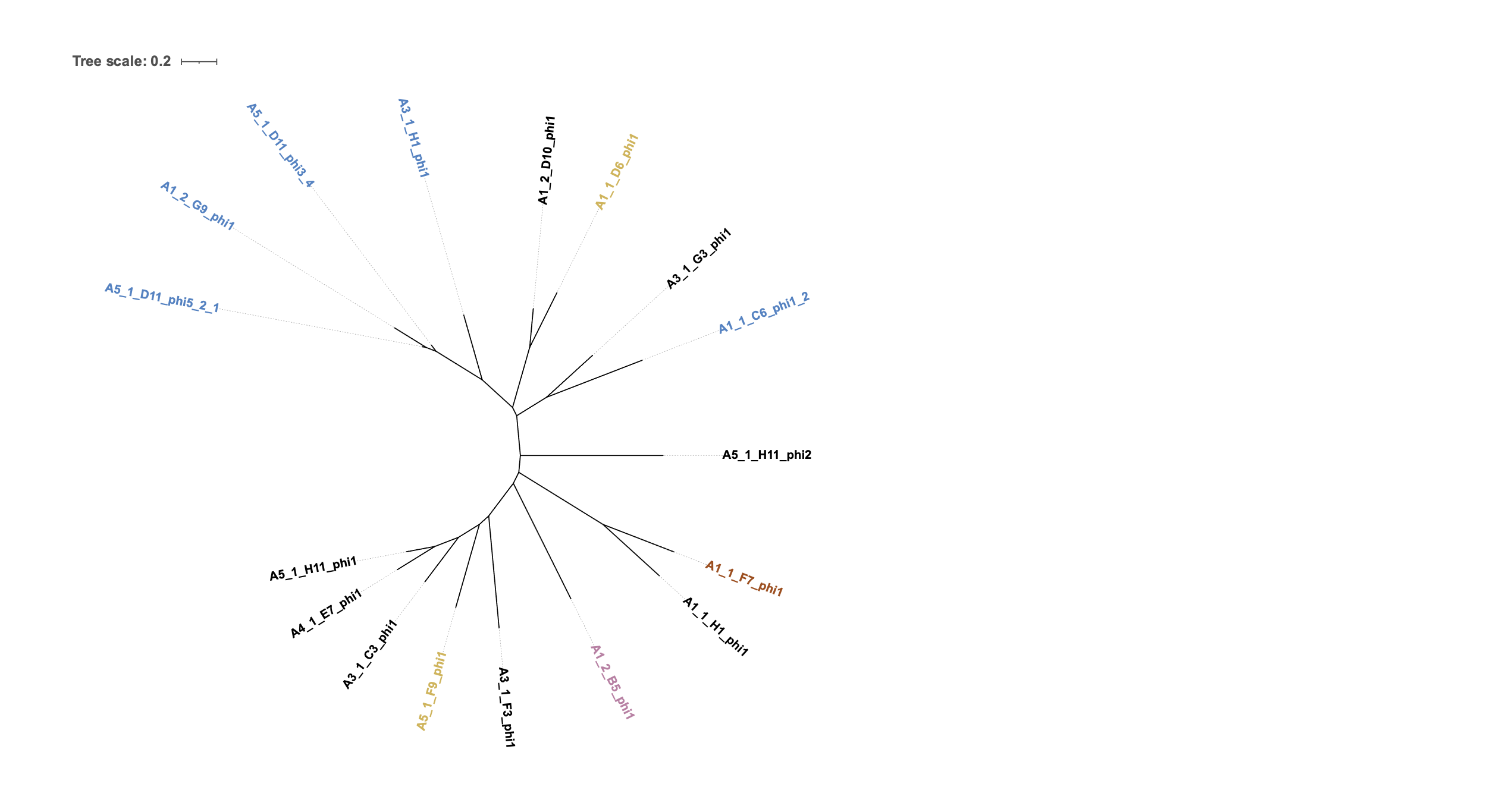

**Supplementary Figure 15.** None of the putative prophages were completely identical to each other. Putative prophages were aligned and compared using an unrooted tree generated with PhyML and visualised on iTOL. Putative prophages were coloured by the species of the source isolate (Blue: *C. kefirresidentii*, Black: *C. marquesiae*, Gold: *C. axilliensis*, Orange: *C. jamesii* and Pink: *C. amycolatum*).

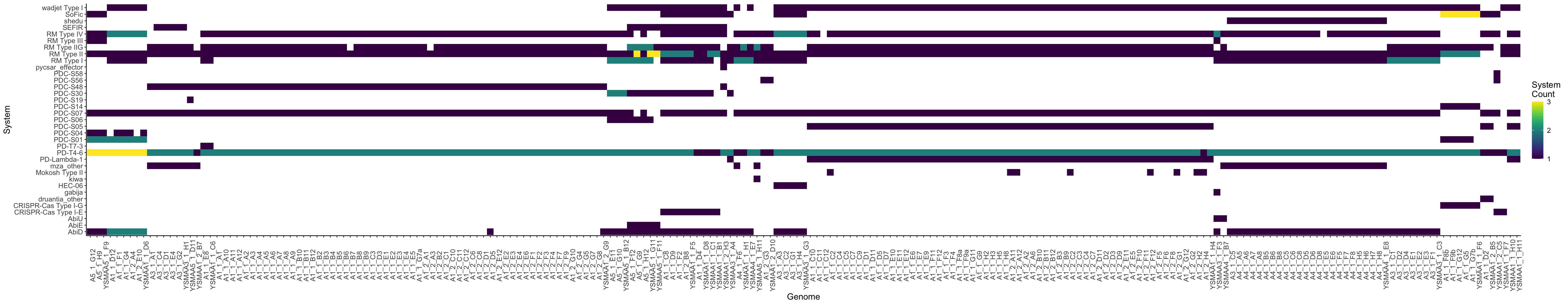

**Supplementary Figure 16.** Global PADLOC analysis reveals profiles of phage defence systems are generally conserved within drep secondary clusters.

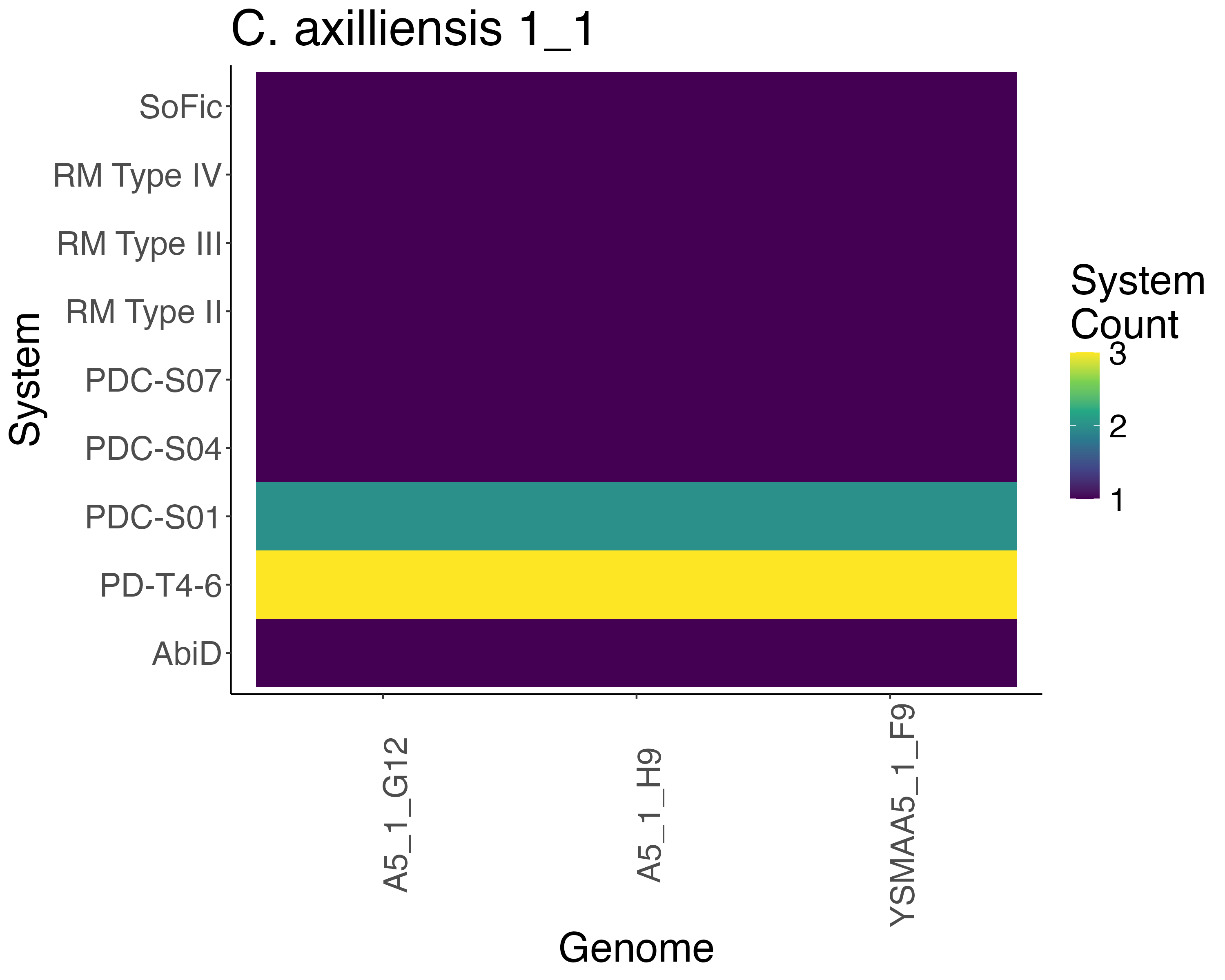

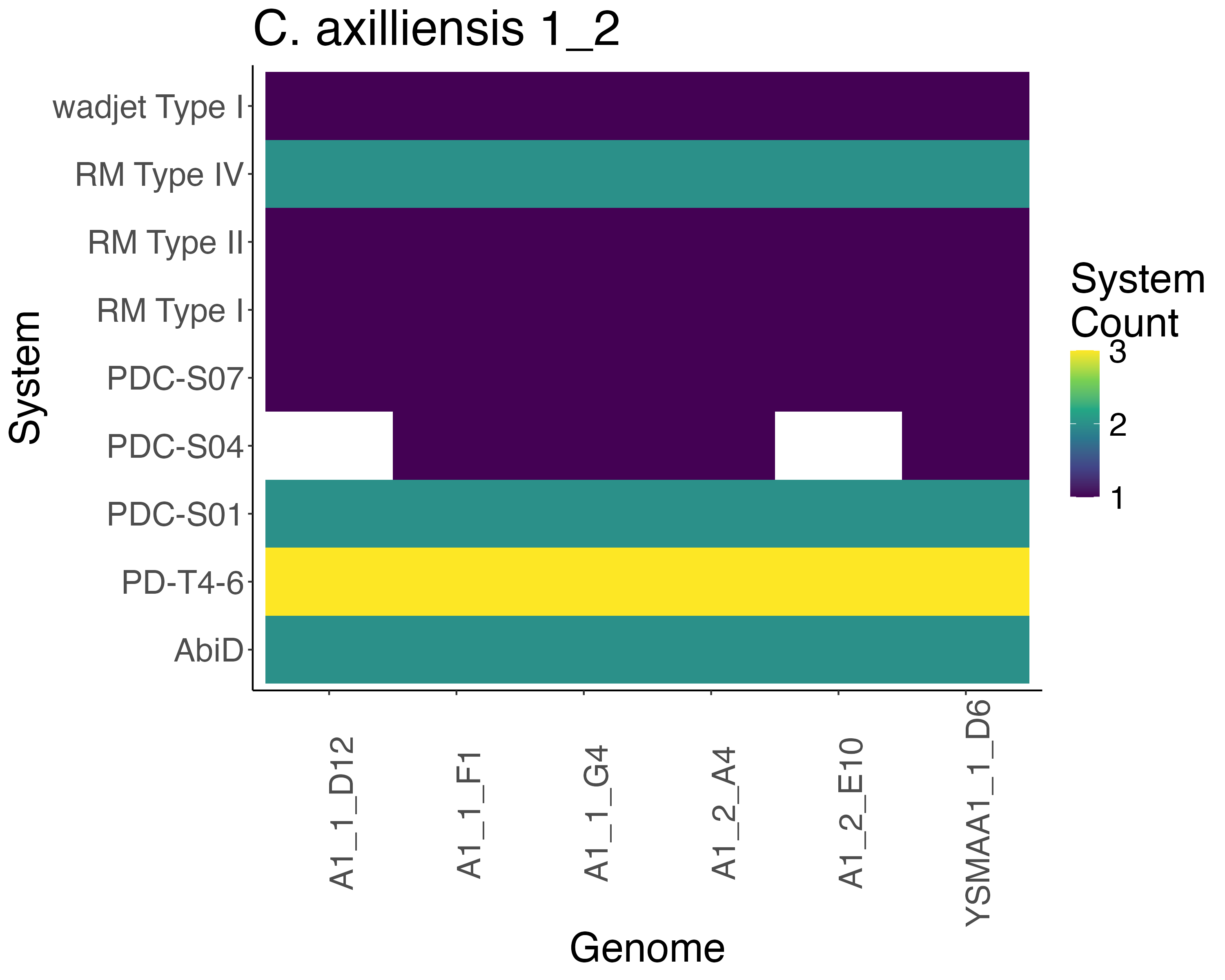

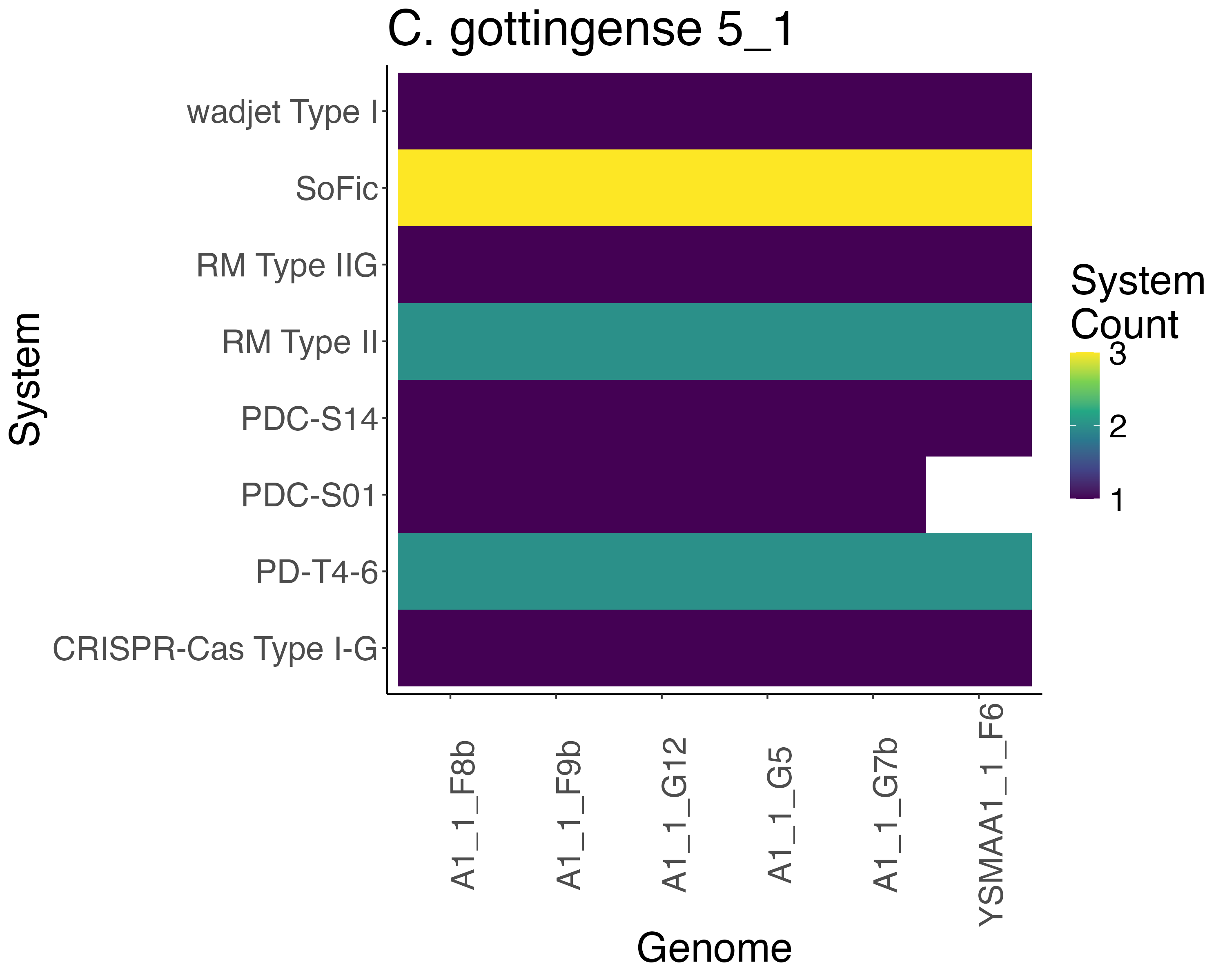

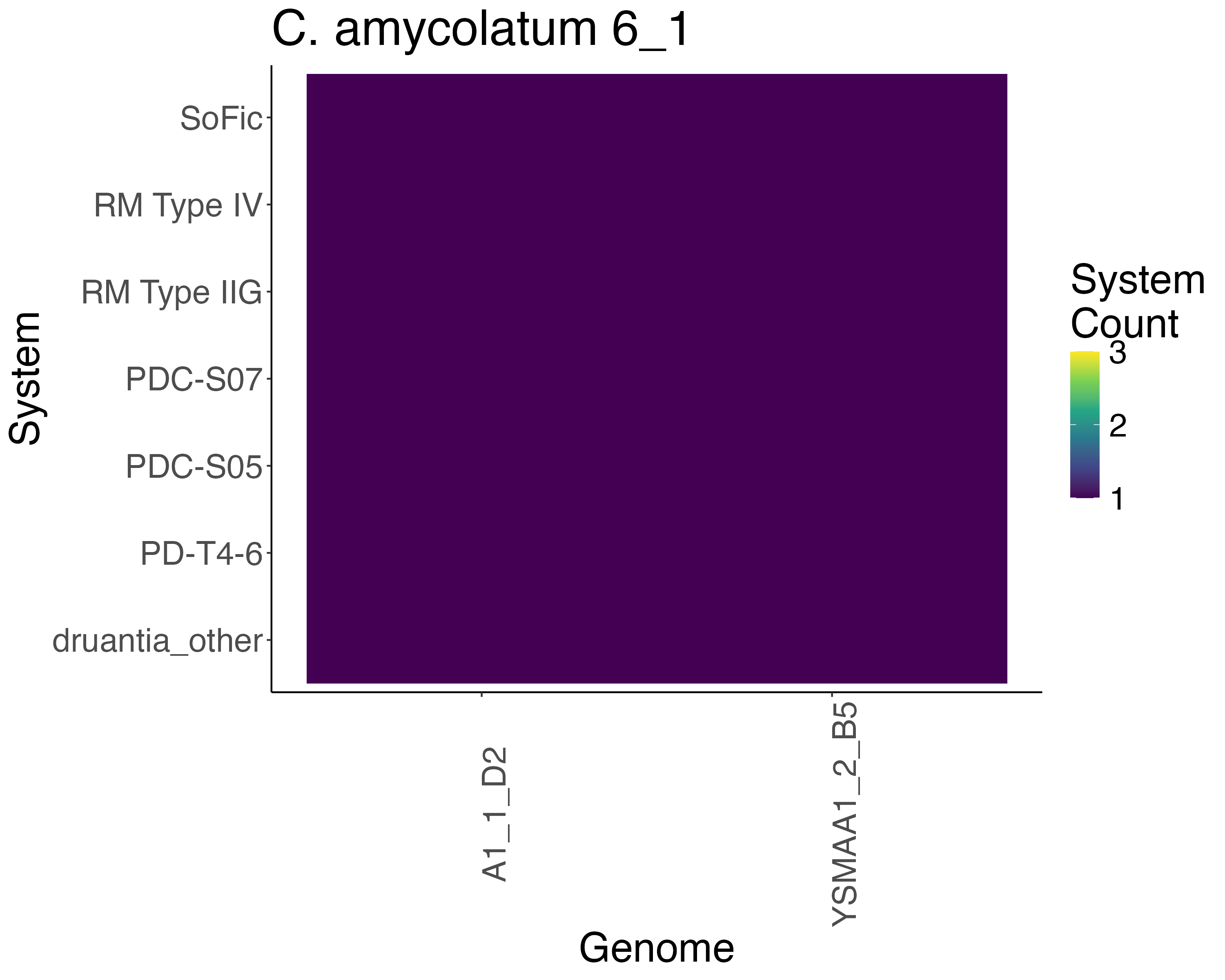

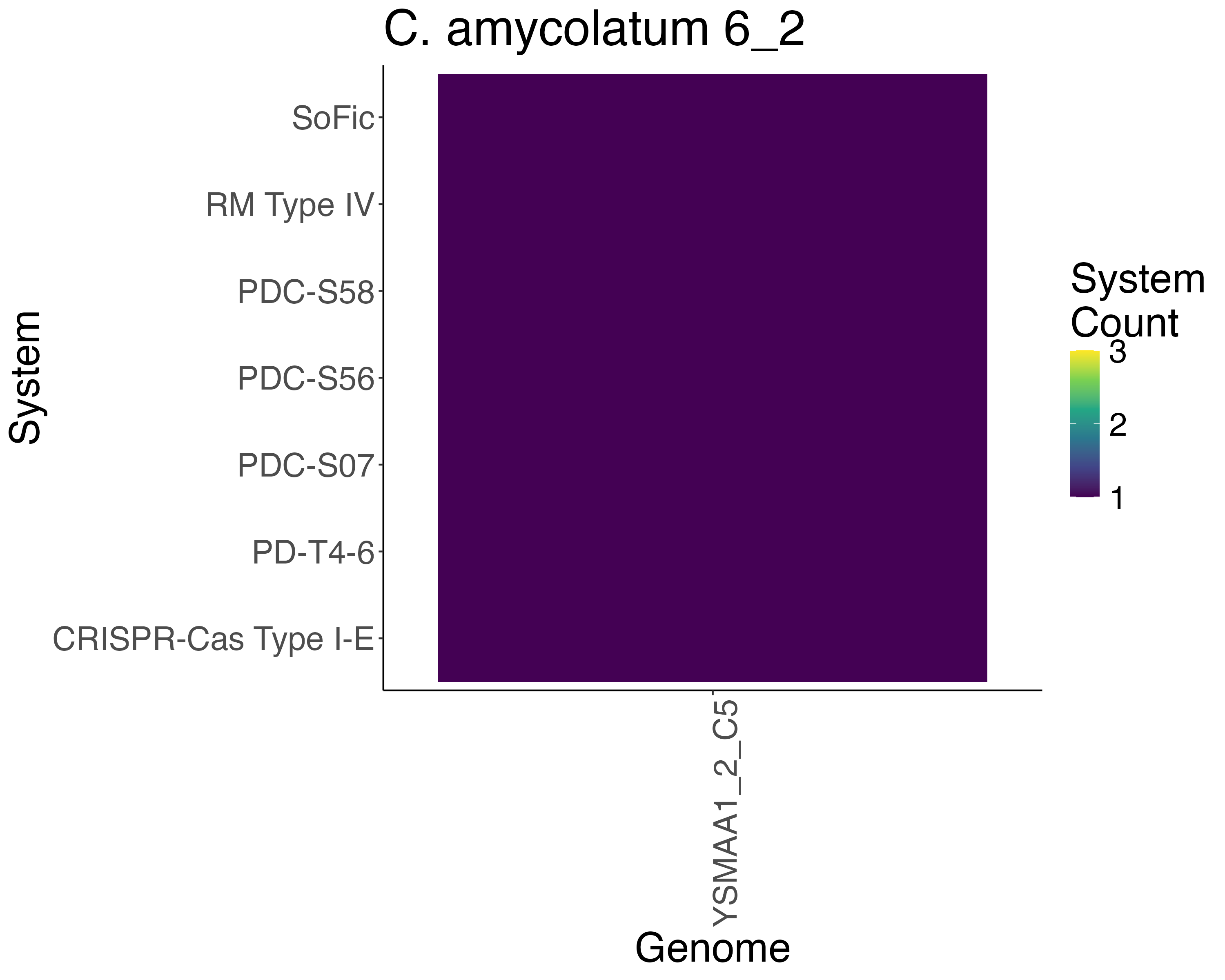

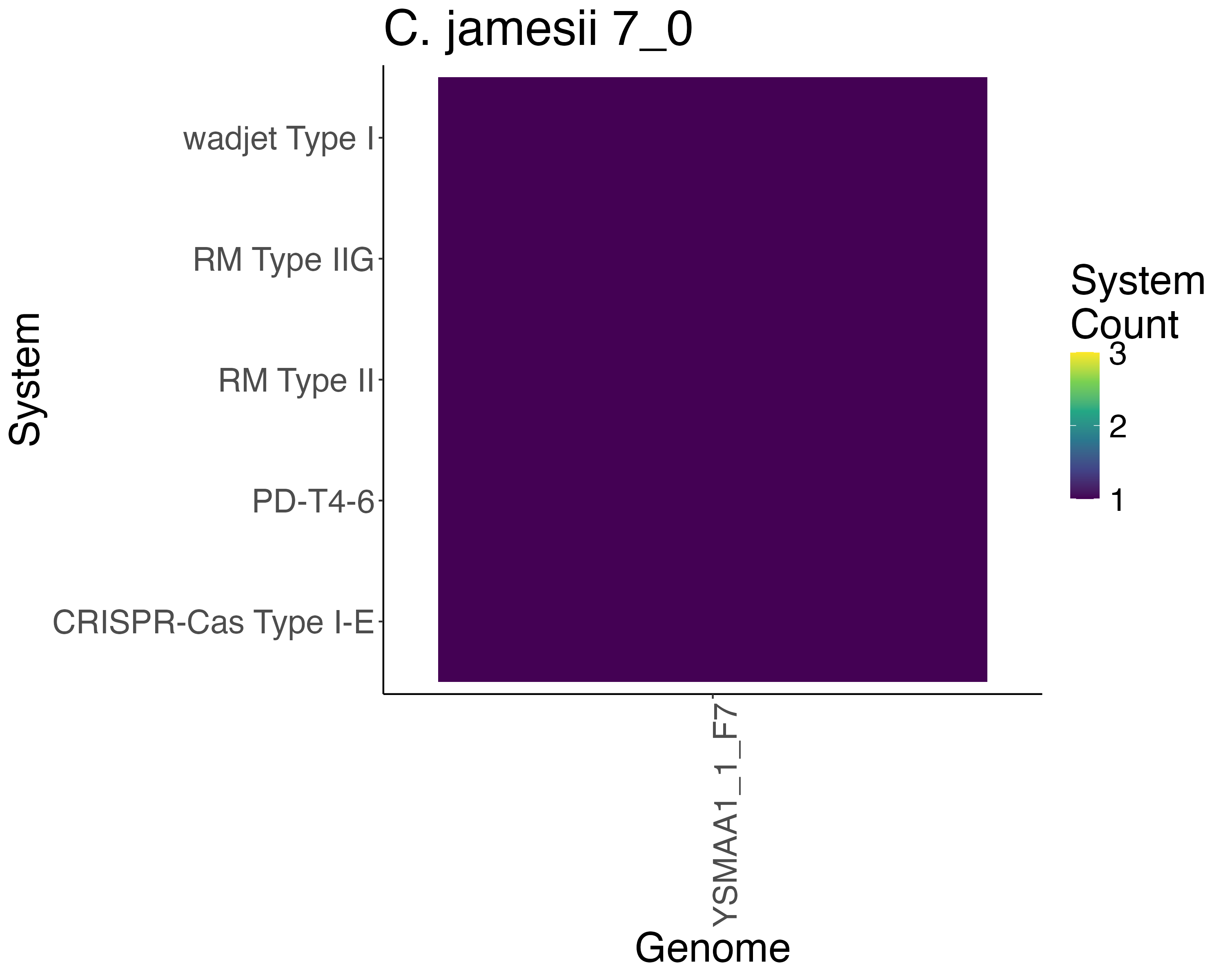

**Supplementary Figure 17. Phage defence systems found in drep secondary clusters of (A-B) *C. axilliensis*, (C) *C. jamesii*, (D-E) *C. amycolatum* and (F) *C. gottingense*.** The heatmap describes the number of each identified system with (Yellow: 3, Green: 2, Purple: 1 and White: 0).

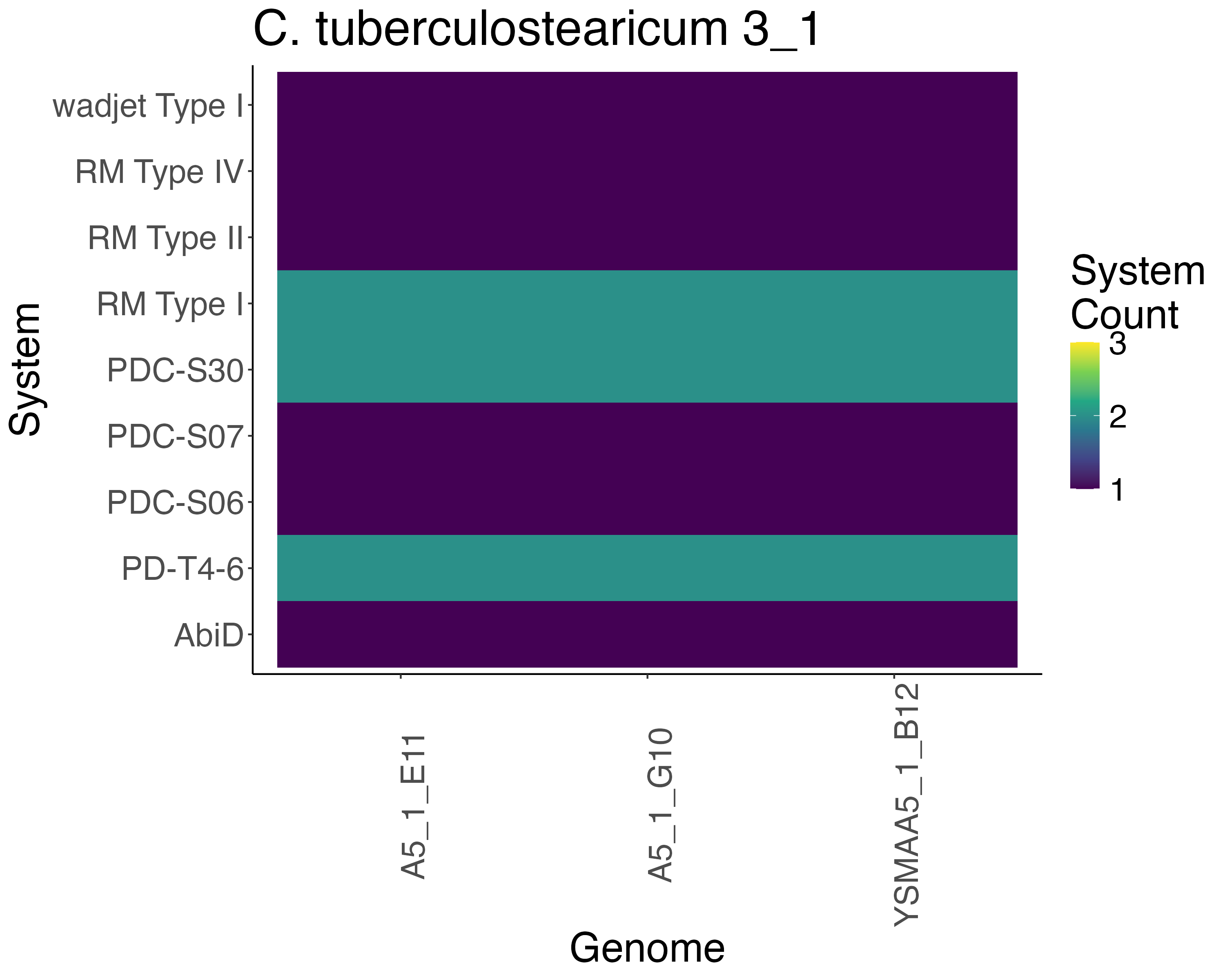

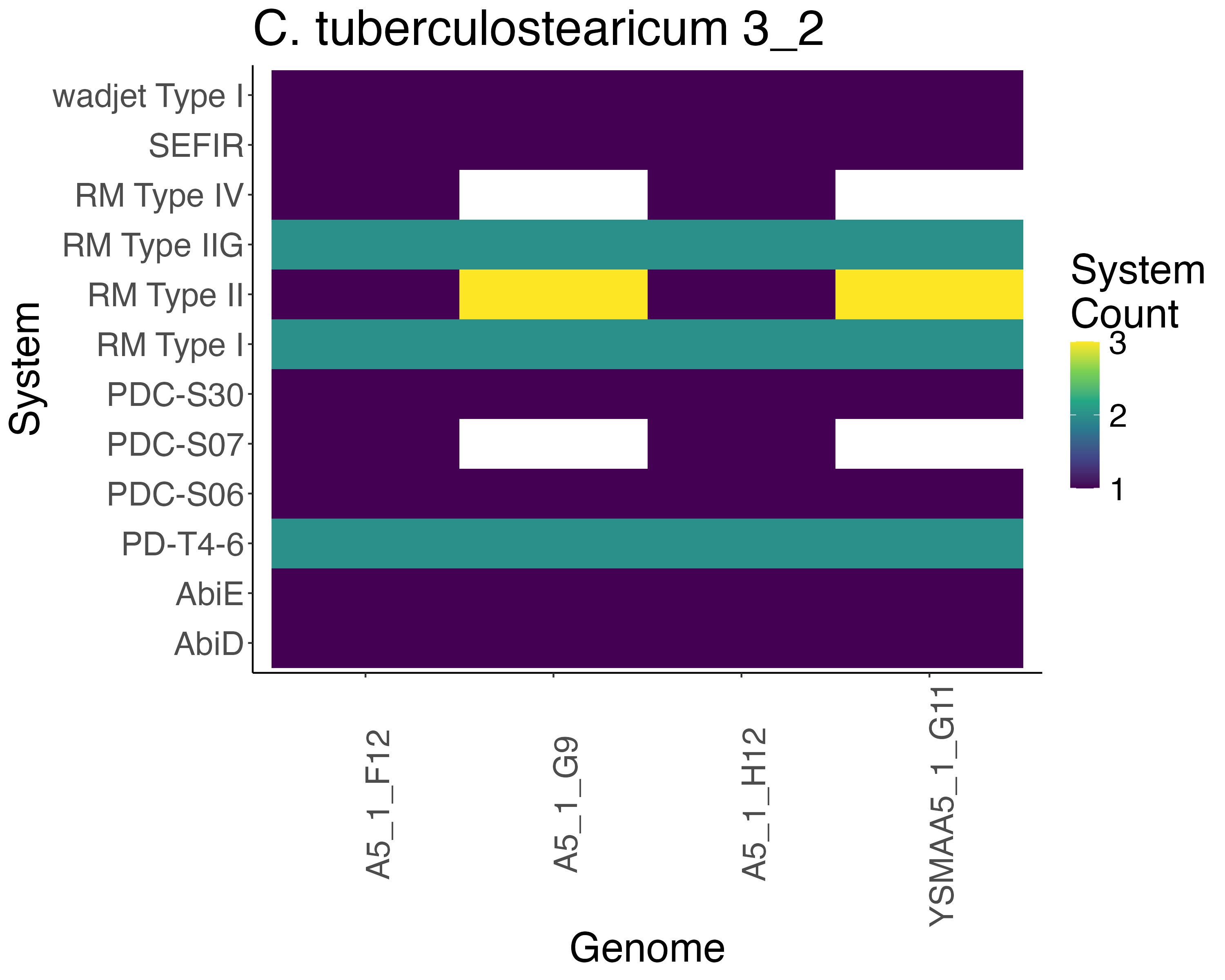

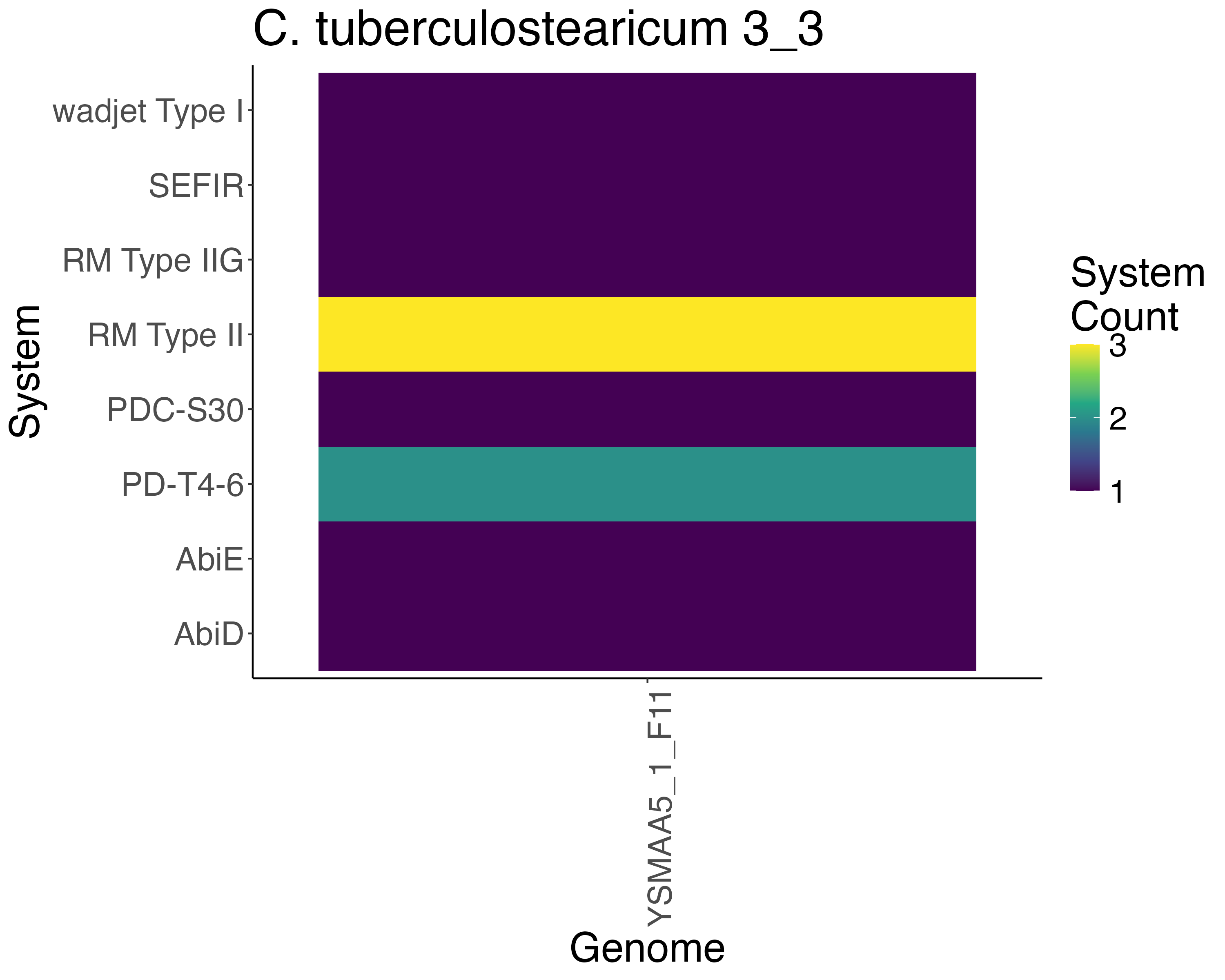

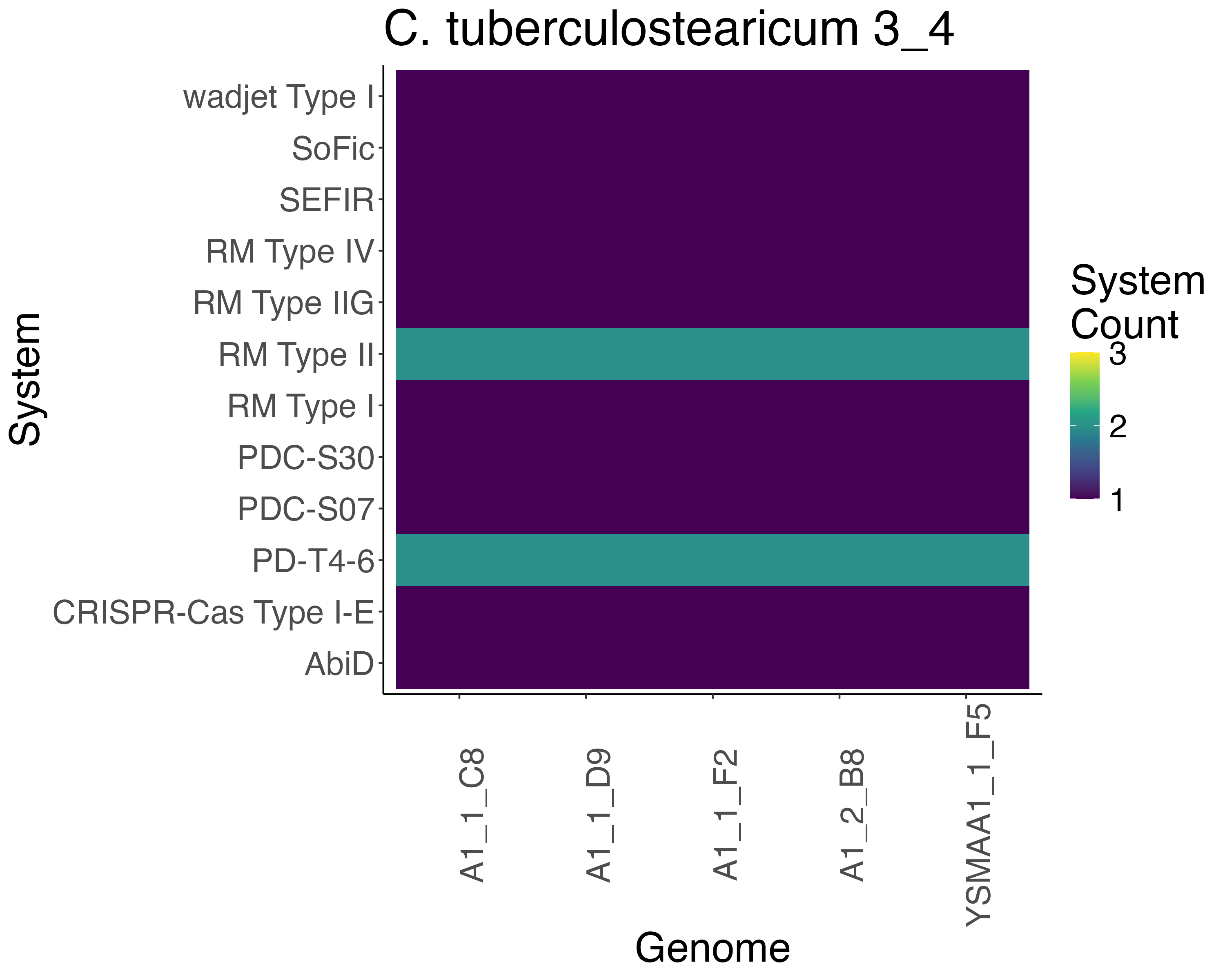

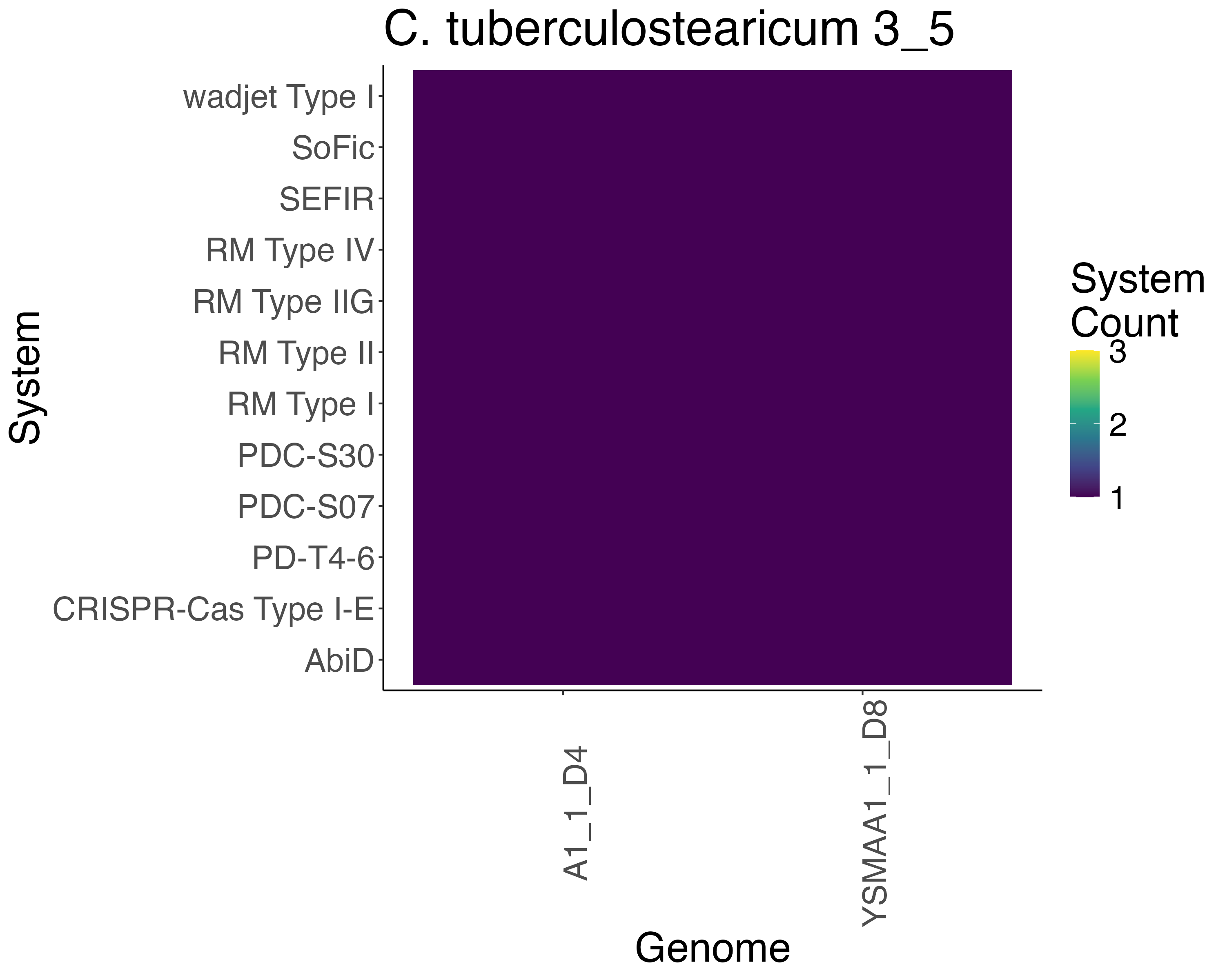

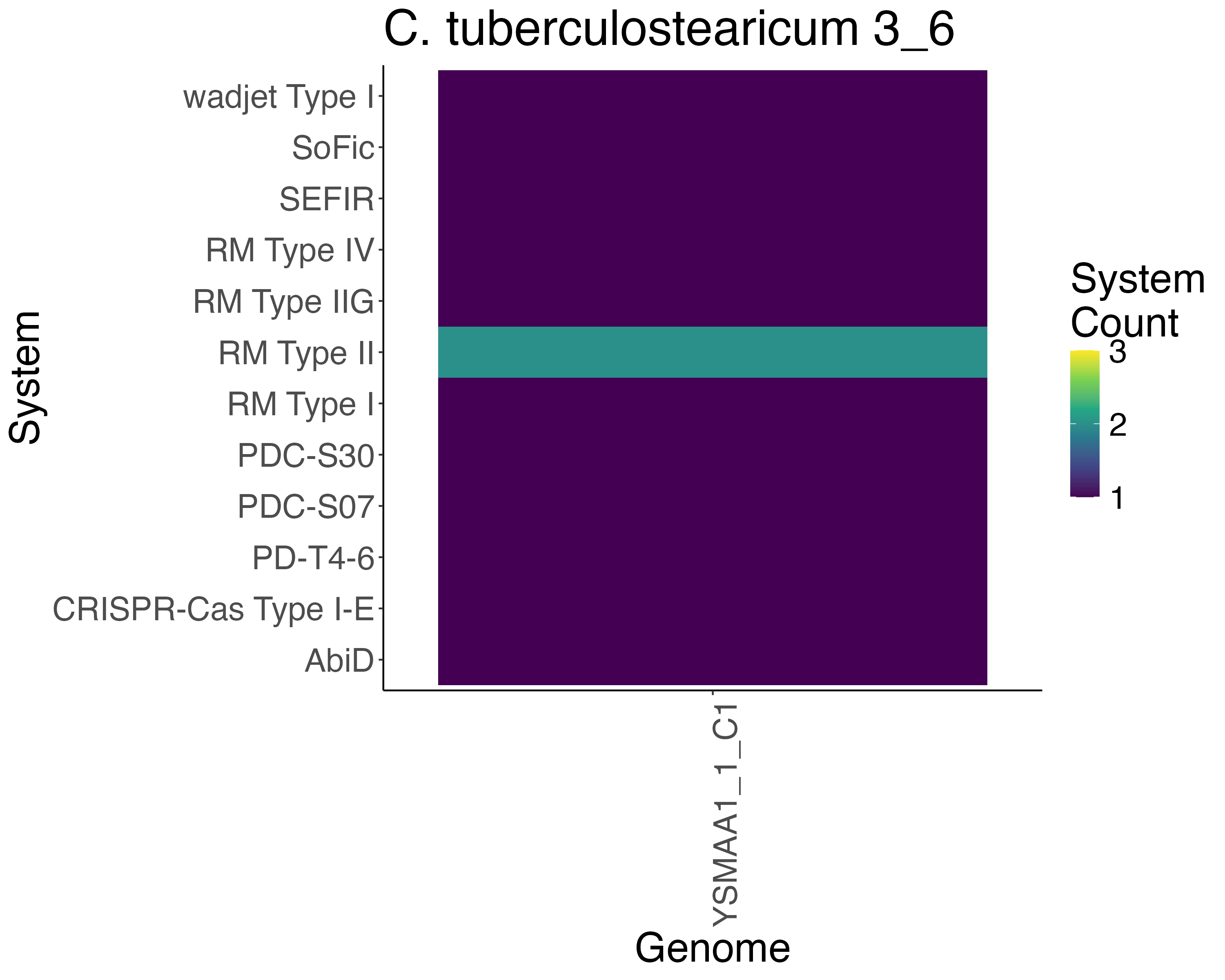

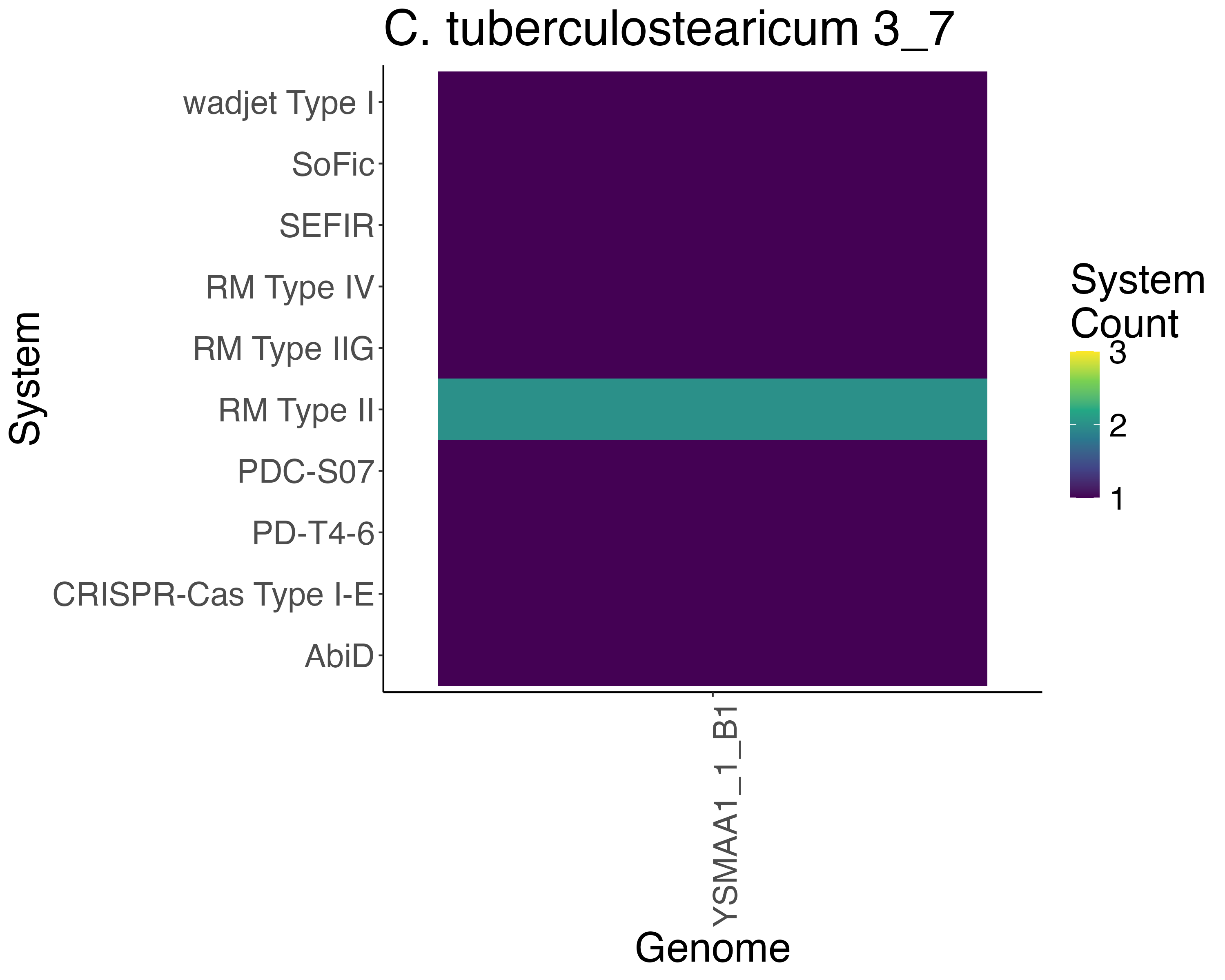

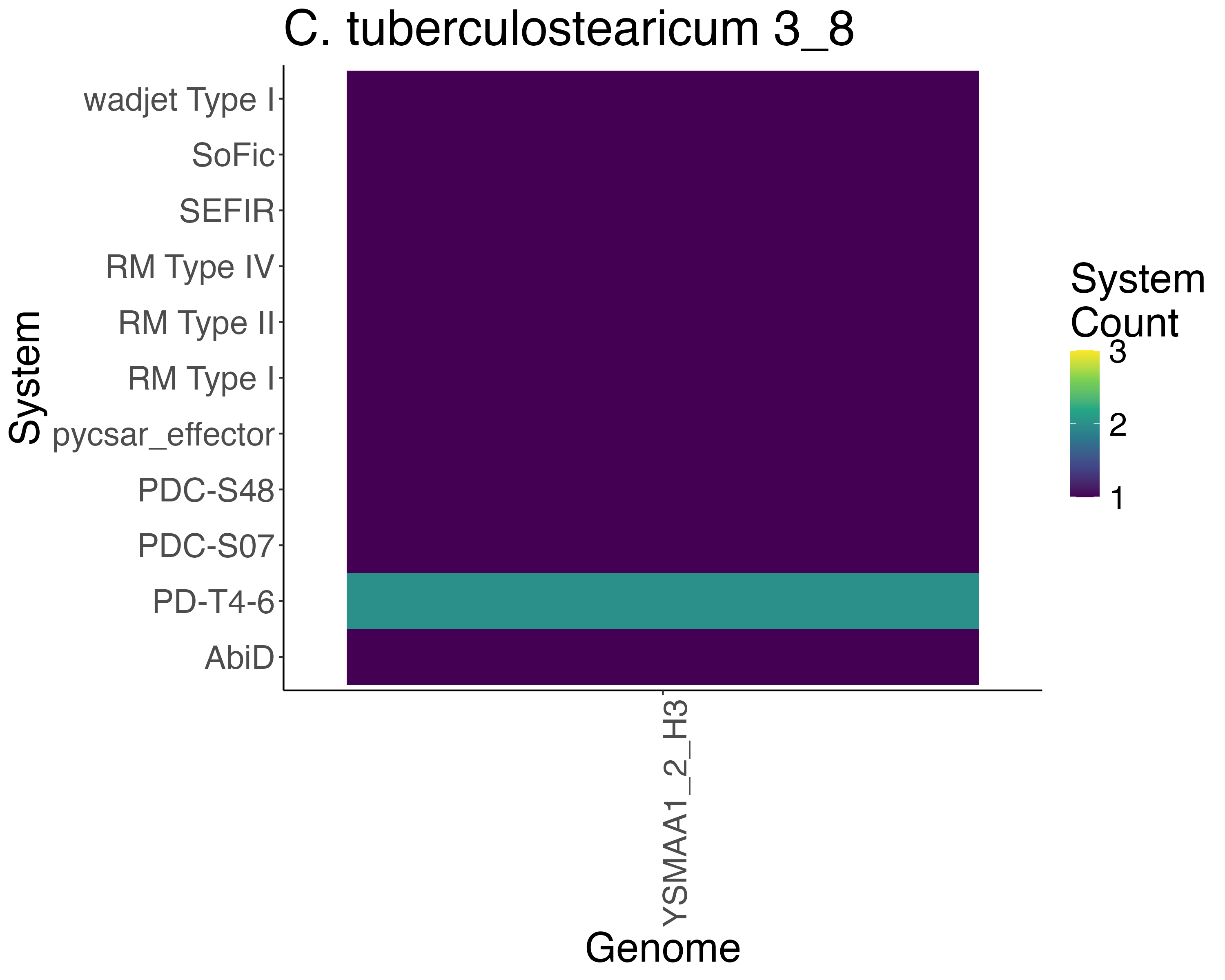

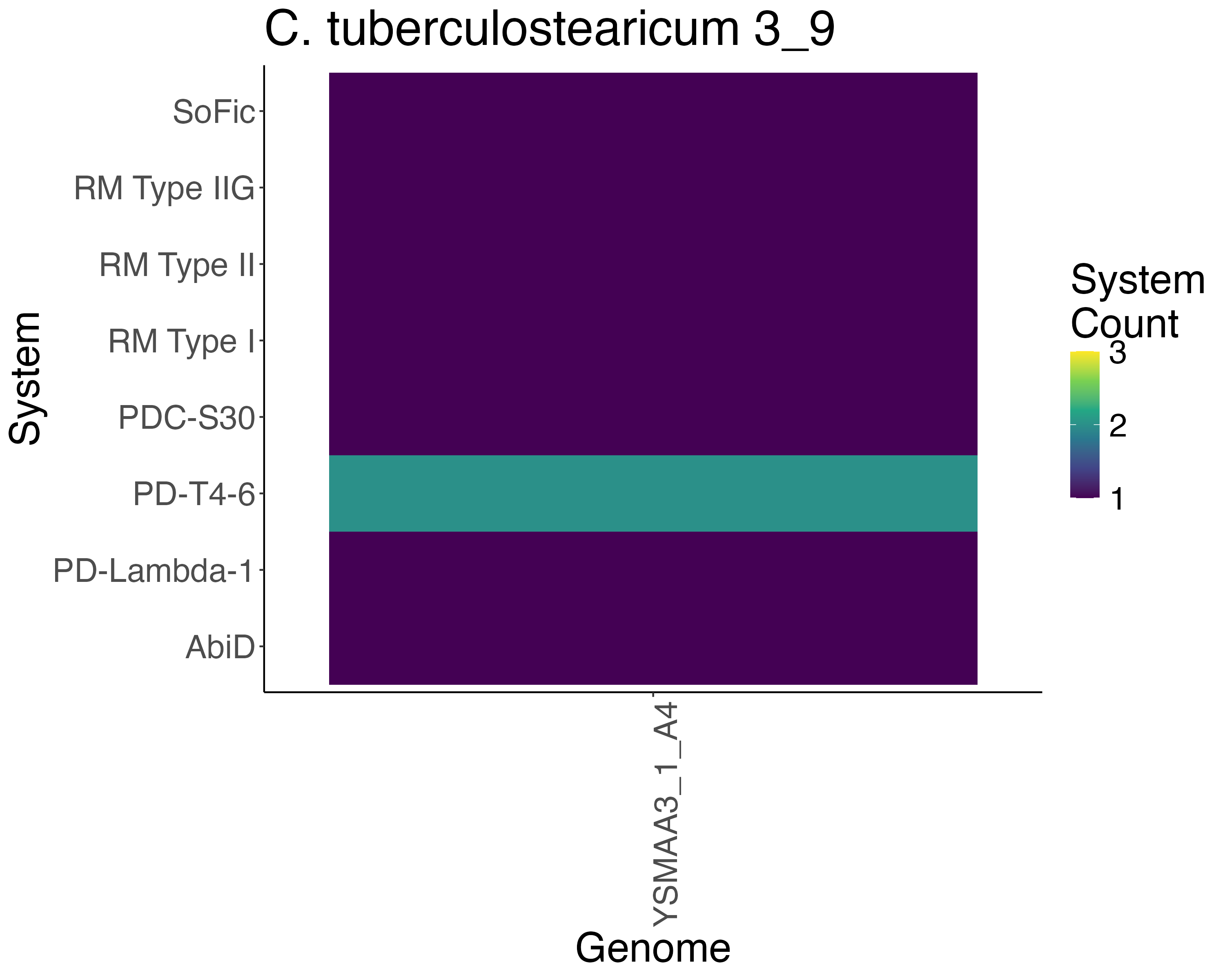

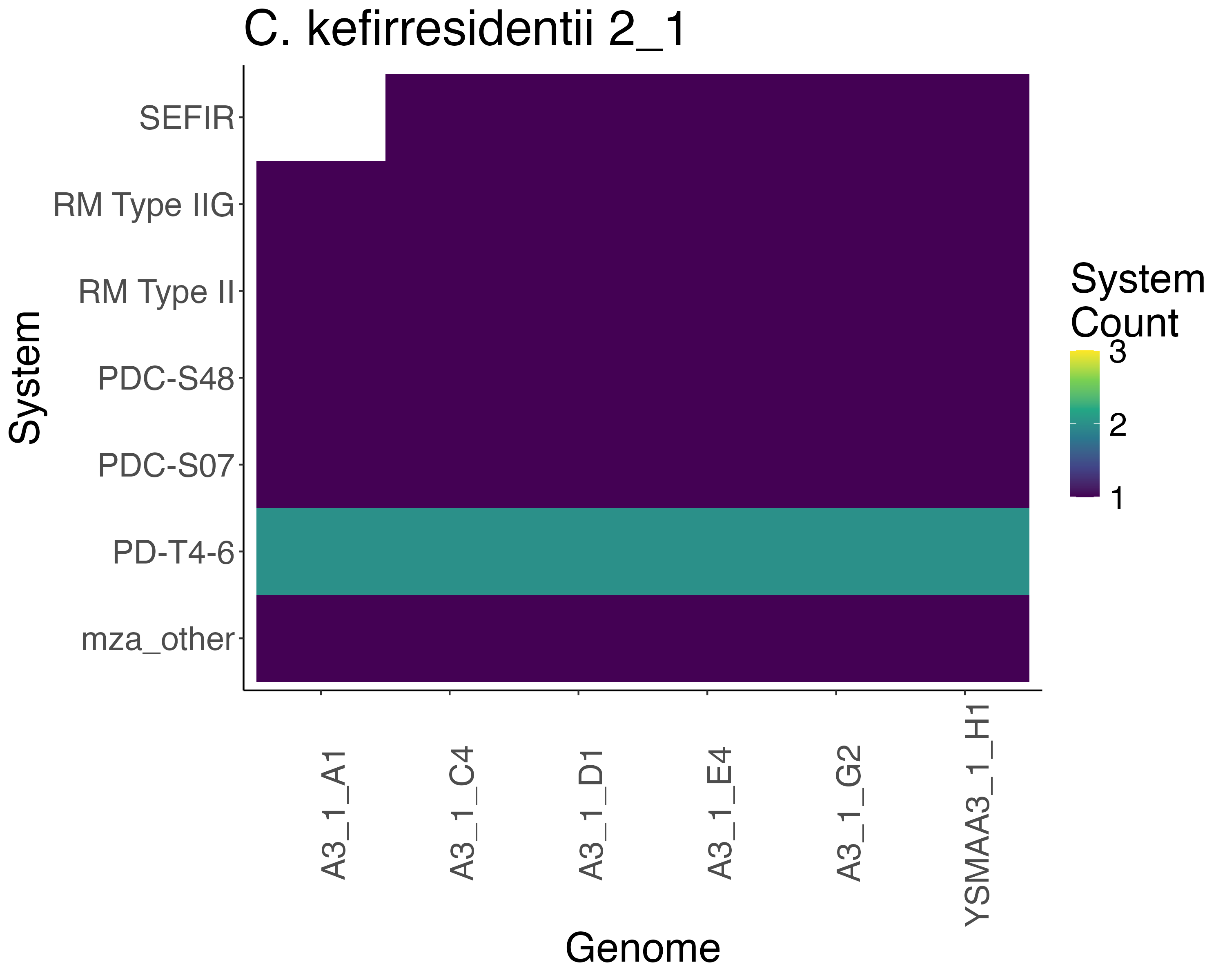

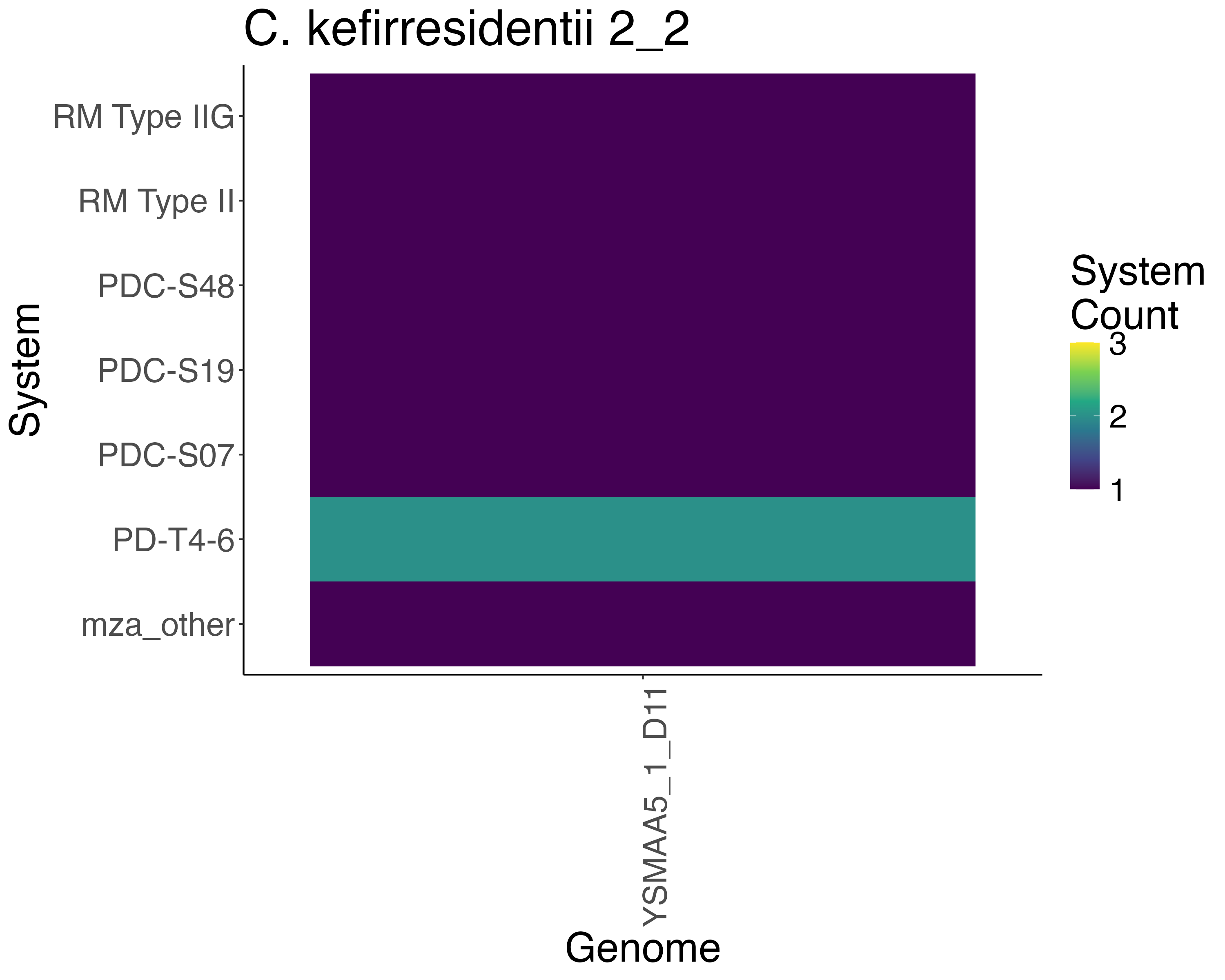

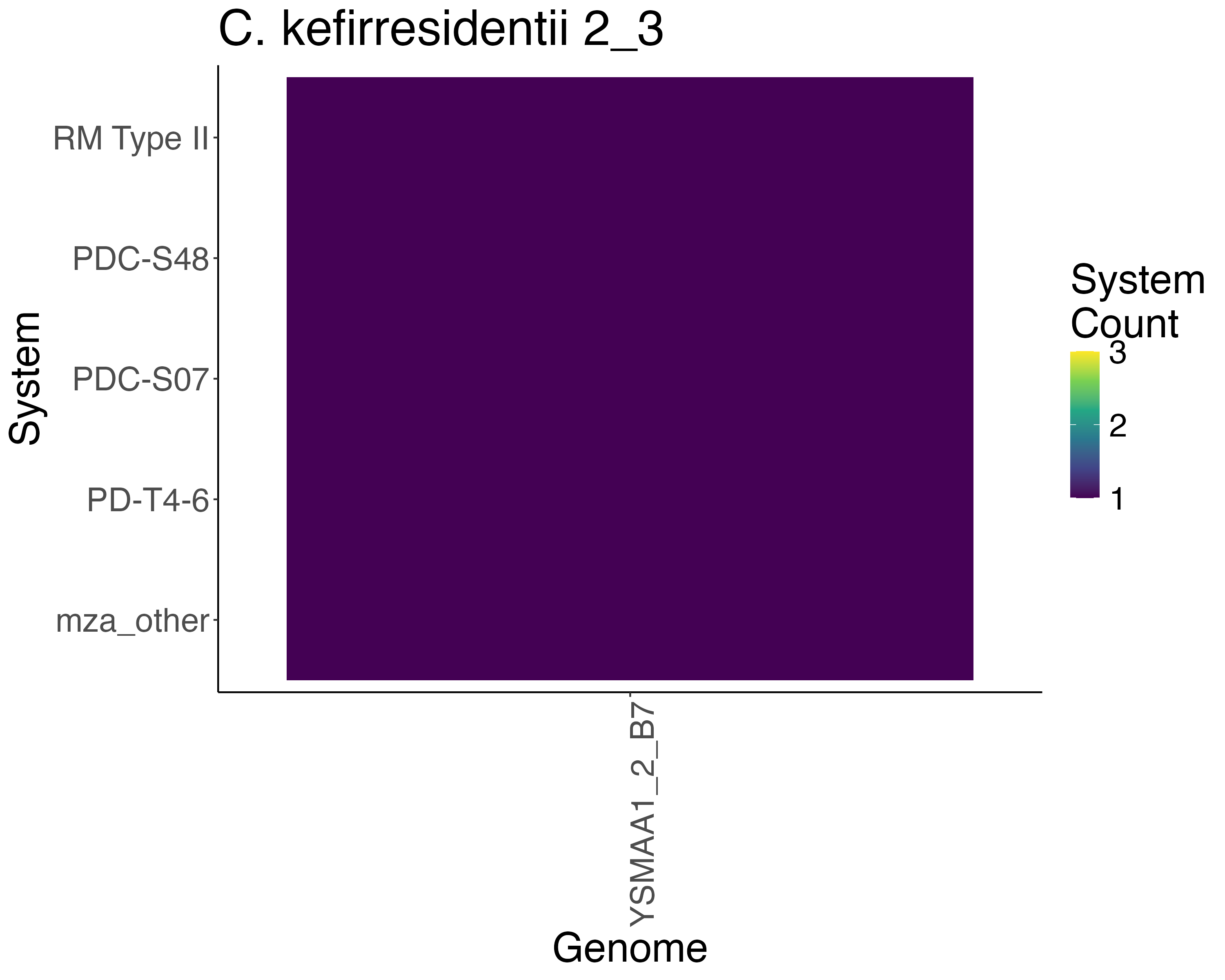

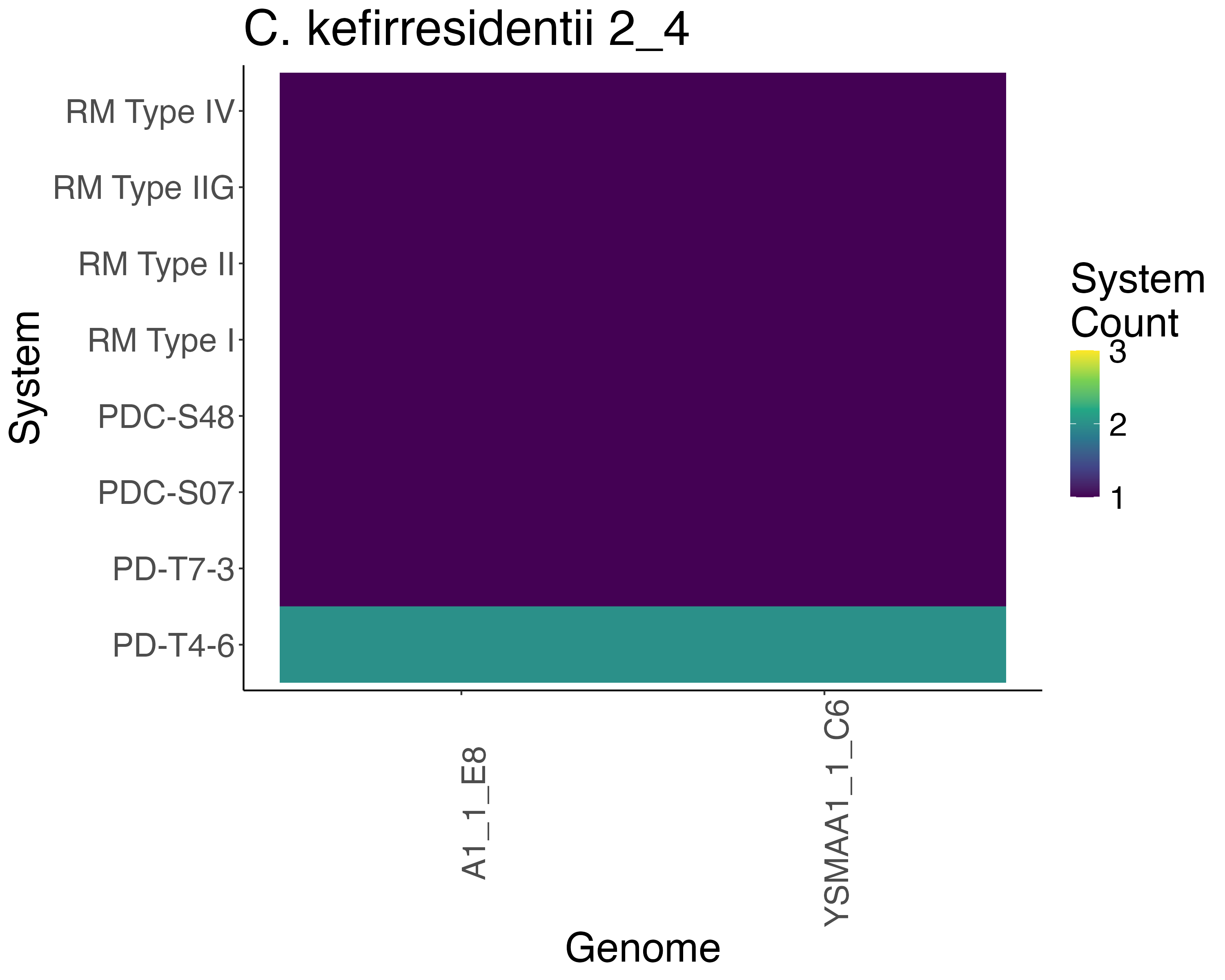

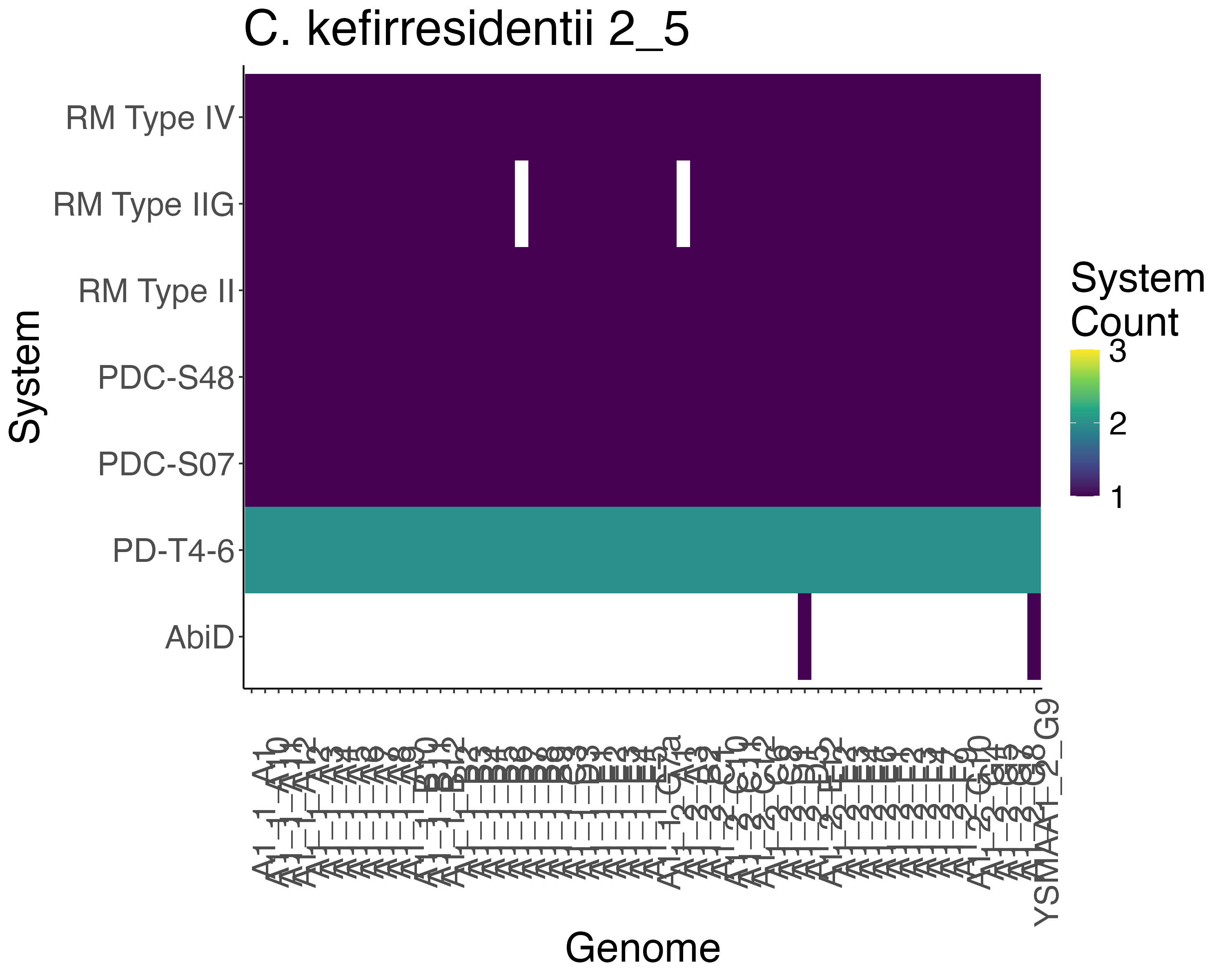

**Supplementary Figure 18. Phage defence systems found in drep secondary clusters of (A-I) *C. tuberculostearicum* and (J-N) *C. kefirresidentii****.* The heatmap describes the number of each identified system with (Yellow: 3, Green: 2, Purple: 1 and White: 0).

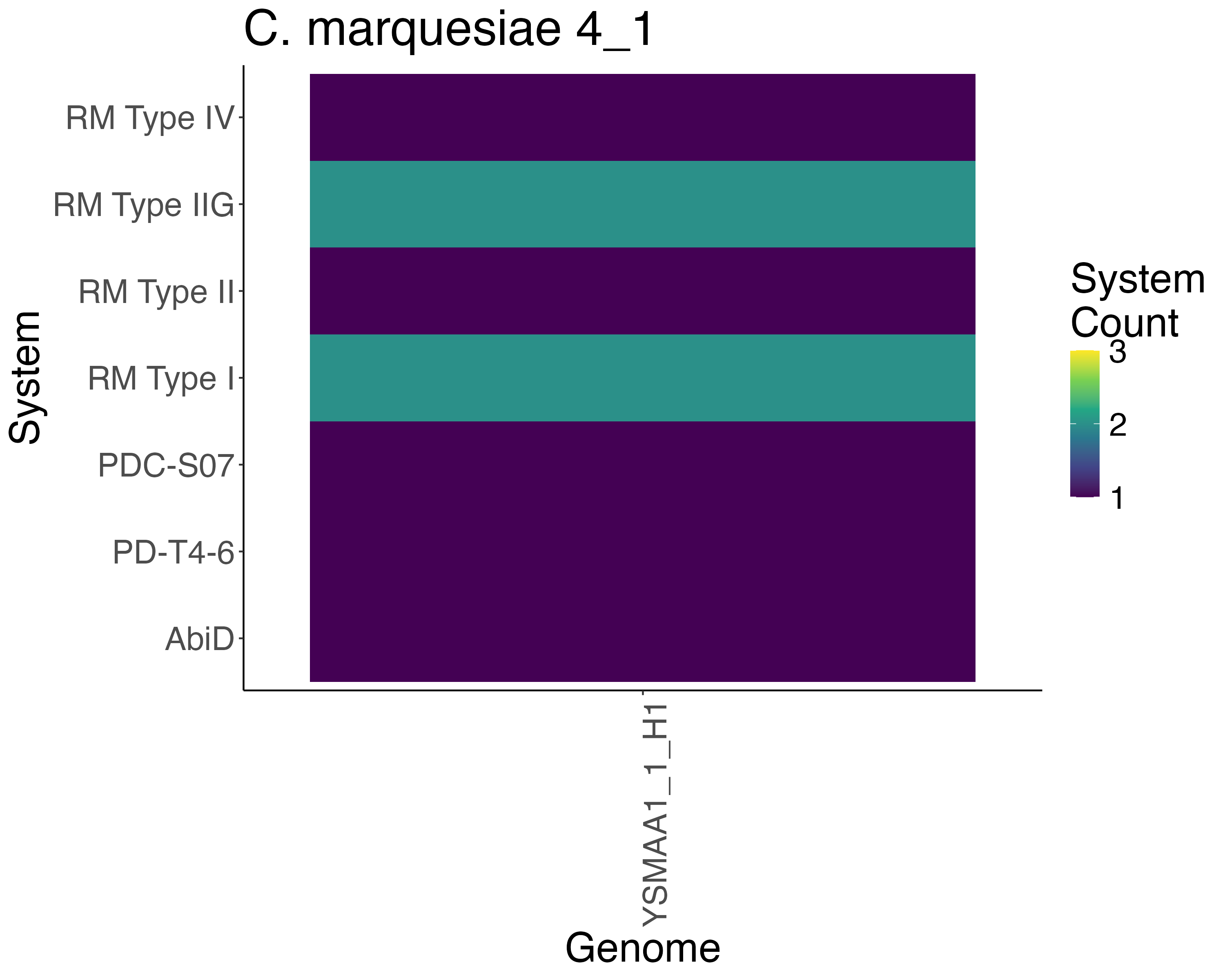

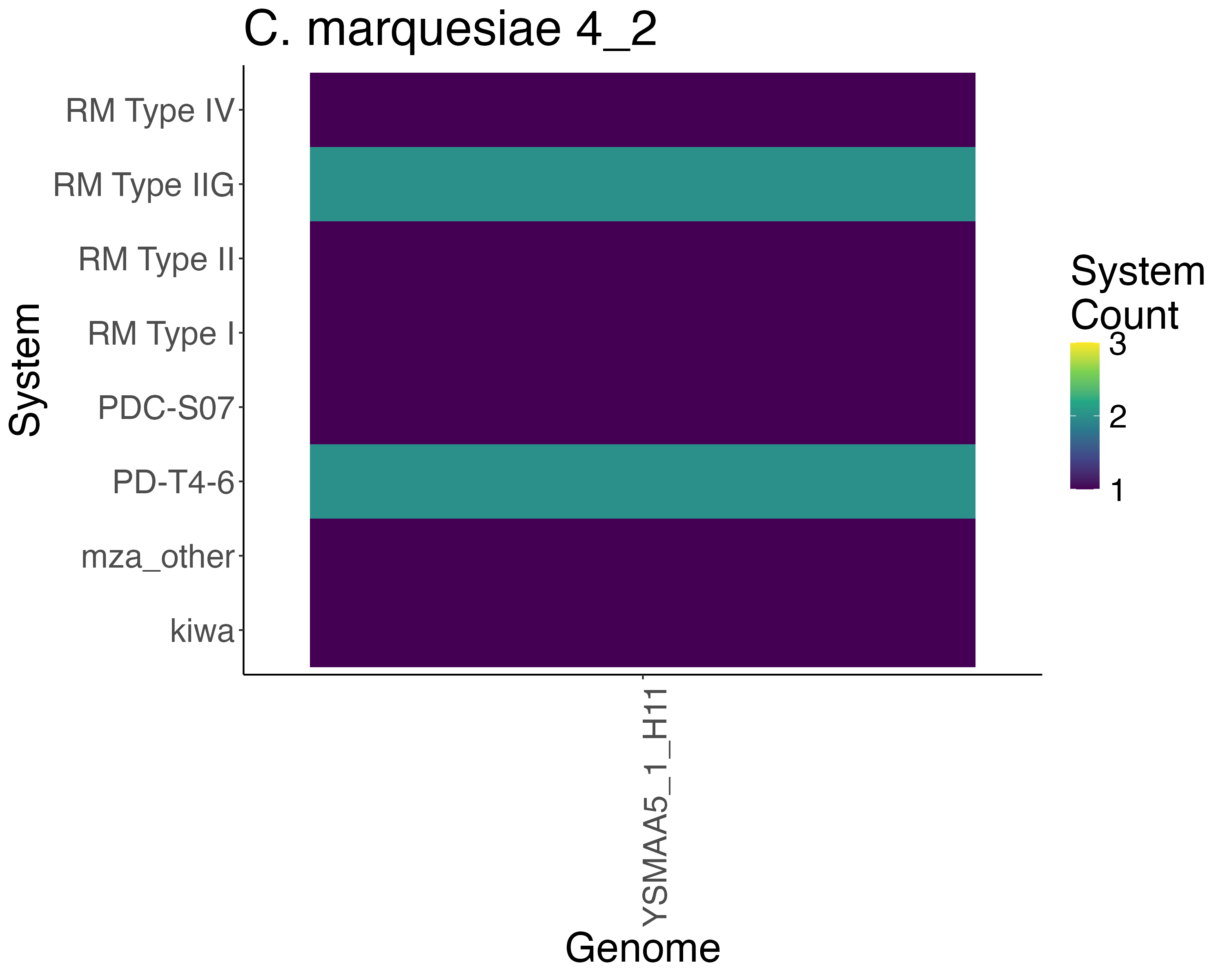

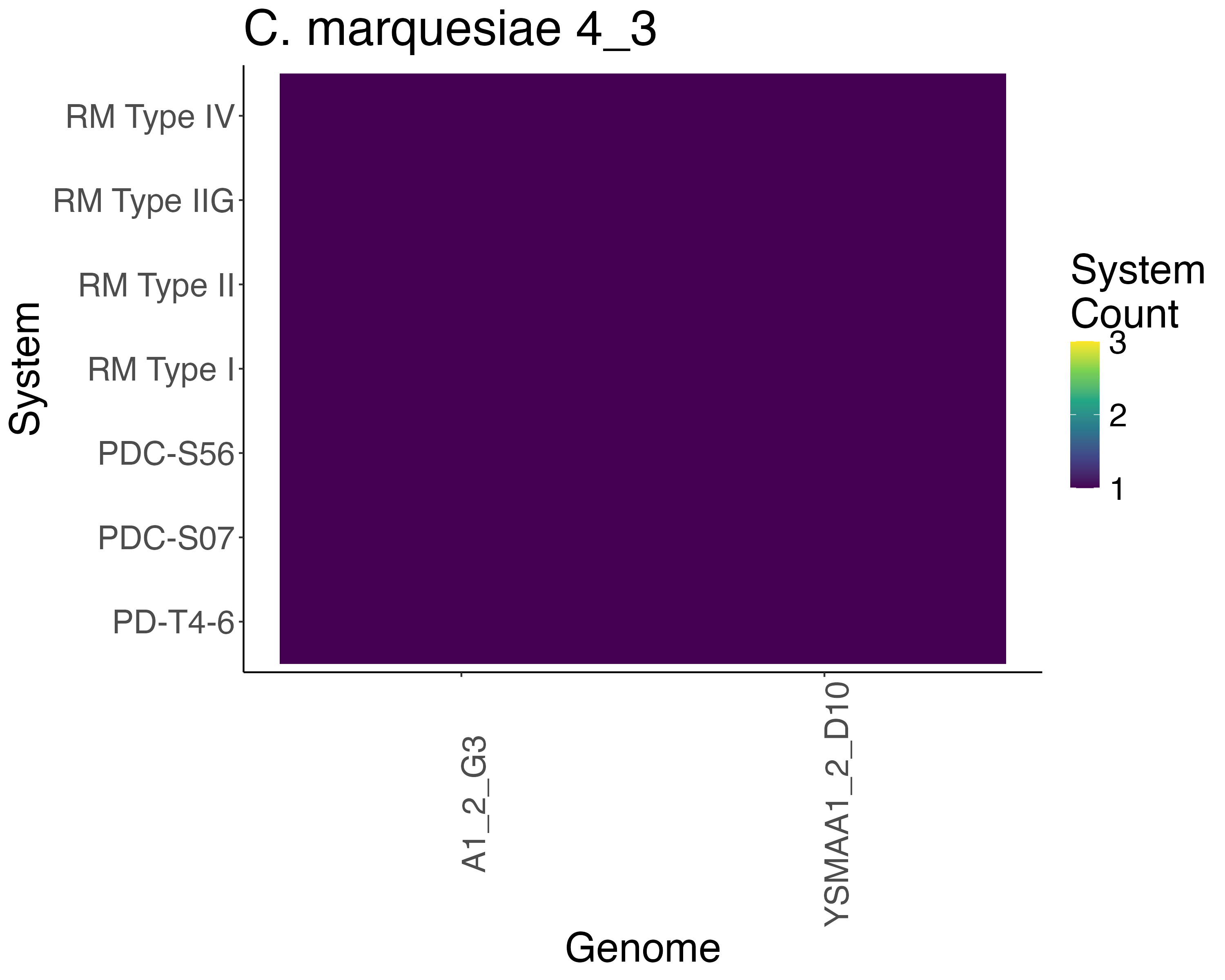

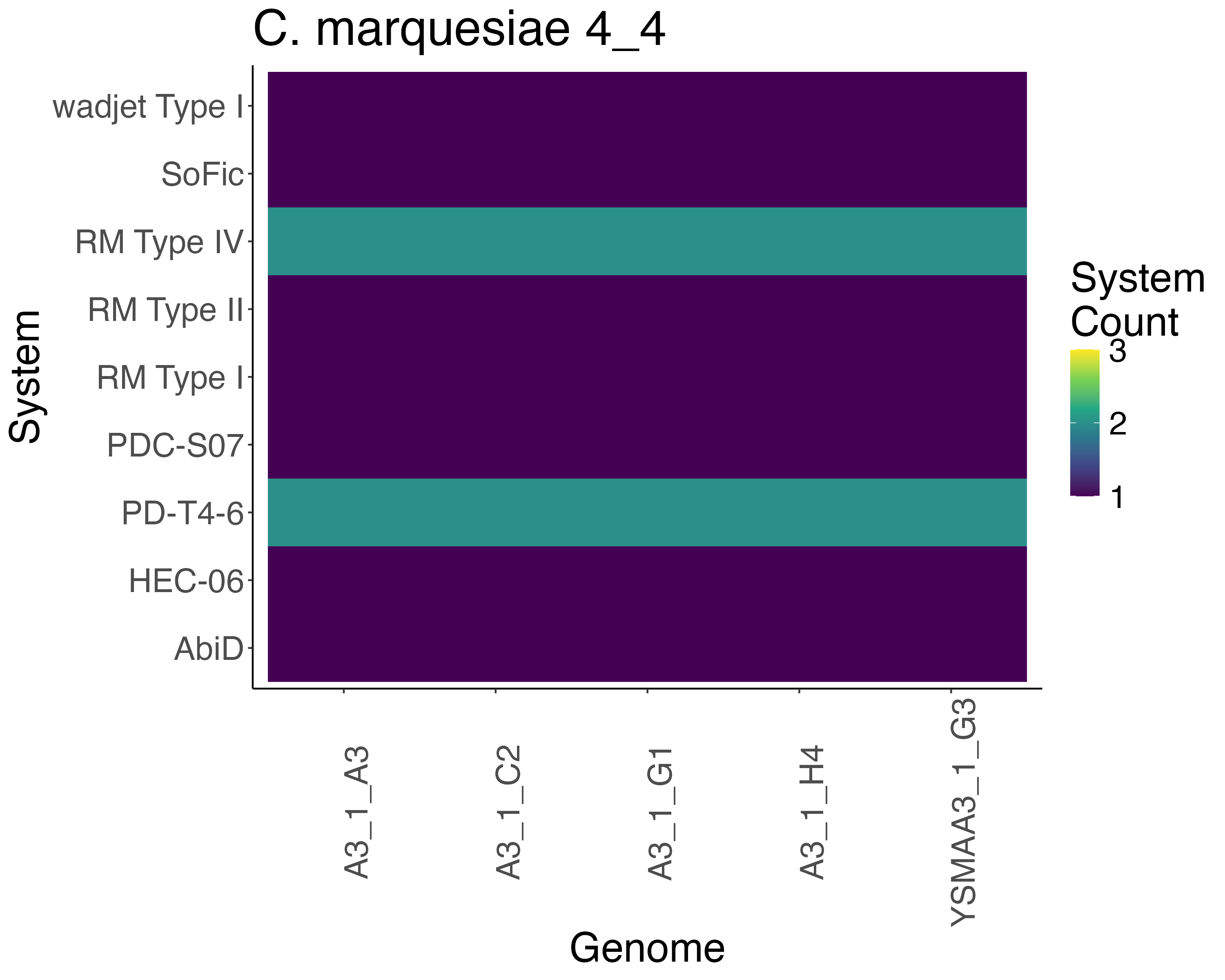

**Supplementary Figure 19. Phage defence systems found in drep secondary clusters of (A-J) *C. marquesiae****.* The heatmap describes the number of each identified system with (Yellow: 3, Green: 2, Purple: 1 and White: 0).

**Supplementary Figure 20.** The proportion of the total number of isolates derived from each of the 4 volunteers which bin to the 7 identified species (*C. tuberculostearicum*, *C. marquesiae*, *C. kefirresidentii*, *C. amycolatum*, *C. axilliensis*, *C. gottingense* and *C. jamesii*)
